# Widespread selection and gene flow shape the genomic landscape during a radiation of monkeyflowers

**DOI:** 10.1101/342352

**Authors:** Sean Stankowski, Madeline A. Chase, Allison M. Fuiten, Murillo F. Rodrigues, Peter L. Ralph, Matthew A. Streisfeld

## Abstract

Speciation genomic studies aim to interpret patterns of genome-wide variation in light of the processes that give rise to new species. However, interpreting the genomic ‘landscape’ of speciation is difficult, because many evolutionary processes can impact levels of variation. Facilitated by the first chromosome-level assembly for the group, we use whole-genome sequencing and simulations to shed light on the processes that have shaped the genomic landscape during a recent radiation of monkeyflowers. After inferring the phylogenetic relationships among the nine taxa in this radiation, we show that highly similar diversity (π) and differentiation (*F_ST_*) landscapes have emerged across the group. Variation in these landscapes was strongly predicted by the local density of functional elements and the recombination rate, suggesting that the landscapes have been shaped by widespread natural selection. Using the varying divergence times between pairs of taxa, we show that the correlations between *F_ST_* and genome features arose almost immediately after a population split and have become stronger over time. Simulations of genomic landscape evolution suggest that background selection (i.e., selection against deleterious mutations) alone is too subtle to generate the observed patterns, but scenarios that involve positive selection and genetic incompatibilities are plausible alternative explanations. Finally, tests for introgression among these taxa reveal widespread evidence of heterogeneous selection against gene flow during this radiation. Thus, combined with existing evidence for adaptation in this system, we conclude that the correlation in *F_ST_* among these taxa informs us about the genomic basis of adaptation and speciation in this system.

**Author summary:** What can patterns of genome-wide variation tell us about the speciation process? The answer to this question depends upon our ability to infer the evolutionary processes underlying these patterns. This, however, is difficult, because many processes can leave similar footprints, but some have nothing to do with speciation *per se*. For example, many studies have found highly heterogeneous levels of genetic differentiation when comparing the genomes of emerging species. These patterns are often referred to as differentiation ‘landscapes’ because they appear as a rugged topography of ‘peaks’ and ‘valleys’ as one scans across the genome. It has often been argued that selection against deleterious mutations, a process referred to as background selection, is primarily responsible for shaping differentiation landscapes early in speciation. If this hypothesis is correct, then it is unlikely that patterns of differentiation will reveal much about the genomic basis of speciation. However, using genome sequences from nine emerging species of monkeyflower coupled with simulations of genomic divergence, we show that it is unlikely that background selection is the primary architect of these landscapes. Rather, differentiation landscapes have probably been shaped by adaptation and gene flow, which are processes that are central to our understanding of speciation. Therefore, our work has important implications for our understanding of what patterns of differentiation can tell us about the genetic basis of adaptation and speciation.

## Introduction

The primary goal of speciation genomics is to interpret patterns of genome-wide variation in light of the ecological and evolutionary processes that contribute to the origin of new species (Ravinet et al. 2017, Wolf and Ellegren 2017, Campbell et al. 2018). Advances in DNA sequencing now allow us to capture patterns of genome-wide variation from organisms across the tree of life, but inferring the processes underlying these patterns remains a formidable challenge (Ravinet et al. 2017). This is because speciation is highly complex, involving a range of factors and processes that shape genomes through time and across different spatial and ecological settings (Abbott et al. 2013).

The difficulty of inferring process from pattern is illustrated by recent efforts to characterize the genomic basis of reproductive isolation using patterns of genome-wide variation (Ravinet et al. 2017, Wolf and Ellegren 2017, Campbell et al. 2018). Numerous studies have revealed highly heterogeneous differentiation ‘landscapes’ between pairs of taxa at different stages in the speciation process (Turner et al. 2005, Hohenlohe et al. 2010, Ellegren et al. 2012, Martin et al. 2013, Renaut et al. 2013, Poelstra et al. 2014, Soria-Carrasco et al. 2014, Lamichhaney et al. 2015, Malinsky et al. 2015). This pattern, which is characterized by peaks and valleys of relative differentiation (i.e., *F_ST_*) across the genome, was initially thought to provide insight into the genomic architecture of porous species boundaries (Wu 2001). Peaks in the differentiation landscape were interpreted as genomic regions containing loci underlying reproductive barriers, while valleys were thought to reflect regions that were homogenized by ongoing gene flow (Turner et al. 2005, Nosil et al. 2009, Feder et al. 2012). However, as the field of speciation genomics has matured, it has become clear that heterogeneous differentiation landscapes can be influenced by factors that have nothing to do with speciation *per se*.

For example, it is now clear that levels of genome-wide differentiation (*F_ST_*) can be influenced by the genomic distribution of intrinsic properties, including the recombination rate and the local density of functional sites (Cruickshank and Hahn 2014). This is because these properties affect the way that natural selection impacts levels of variation across the genome. First, regions enriched for functional sequence are more likely to be subject to selection, because they provide a larger target size for mutations with fitness effects. Second, when positive or negative directional selection acts on these mutations, it can indirectly reduce levels of genetic variation at statistically associated sites (Maynard-Smith and Haigh 1974, Charlesworth et al. 1993, Hudson and Kaplan 1995, Charlesworth 1998, Gillespie 2000, Coop and Ralph 2012, Cruickshank and Hahn 2014). Because these indirect effects of selection are mediated by linkage disequilibrium (LD) between selected and neutral sites, stronger reductions in diversity (and corresponding increases in *F_ST_*) are expected in genomic regions with low recombination, as this is where LD breaks down most gradually (Baird 2015). This is also why these indirect effects of selection are referred to as ‘linked selection’, even though physical linkage is not actually the cause. In this paper, we refer to the indirect effects of selection, as ‘indirect selection’ for short.

Although genomic features were not initially expected to play a major role in shaping patterns of between species variation (Kimura 1968, Ohta 1973) (but see (Charlesworth 1998, Gillespie 2000)), recent empirical studies indicate that they can have a strong impact on the topography of the differentiation landscape (Hahn 2008, Cruickshank and Hahn 2014, Burri 2017a, b, Wolf and Ellegren 2017). The most compelling evidence comes from studies that compare not one pair of taxa, but several closely related taxa that share a similar distribution of genomic elements due to their recent common ancestry (Burri 2017a, b, Wolf and Ellegren 2017). For example, Burri et al. (Burri et al. 2015) examined multiple episodes of divergence in *Ficedula* flycatchers and found strikingly similar differentiation landscapes between distinct pairs of taxa, presumably due to a shared pattern of indirect selection across the genome. Highly correlated differentiation landscapes have also been found in other groups of taxa, including sunflowers (Renaut et al. 2013), *Heliconius* butterflies (Kronforst et al. 2013, Martin et al. 2013), Darwin’s finches (Han et al. 2017), and other birds (Van Doren et al. 2017, Delmore et al. 2018).

But what, if anything, do these parallel signatures of selection reveal about the genomic basis of adaptation and speciation? Part of the answer lies in understanding which forms of selection cause these patterns. In a recent article, Burri (Burri 2017b) argued that selection against deleterious mutations (*i.e*., background selection, or BGS) is primarily responsible for the evolution of differentiation landscapes that are correlated both among taxa and with the distribution of intrinsic features. This argument was based on two premises: first, deleterious mutations are far more common than beneficial ones, so BGS has a greater opportunity to generate a genome-wide correlation between levels of variation and the distributions of intrinsic properties; second, although all functional elements are potential targets of BGS, the same loci are not expected to be repeatedly involved in adaptation or speciation across multiple taxa (Burri 2017b). Although this interpretation implies that correlated differentiation landscapes are unlikely to inform us about adaptation or speciation directly, it has been argued that this shared pattern of differentiation can be used to control for the effects of BGS when attempting to identify regions of the genome that have been affected by positive selection (Burri 2017b).

Even though it is often argued that correlated differentiation landscapes reflect the action of recurrent background selection, it stands to reason that they also could arise as a direct consequence of adaptation and speciation. For example, when adaptation occurs from standing variation, the rate of adaptation is not limited by the mutation rate, meaning that heterogeneous differentiation landscapes can evolve rapidly (Barrett et al. 2008, Bassham et al. 2018). Moreover, if adaptation is highly polygenic, positive selection will inevitably impact most of the genome (Rockman 2012). This could cause levels of differentiation to become correlated with the distribution of intrinsic genomic features, even across multiple taxa that are adapting to different environments. Also, correlated differentiation landscapes may arise across multiple closely related taxa due to a common basis of reproductive isolation among taxa. This is because incomplete isolating barriers generate a heterogeneous pattern of selection against gene flow across the genome. The loci underlying these porous barriers, including genetic incompatibilities and locally adapted alleles, are expected to accumulate in gene rich regions and will have stronger barrier effects in genomic regions where the recombination rate is low (Schumer et al. 2018, Martin et al. 2019). Thus, speciation also may result in genome-wide correlations between levels of differentiation and genome features, especially if reproductive isolation is highly polygenic.

Facilitated by a new chromosome-level genome assembly, genetic map, and annotation, we combine analyses of whole-genome sequencing with simulations to understand how different processes have contributed to the evolution of correlated genomic landscapes during a recent radiation. The bush monkeyflower radiation consists of eight taxa of *Mimulus aurantiacus* distributed mainly throughout California. (Fig. 1; (Chase et al. 2017). Together with their sister species *M. clevelandii,* they span a range of divergence times over the past approximately one million years. The plants inhabit a range of environments, including temperate coastal regions, mountain ranges, semi-arid habitats, and offshore islands (Thompson 2005). Crossing experiments have shown that all of these taxa are at least partially inter-fertile (McMinn 1951), and many hybridize in nature in narrow regions where their distributions overlap (Streisfeld and Kohn 2005, 2007, Sobel and Streisfeld 2015, Stankowski et al. 2015, 2017). Despite their close evolutionary relationships and opportunities for gene flow, these taxa show striking phenotypic differentiation (Chase et al. 2017). The most conspicuous trait differences are associated with their flowers, which show heritable variation in color, size, shape, and the placement of the reproductive organs (Streisfeld and Kohn 2005, Stankowski et al. 2015). In his seminal works on plant speciation, Grant (Grant 1981, 1993b, a) postulated that these floral trait differences were due to pollinator-mediated selection by different avian and insect pollinators. Detailed studies in one pair of taxa support this hypothesis and have shown that pollinator-mediated selection can generate strong premating reproductive isolation in the face of extensive gene flow (Streisfeld and Kohn 2005, 2007, Sobel and Streisfeld 2015, Stankowski et al. 2015, 2017).

**Figure 1.**
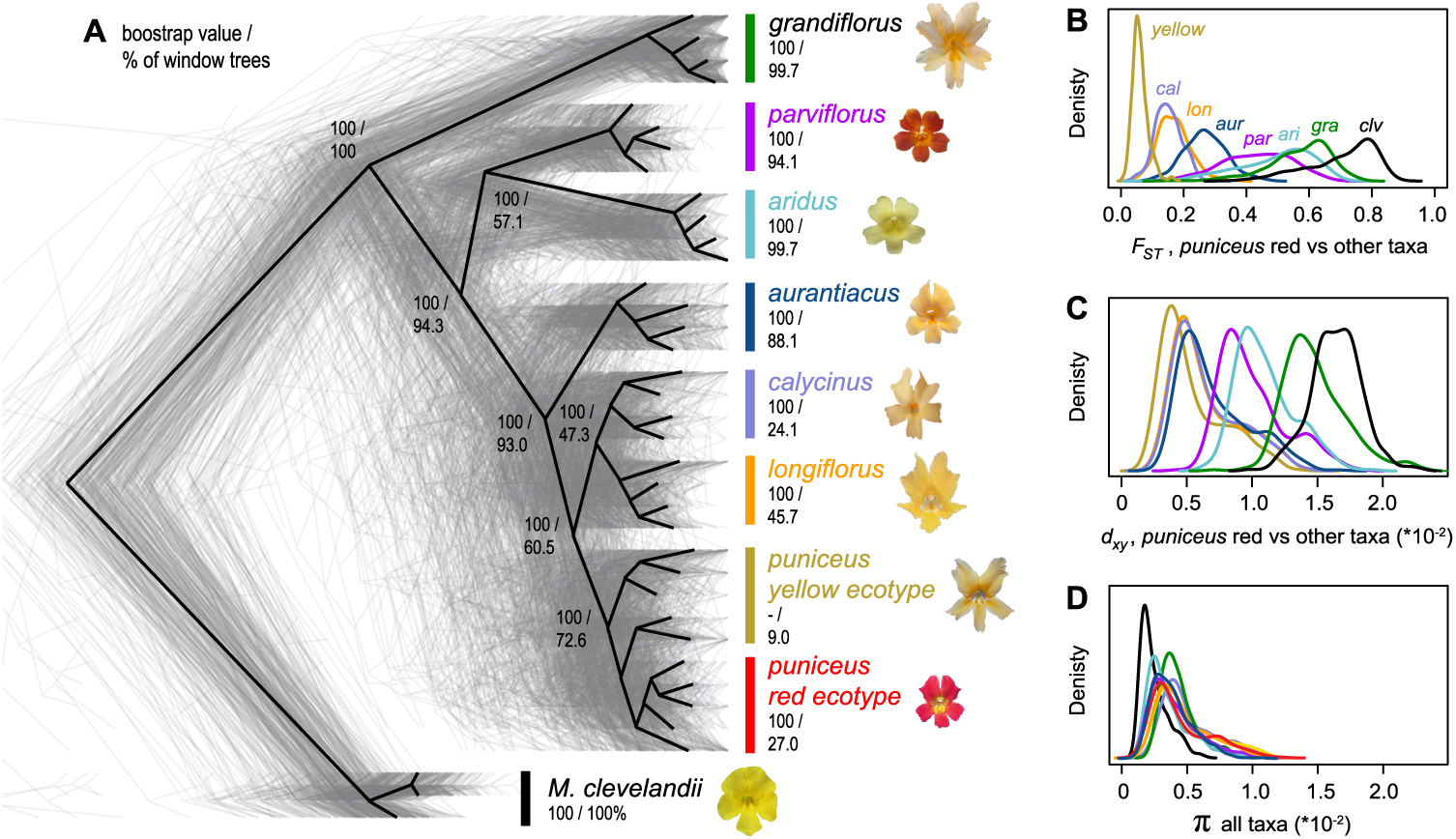
Evolutionary relationships and patterns of genome-wide variation across the radiation. A) The black tree was constructed from a concatenated alignment of genome-wide SNPs and is rooted using *M. clevelandii*. The 387 gray trees were constructed from 500 kb genomic windows. The first number associated with each node or taxon is the bootstrap support for that clade in the whole genome tree, and the second number is the percentage of window-based trees in which that clade is present. B) Levels of differentiation (*F_ST_*), C) divergence (*d*_xy_), and D) diversity (π) within and among taxa based on the same 500 kb windows. For simplicity, *F*_ST_ and *d*_xy_ are shown only for comparisons with the red ecotype of subspecies *puniceus*. See Fig. S7 for the distributions of *F*_ST_ and *d*_xy_ across all pairs of taxa.

After inferring the phylogenetic relationships among these taxa, we show that highly similar diversity (π) and differentiation (*F_ST_*) landscapes have emerged across the group. Variation in these landscapes was strongly predicted by the local density of functional elements and the recombination rate, suggesting that they have been shaped by widespread selection. Using the varying divergence times between pairs of taxa, we show that the correlations between *F_ST_* and genome features arose almost immediately after a population split and have become stronger over the course of time. Simulations of genomic landscape evolution suggest that background selection (i.e., selection against deleterious mutations) alone is too subtle to generate the observed patterns, but scenarios that involve positive selection and genetic incompatibilities are plausible alternative explanations. Finally, tests for introgression among these taxa reveal widespread evidence of heterogeneous selection against gene flow during this radiation. We discuss the implications of these results for our general understanding of genomic landscape evolution, particularly in light of recent efforts to reveal the genomic basis of adaptation and speciation.

## Results and Discussion

### A chromosome-level genome assembly, map, and annotation for the bush monkeyflower

To facilitate the analysis of genome-wide variation in this group, we constructed the first chromosome-level reference genome for the bush monkeyflower using a combination of long-read Single Molecule Real Time (SMRT) sequencing (PacBio), overlapping and mate-pair short-reads (Illumina), and a high-density genetic map (7,589 segregating markers across 10 linkage groups; Fig. S1; Table S1). Contig building and scaffolding yielded 1,547 scaffolds, with an N50 size of 1.6 Mbp, and a total length of 207 Mbp. The high-density map allowed us to anchor and orient 94% of the assembled genome onto 10 linkage groups, which is the number of chromosomes inferred from karyotypic analyses in all subspecies of *M. aurantiacus* and *M. clevelandii* (Vickery 1995). Analysis of assembly completeness based on conserved gene space (Simao et al. 2015) revealed that 93% of 1440 universal single copy orthologous genes were completely assembled, with a further 2% partially assembled (Table S2). Subsequent annotation yielded 23,018 predicted genes.

### Phylogenetic relationships among taxa and patterns of discordance across the genome

To infer phylogenetic relationships among the taxa in this radiation, we sequenced 37 whole genomes from the seven subspecies and two ecotypes of *Mimulus aurantiacus* (*n* = 4-5 per taxon) and its sister taxon *M. clevelandii* (*n =* 3) (Fig. S2; Table S3). Close sequence similarity allowed us to align reads from all samples to the reference assembly with high confidence (average 91.7% reads aligned; Table S3). After mapping, we identified 13.2 million variable sites that were used in subsequent analyses (average sequencing depth of 21x per individual, Table S3). Relationships were then inferred among the nine taxa from a concatenated alignment of genome-wide SNPs using maximum-likelihood (ML) phylogenetic analysis in *RAxML* (Stamatakis 2014).

The tree topology obtained from this analysis (Fig. 1) confirmed the same phylogenetic relationships as previous analyses based on reduced-representation sequencing and five different methods of phylogenetic reconstruction (Stankowski and Streisfeld 2015, Chase et al. 2017), and was supported by patterns of clustering from principal components analysis (Fig. S3). Individuals of each of the seven subspecies formed monophyletic groups with 100% bootstrap support (Fig. 1). Relationships within subspecies *puniceus* were more complex, as the red ecotype formed a monophyletic sub-clade within the paraphyletic yellow ecotype. This is consistent with the recent origin of red flowers from a yellow-flowered ancestor (Stankowski and Streisfeld 2015).

Although the whole genome phylogeny provides a well-supported summary of the relationships among these taxa, concatenated phylogenies can obscure phylogenetic discordance in more defined genomic regions (Pease et al. 2016) (Fig. 1A). To test for fine-scale phylogenetic discordance, we next constructed ML phylogenies for 500 kb and 100 kb genomic windows. We then calculated a ‘concordance score’ for each tree by computing the correlation between the distance matrix generated from each window-based tree and the whole-genome tree, with a stronger correlation indicating that two trees have a more similar topology.

At the 500 kb scale, only 22 (6%) trees showed the same taxon branching order as the whole-genome tree. However, concordance scores tended to be very high for all of the trees (mean = 0.964, s.d. 0.039; min 0.719), suggesting that variation in the topologies was due primarily to minor differences in branching order. This was confirmed by quantifying how often each node in the genome-wide tree was recovered in the set of window-based trees (Fig. 1). Specifically, we found that differences in branching order were associated with the most recent splits, which included pairs of closely related taxa. For example, even though subspecies *puniceus* was monophyletic in the majority of trees (72.6%), individuals from the red ecotype only formed a monophyletic group in 27% of the trees. Similarly, the closely related subspecies *longiflorus* and *calycinus* were monophyletic in only 45.7% and 24.1% of trees, respectively. Higher discordance among these closely related, geographically proximate taxa is likely due to two factors. First, only a short time has passed since they shared a common ancestor, meaning that ancestral polymorphisms have had little time to sort among lineages. Second, ongoing gene flow between some taxa may have opposed sorting, prolonging the retention of ancestral variants among them.

Next, we examined how the level of phylogenetic discordance varied across the bush monkeyflower genome. If the discordance was due to the stochastic effects of neutral processes, variation in tree concordance scores should be distributed randomly across the genome (Hudson 1990). To test this prediction, we plotted the tree concordance scores across the 10 linkage groups (Fig. 2A; Fig. S4 for results from 100 kb windows and Fig. S5 for plots along each chromosome). Rather than being randomly distributed, trees with lower concordance scores tended to cluster in relatively narrow regions of all 10 chromosomes (Fig. 2A; autocorrelation analysis permutation tests *p* = 0.001 - 0.023; Fig. S6). This non-random pattern indicates that the rate of sorting varies along chromosomes, which could be due to variation in the strength of indirect selection across the bush monkeyflower genome.

**Figure 2.**
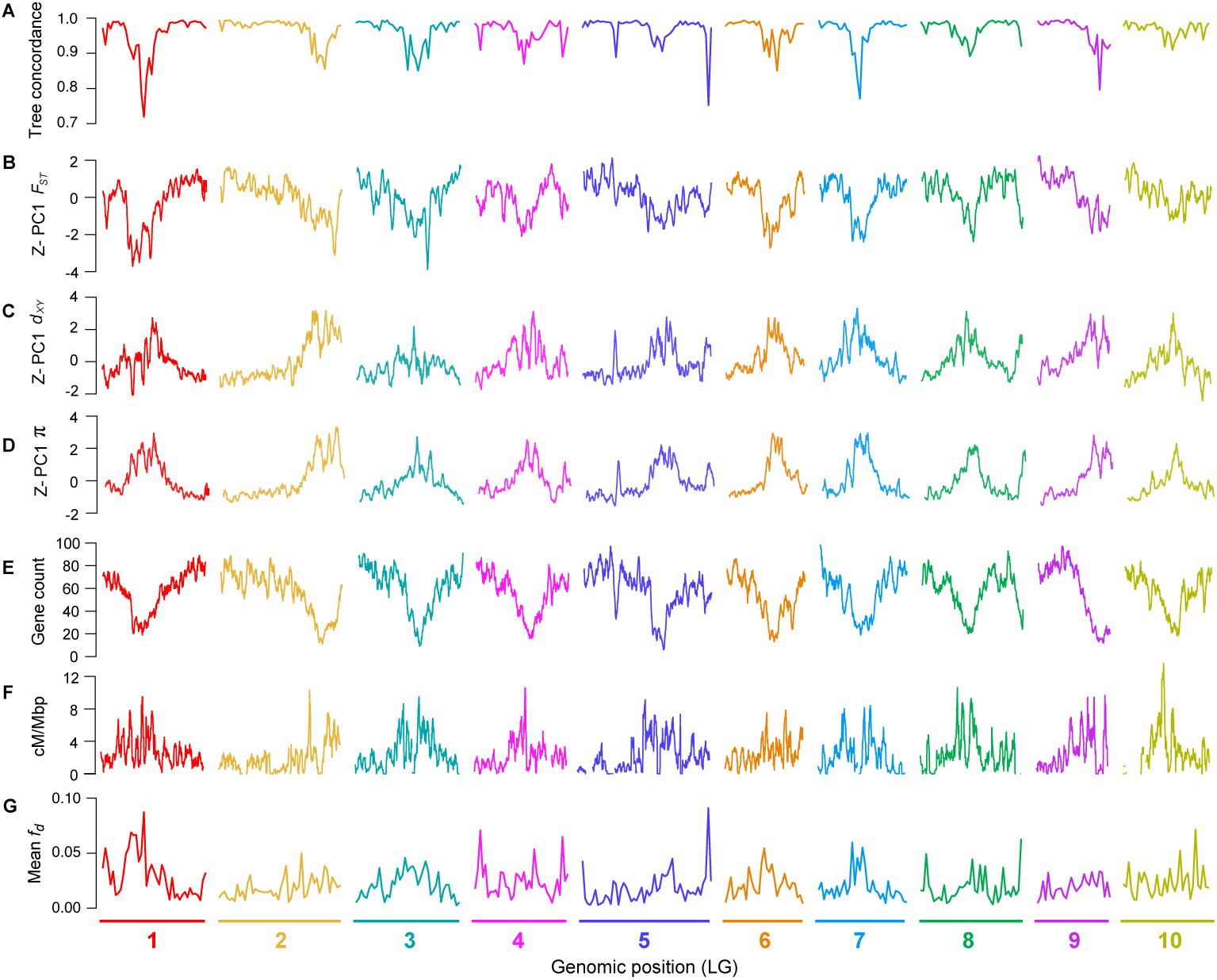
Common genomic landscapes mirror variation in the local properties of the genome. A) Tree concordance scores for 500 kb non-overlapping genomic windows plotted across the 10 bush monkeyflower chromosomes. B – D) Plots of the first principal component (PC1) for *F_ST_*, *d*_xy_, and π in overlapping 500 kb windows (step size = 50 kb). PC1 explains 66%, 70%, and 85% of the variation in *F_ST_*, *d*_xy_ and π, respectively, and is Z*-*transformed such that above average values have positive values and below average values have negative values. E – F) Gene count and recombination rate (cM/Mbp) in overlapping 500 kb windows. G), Mean *f_d_*(admixture proportion) in 500 kb non-overlapping genomic windows. See Fig. S4 for the same plot made for 100 kb windows.

### Correlated patterns of genome-wide variation across the radiation

To gain deeper insight into the evolutionary processes that have shaped patterns of genome-wide variation during this radiation, we used three summary statistics to quantify patterns of diversity (π), divergence (*d_xy_*), and differentiation (*F_ST_*) within and between these taxa (the same 500 kb and 100 kb sliding windows as above). π, or nucleotide diversity, was used to quantify the level of genetic variation within each taxon. It is defined as the average number of nucleotide differences per site between two sequences drawn from the population (Nei and Li 1979). Second, we calculated *d*_xy_ as a measure of sequence divergence between all 36 pairs of taxa. In principle, π and *d*_xy_ are the same measure, except that the former is calculated between sequences within a taxon and the latter between pairs of taxa; they have the same units and both are proportional to the average coalescence time of the sequences being compared multiplied by mutation rate. Third, we calculated *F_ST_*between each pair of taxa. Unlike *d*_xy_, which is a measure of divergence between sequences drawn from different taxa, *F*_ST_ measures differentiation between the taxa relative to the total diversity in the sample. Between samples *x* and *y*, *F*_ST_ is roughly equal to 1 - π/*d*_xy_, where π is the mean diversity in the two taxa (Slatkin 1991). *F*_ST_ is strongly influenced by current levels of within-taxon diversity, while *d*_xy_ is strongly influenced by the level of ancestral variation when divergence times are small. The use of the term ‘divergence’ to describe both *d*_xy_ and *F*_ST_ has caused some confusion in the literature, leading to alternative naming schemes. Here, we follow a recently proposed convention, referring to *d*_xy_ as ‘divergence’ and *F*_ST_ as ‘differentiation’ (Ravinet et al. 2017).

The variation in *F_ST_* among all 36 pairs of taxa highlights the continuous nature of differentiation across the group (Fig. 1B; Fig. S7), with mean window-based estimates ranging from 0.06 (red vs. yellow ecotypes of *puniceus*) to more than 0.70. Distributions of divergence (*d*_xy_) show a similar pattern (Fig. 1C), with mean values ranging from 0.54% (red vs. yellow ecotypes) to 1.6% (yellow ecotype vs. *M. clevelandii*). Broad distributions of window-based estimates indicate high variability in levels of differentiation and divergence among genomic regions (Fig. 1B & 1C). Window-based estimates of nucleotide diversity also varied markedly (π; Fig. 1D), ranging from 0.09% to 1.26%, even though mean estimates were very similar among the ingroup taxa (0.37% to 0.53%) and were only slightly lower in *M. clevelandii* (0.26%).

As with tree concordance, variation in these summary statistics was non-randomly distributed across broad regions of each chromosome (*p* < 0.005; Fig. 2; Fig. S4; Fig. S5; Fig. S6). To account for the magnitude of variation in these statistics across all nine taxa (for π) or among the 36 pairs of taxa (for *d_xy_* and *F_ST_*), we normalized the window-based estimates using *Z*-transformation and plotted them across the genome (Fig. S5). Visual inspection of these data revealed that patterns of genome-wide variation in each statistic were qualitatively similar in all comparisons. We therefore used principal components analysis to quantify their similarity and extracted a single variable (PC1) that summarized the common pattern (Fig. S5).

These analyses confirmed that patterns of genome-wide variation were highly correlated across this group of taxa. Indeed, PC1 explained 65.9% of the variation in *F_ST_* across the 36 pairwise comparisons. Further, all comparisons loaded positively onto PC1 (mean loading = 0.78 s.d. 0.18; Table S5 for all loadings), indicating that peaks and troughs of *F_ST_* tended to occur in the same genomic regions across all comparisons. Patterns of genome-wide divergence (*d_xy_*) and diversity (π) also were highly correlated across comparisons, with PC1 explaining 69.5% and 84.7% of the variation among the window-based estimates, respectively. Again, all taxa (for π) and taxon comparisons (for *d_xy_*) loaded positively onto the first principal component (mean loading for *d_xy_*= 0.78 s.d. 0.18; for π, 0.91 s.d. 0.07). PC1 therefore provides a summary of the original landscapes and is effectively the same as taking the mean window-based scores for each statistic (*r*^2^ between PC1 and mean scores > 0.995 for all three statistics).

### Common genomic landscapes have been shaped by heterogeneous indirect selection

Observing highly similar genomic landscapes suggests that a common pattern of heterogeneous selection has shaped variation in all nine taxa. Indeed, if a region experiences recurrent indirect selection across the phylogenetic tree, then it should show lower diversity (π) within species and lower divergence (*d_xy_*) between species, because *d_xy_*is influenced by levels of diversity in the common ancestor (Charlesworth et al. 1993, Charlesworth 1998, Burri 2017b). In agreement with this prediction, we observed a strong positive correlation between PC1 *d*_xy_ and PC1 π (*r =* 0.84), indicating that regions of the genome with lower diversity tended to be less diverged between these taxa (and thus had lower ancestral diversity) (Fig. 3, Fig. S8 for scatterplots and Fig. S9 for results at 100 kb scale). Regions with reduced diversity also tended to show higher levels of differentiation (*F_ST_*) (*r* = −0.84) and tree concordance (*r* = −0.69). These relationships are also predicted by models of recurrent indirect selection, because local reductions in diversity decrease the amount of genetic variation in future generations, similar to a local reduction in *N_e_*. As a result, ancestral variants in impacted regions sort more rapidly than under neutrality (Pease and Hahn 2013).

**Figure 3.**
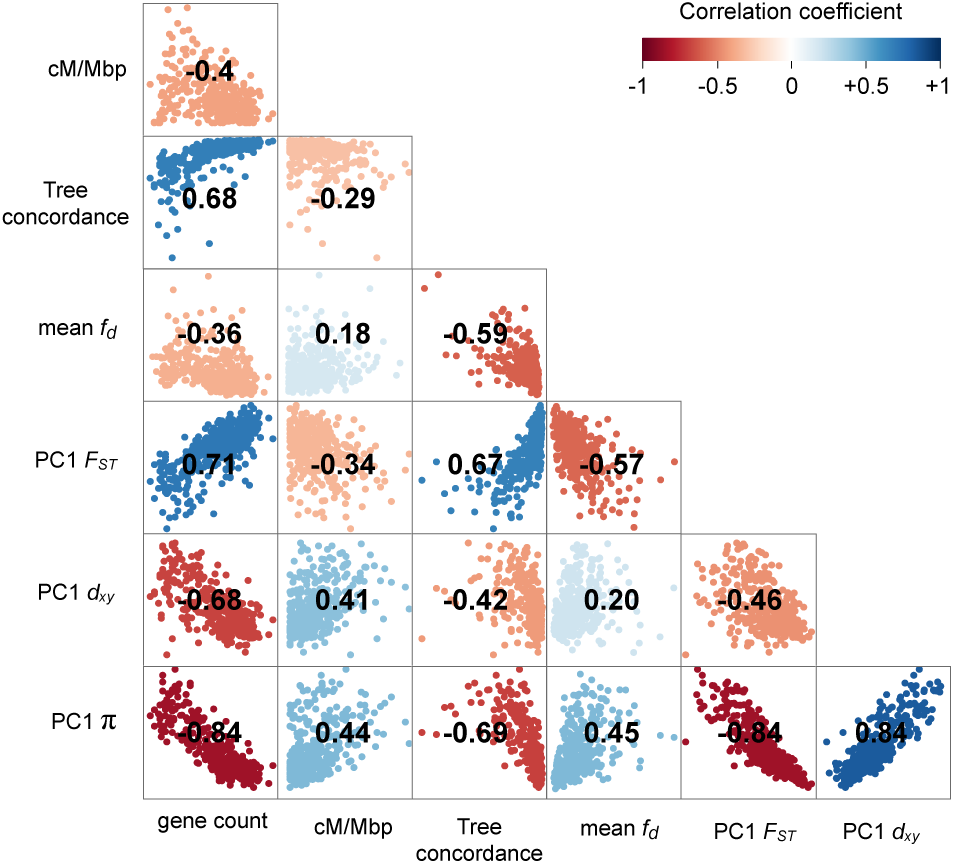
Correlations between measures of diversity and intrinsic features reveal the impact of heterogeneous indirect selection. Matrix of pairwise correlations between PC1 *F_ST_*, PC1 *d*_xy_, PC1 π, tree concordance, mean *f_d_*, gene density, and recombination rate, all estimated in 500 kb non-overlapping windows. The heat map indicates the strength of the correlation and its sign. All correlations are statistically significant at *p <* 0.001. (See Fig. S8 for a more detailed correlation matrix and Fig. S9 for the correlation matrix from 100 kb windows.

In models of recurrent indirect selection, variation in its impact on associated sites is determined by the distribution of genomic features, including the local density of functional elements and the recombination rate (Maynard-Smith and Haigh 1974, Charlesworth et al. 1993). To test these theoretical predictions, we used our genome annotation and genetic map to calculate the number of protein coding genes and the average recombination rate (cM/Mbp) in each 500 kb window (Fig. 2E-F; Fig. S4; Fig. S5). Regions of the genome with more functional elements tended to have a lower recombination rate (*r* = −0.40; *p* < 0.0001), leading to large variation in the predicted strength of indirect selection among regions. Consistent with the theoretical predictions outlined above, we found strong correlations between PC1 π and gene count (*r* = −0.84; *p* < 0.0001) and PC1 π and recombination rate (*r* = 0.44; *p* < 0.0001; Fig. 3; Figs. S8 & S9), indicating that diversity is indeed lower in regions where selection is predicted to have stronger indirect effects. However, we did not observe a significant interactive effect of gene count and recombination rate on diversity (*p* = 0.057). This may be because the distribution of these features is correlated, making it difficult to tease apart their relative impacts.

Taken together, our results indicate that a common pattern of indirect selection has caused correlated genomic landscapes to evolve across this radiation. As predicted by theory (Maynard-Smith and Haigh 1974, Charlesworth et al. 1993, Langley et al. 2012) (Hudson and Kaplan 1995) and observed in diverse taxa (Pease and Hahn 2013, Burri et al. 2015, Corbett-Detig et al. 2015), genome-wide variation in the strength of indirect selection is caused by the heterogeneous distributions of intrinsic genome features, namely the density of functional elements and the local recombination rate.

### A known adaptive locus shows a strong deviation from the common pattern of differentiation

Because widespread signatures of indirect selection are often assumed to reflect the impact of heterogeneous background selection (BGS) across the genome, several authors have proposed that the correlation in *F*_ST_ across multiple pairs of taxa can be used as a baseline for detecting genomic regions that have been affected by positive selection (Berner and Salzburger 2015, Burri 2017b). More specifically, loci that have contributed to adaptation or speciation should be detectable as a positive deviation from the common pattern of differentiation (i.e., PC1), which is considered to reflect processes that are unrelated to ecologically relevant positive selection (Burri 2017b).

We have a unique opportunity to test the effectiveness of this comparative genomic approach by examining patterns of differentiation around a locus that is known to contribute to adaptation and speciation in subspecies *puniceus*. Using a candidate gene approach, Streisfeld et al. (Streisfeld et al. 2013) showed that the shift from yellow to red flowers in *puniceus* was caused by a *cis-*regulatory mutation in the R2R3-MYB transcription factor *MaMyb2*. Patterns of haplotype variation within the gene indicate that the red allele was subject to strong positive selection and rapidly swept to fixation in what is now the geographic range of the red ecotype (Stankowski and Streisfeld 2015).

To test for a linage-specific signature of differentiation at this locus, we examined patterns of Z-*F_ST_* in a relatively narrow region (∼3 Mbp) surrounding *MaMyb2*. (Fig. 4; Fig. S10). At the 500 kb scale, the comparison between the red and yellow ecotypes shows a sharp peak centered near the flower color locus. The peak is strongly elevated above PC1 Z-*F_ST_*(3.33 standard deviations), indicating that differentiation is indeed accentuated relative to the level observed between other taxon pairs. Further, other comparisons, including those with the red ecotype, do not show accentuated differentiation in this region, indicating that the signal is specific to this taxon pair. The peak is even more pronounced in the analysis at the 100 kb scale, rising 9.5 standard deviations above PC1 Z-*F_ST_*. At this scale, the signature of positive selection is visible in other comparisons that include the red ecotype, though the signal is less pronounced than in the comparison with the yellow ecotype (Fig. S10).

**Figure 4.**
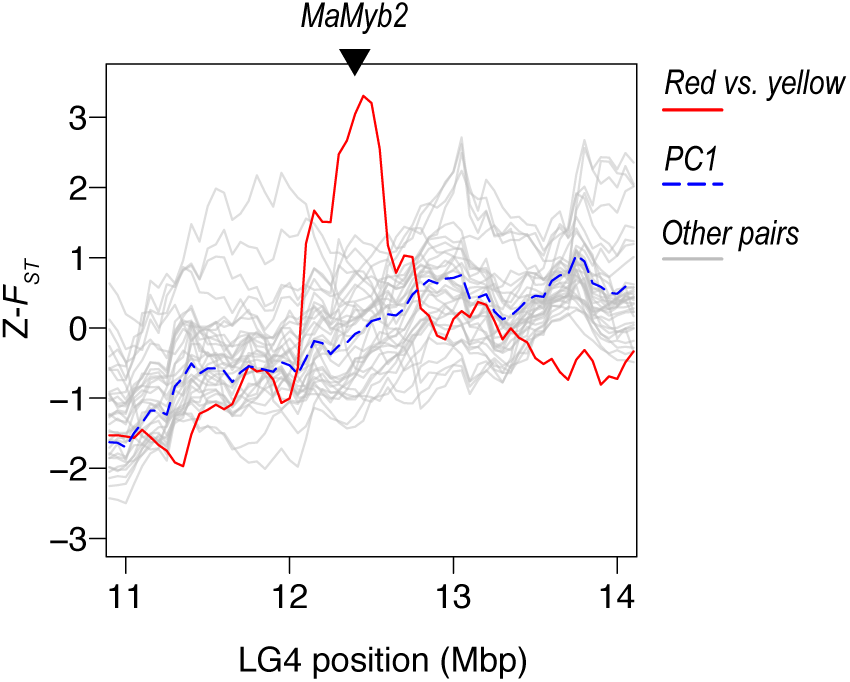
A large-effect adaptive locus shows a lineage-specific signature of positive selection. Plots of Z-transformed *F_ST_*across the genome, estimated in 500 kb sliding windows (step size 50 kb). The red line shows values between the red and yellow ecotypes of subspecies *puniceus*, while the gray lines show the values of all other comparisons. The dashed blue line shows the first PC calculated across all of the comparisons. The triangle marks the position of the gene *MaMyb2*. A *cis-*regulatory mutation that is tightly linked to this gene is responsible for the shift from yellow to red flowers. See Fig. S10 for the same plot made for 100 kb windows.

### Correlated differentiation landscapes emerge rapidly following a population split

Although correlated landscapes may evolve due to the indirect effects of widespread background selection, it is also conceivable that they could evolve due to other evolutionary processes. To gain insight into the role that background selection may have played in shaping these patterns, we next tested several hypotheses about how BGS is expected to shape the genomic landscape over time. To do this, we used the level of genetic distance between each pair of taxa as a proxy for their divergence time and constructed a temporal picture of genomic landscape evolution that spans the first million years following a population split.

In a verbal model, Burri (Burri 2017b) made several predictions about how correlations between measures of variation and genome features should be impacted by BGS following a population split. First, the correlations between diversity (π) and intrinsic features should be relatively consistent over time, as heterogeneous BGS will continue to constrain patterns of diversity in each daughter population after the split. Second, π and *d_xy_* should remain highly correlated with one another, because *d_xy_* is largely influenced by ancestral diversity at these short timescales. Third, levels of genome-wide differentiation (*F_ST_*) initially should be low and vary stochastically across the genome, due mainly to the sampling effect that accompanies a split. Therefore, *F_ST_* should not be correlated with the distribution of genomic properties early in divergence. However, as time passes, and the indirect effects of ancestral and lineage-specific BGS accumulate, levels of differentiation should gradually become more correlated with underlying genome features.

To test these hypotheses, we computed correlations between relevant window-based measures of variation and genomic features between all 36 pairs of taxa. We then plotted these correlations on corresponding measures of between-taxon *d*_a_, which ranged from very low (0.05%) between the parapatric red and yellow ecotypes of subspecies *punicieus*, up to 1.3% in allopatric comparisons that included *M. clevelandii*. We used *d*_a_ (*d_xy_* minus mean π), because it corrects for sequence variation present in the common ancestor, making it a better proxy of recent divergence time than *d_xy_*.

This analysis revealed very clear temporal signatures of genomic landscape evolution, some of which were consistent with the above predictions (Fig. 5; See Fig. S11 for results with *d_xy_* as a proxy for time). First, the presence of strong correlations between π (mean of the two taxa) and the distribution of genomic features barely changed with increasing divergence time between populations (Fig. 5A & 5B; Table S6). This is consistent with the prediction that diversity landscapes are inherited from the ancestor and are then maintained by the indirect effects of selection in each daughter population following the split. Second, the relationships between *F_ST_* and gene count, *F_ST_*and recombination rate, and *F_ST_* and π all become stronger as *d*_a_ increases (Fig. 5C & 5D; Table S6). Thus, as predicted, the differentiation landscape increasingly reflects the distribution of intrinsic features as divergence time increases.

**Figure 5.**
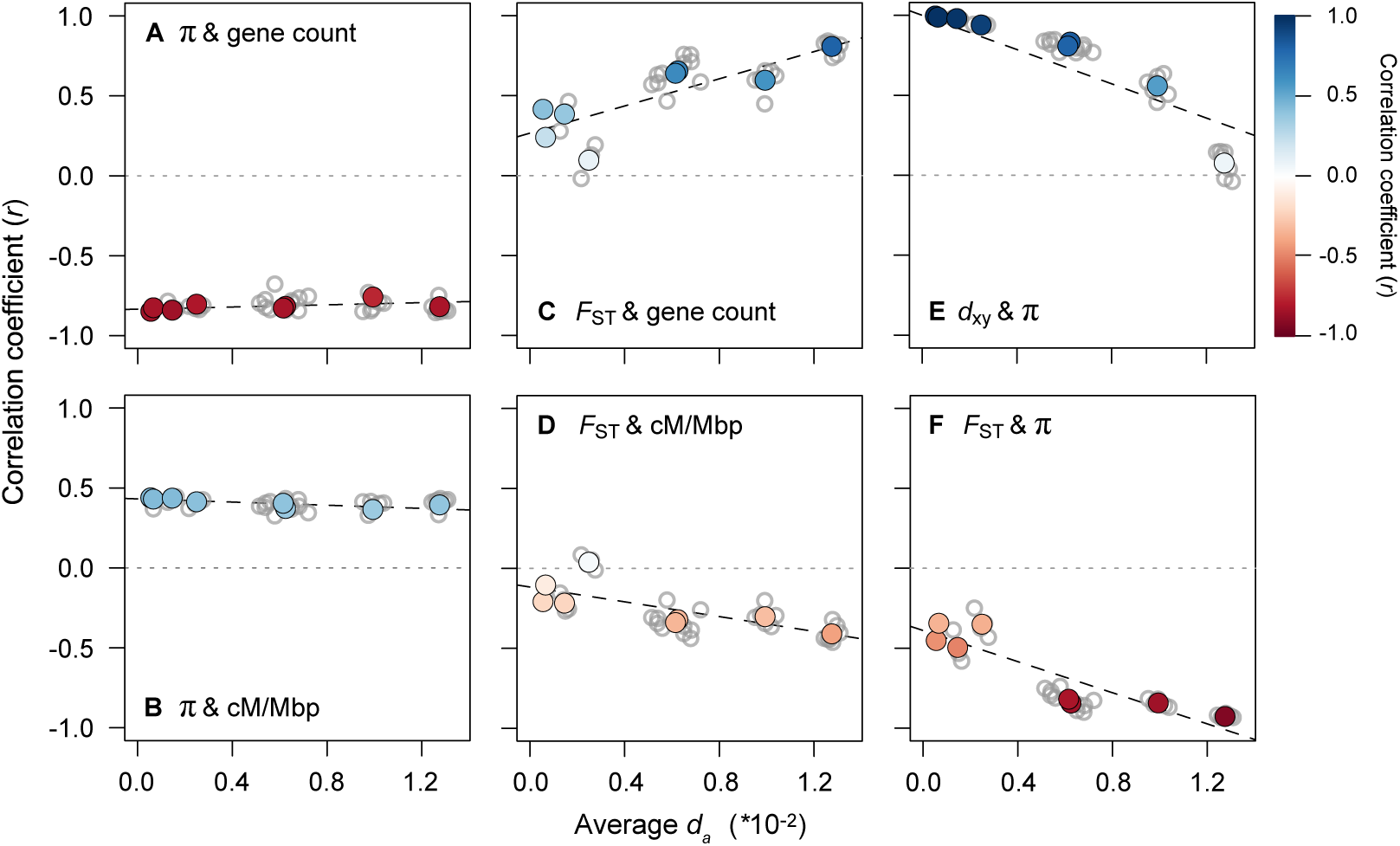
The range of divergence times reveals static and dynamic signatures of recurrent indirect selection. Correlations between variables (500 kb windows) for all 36 taxonomic comparisons (gray dots) plotted against the average *d*_a_ as a measure of divergence time. The left panels show how the relationships between π (each window averaged across a pair of taxa) and (A) gene count and (B) recombination rate vary with increasing divergence time. The middle panels (C & D) show the same relationships, but with *F_ST_*. The right panels show the relationships between (E) *d_xy_* and π and (F) *F_ST_*and π. The regressions (dashed lines) in each plot are fitted to the eight independent contrasts (colored points) obtained using a phylogenetic correction. The color gradient shows the strength of the correlation.

However, two of the observed patterns differed markedly from existing predictions (Burri 2017b). First, π and *d_xy_* did not remain highly correlated over this relatively short timeframe. Although the correlation is almost perfect at the earliest time points (*r* > 0.99), it decays rapidly and is not significantly different from zero in the most diverged taxon pairs (Fig. 5E). In addition, the correlations between *F_ST_*and proxies of the strength of indirect selection are much higher than expected at these early divergence times. For example, for the red and yellow ecotypes, the correlations between *F_ST_*and gene density (*r* = 0.415), recombination rate (*r* = −0.208), and mean π (*r* = −0.452) are substantial and highly significant (*p* < 0.0001) at a very early stage of divergence. The strength of these correlations also increases rapidly, with the diversity and differentiation landscapes almost perfectly mirroring one another in the most divergent comparisons (Fig. 5F; *r* = −0.94; Fig. 6; Fig. S12). As our results only partially match the above predictions, it is possible that other forces aside from BGS are responsible for the rapid emergence of differentiation landscapes in this system. Therefore, we next considered the potential for other evolutionary processes to generate these patterns.

**Figure 6.**
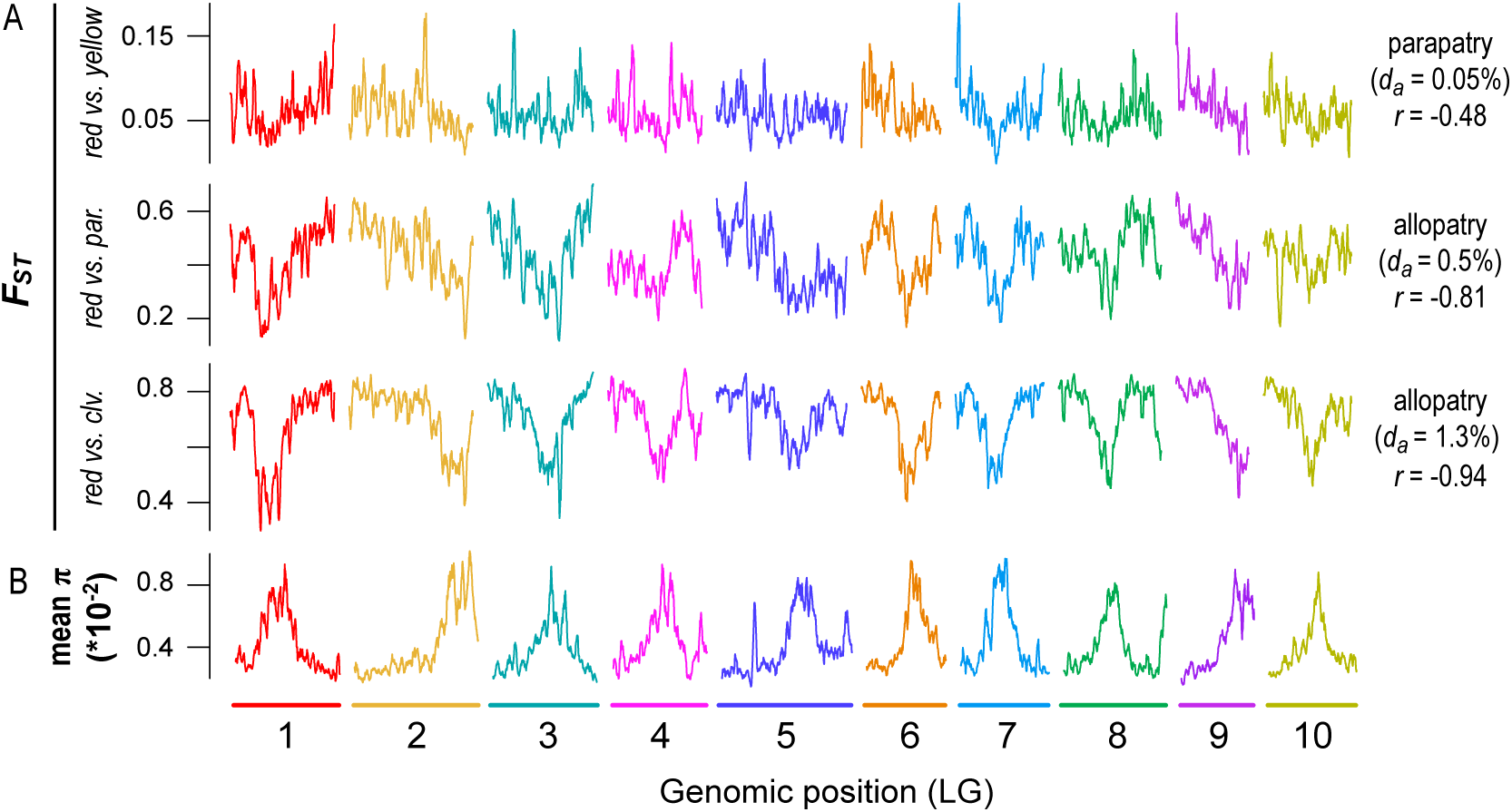
Emergence of a heterogeneous differentiation landscape across one million years of divergence. A) Plots of *F_ST_*(500 kb windows) across the genome for pairs of taxa at early (red vs. yellow), intermediate (red vs. *parviflorus*), and late stages (red vs. *M. clevelandii*) of divergence. B) Average nucleotide diversity (for the red ecotype of subspecies *puniceus*, yellow ecotype of subspecies *puniceus*, subspecies *parviflorus*, and *M. clevelandii*) across the genome in 500 kb windows. The geographic distribution (parapatric or allopatric), sequence divergence (*d_a_* × 10^-2^), and correlation between *F_ST_* and mean π are provided next to each taxon pair.

### Simulations suggest that adaptation has played an important role in genomic landscape evolution

To assess the plausibility of different modes of selection for generating the observed genomic landscapes and temporal patterns, we used individual-based simulations in SLiM (Haller et al. 2019, Haller and Messer 2019). Our basic model consisted of an ancestral population (N = 10,000) that evolved for 10N generations and then split into two daughter populations that diverged for a further 10N generations. Each individual carried a 21 Mbp chromosome (similar to the physical size of a bush monkeyflower chromosome), partitioned into three equally sized regions. In the central third, all mutations were neutral, while some mutations in the two distal ends could affect fitness, depending on the scenario. While simple, this partitioning scheme generates broad-scale variation in the strength of indirect selection across the chromosome and roughly approximates the distributions of genomic features in this system (e.g., LG 7 in Fig. 2).

We implemented six modifications to this basic model: (*i*) *Neutral divergence*: no mutations affect fitness; (*ii*) *Background selection* (BGS): non-neutral mutations are deleterious; (*iii*) *Bateson-Dobzhansky-Muller incompatibility* (BDMI): As *ii*, except that after the split, a fraction of non-neutral mutations is deleterious in one population and neutral in the other (randomly chosen). (*iv*) *Positive selection*: non-neutral mutations have positive fitness effects. (*v*) *BGS and positive selection*: *ii* and *iv combined*; (*vi*) *Local adaptation:* As *iv*, but after the split some non-neutral mutations are beneficial in one population and neutral in the other. Migration between populations was allowed only in scenarios *iii* and *vi*. See Methods for more details and Table S7 for parameter values. To summarize the results, measures of variation (π, *d_xy_*, and *F_ST_*) were calculated in 500 kb regions at 10 time points for each simulation.

This broad range of scenarios generated quite different patterns of variation, both through time and across the chromosome (Fig. 7; Fig. S13). As expected, the neutral model did not produce heterogeneous genomic landscapes. Background selection produced a diversity landscape characterized by higher diversity in the unconstrained central third of the chromosome (Fig 7; Fig S13). Consistent with the predictions of Burri (2017a), π and *d_xy_*remained highly correlated over the course of the simulations. However, even with a high proportion of deleterious mutations (10%) and substantial fitness effects (1%), the variation in π and *d_xy_* was modest, as was variation in the differentiation landscape (Fig. 7).

**Figure 7.**
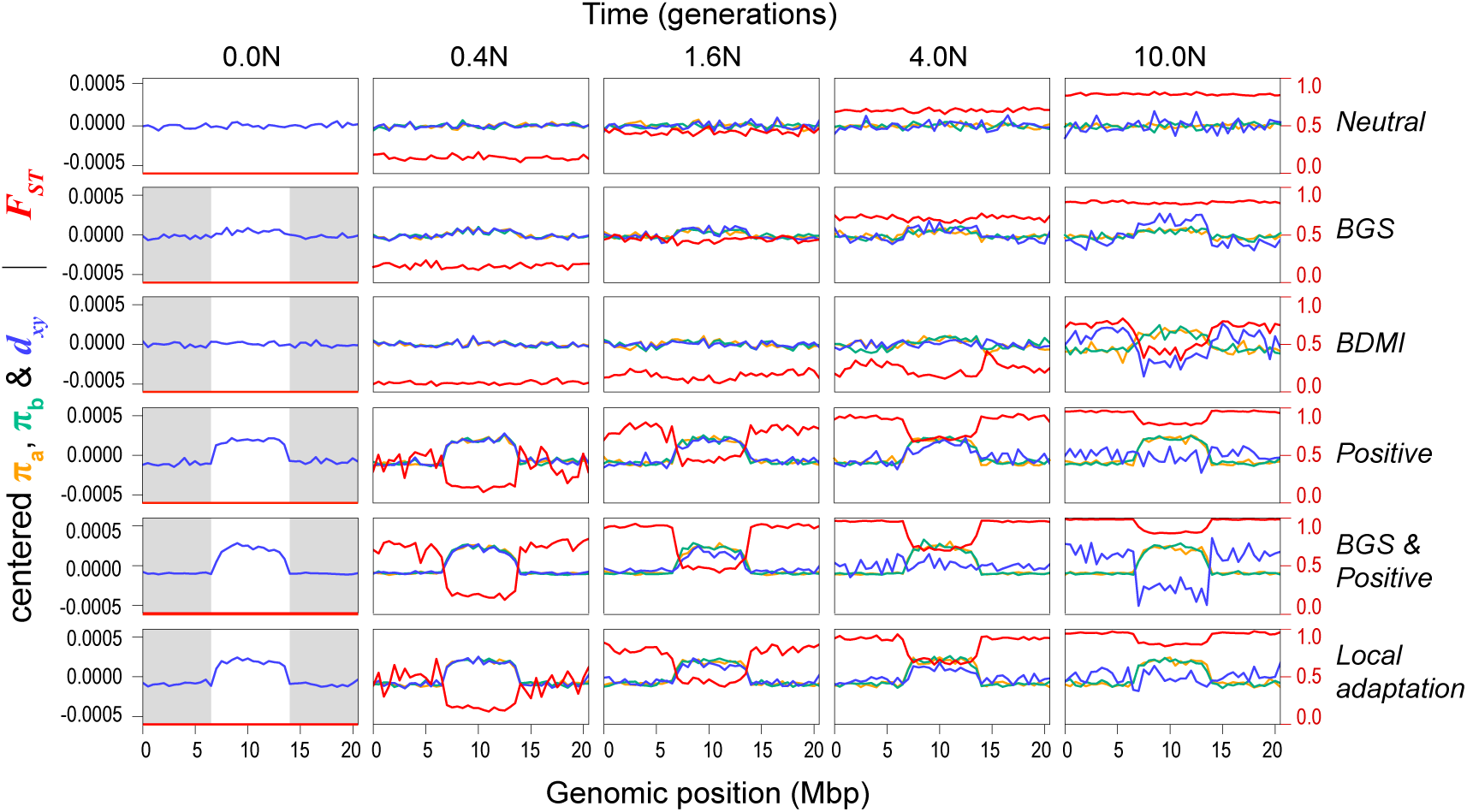
Genomic landscapes simulated under different divergence histories. Each row of plots shows patterns of within- and between-population variation (π, *d*_xy_, and *F_ST_*) across the chromosome (500 kb windows) at five time points (N generations, where N = 10000) during one of scenarios The selection parameter (Ns, where s = Ns/N), proportion of deleterious (-) and positive mutations (+), and number of migrants per generation (Nm; 0 unless stated) for these simulations are as follows: (*i*) neutral divergence (no selection), (*ii*) background selection (-Ns = 100; -prop = 0.01), (*iii*) Bateson-Dobzhansky-Muller incompatibilities (BDMI) (-Ns = 100, -prop = 0.05, Nm = 0.1), (*iv*) positive selection (+Ns = 100, +prop = 0.01), (*v*) background selection and positive selection (-Ns = 100, - prop = 0.01; +Ns = 100, +prop = 0.005), and (*vi*) local adaptation (+Ns = 100, +prop = 0.001, Nm = 0.1). The gray boxes in the first column show the areas of the chromosome that are experiencing selection, while the white central area evolves neutrally. Note that π (in populations a and b) and *d_xy_* have been mean centered so they can be viewed on the same scale. Un-centered values and additional simulations with different parameter combinations and more time points can be found in Fig. S13.

Under some conditions, the BDM incompatibility model produced patterns of variation that were more similar to our empirical findings than the BGS model (Fig. 7, S13). In all simulations, patterns of diversity and divergence were relatively homogenous at the time of the split. However, selection against gene flow induced by incompatibility loci on the chromosome ends caused patterns of variation to become highly structured over time. Indeed, higher π, lower *d_xy_*, and lower *F_ST_* were observed in the neutrally evolving chromosome center due to the homogenizing effects of higher gene flow in this region. As in our empirical data, the correlation between π and *d_xy_* decayed over the course of some simulations, and even became negative depending on rates of gene flow and selection. Although we simulated this scenario starting with undifferentiated populations (i.e., primary contact), widespread BDMIs should produce the same patterns for populations that diverged in allopatry and came back into secondary contact. This is because gene flow will have a stronger homogenizing effect in regions of the genome that are not associated with incompatibilities (Durrett et al. 2000).

The three models with positive selection (Positive, BGS & Positive, and Local adaptation) all produced highly heterogeneous genomic landscapes across the range of parameter values explored (Fig. 7; Fig. S13). In all cases, positive selection in the ancestor created a highly heterogeneous diversity landscape that was inherited by each daughter population and was maintained for the duration of the simulation. As in our data, π and *d_xy_* were perfectly correlated at early divergence times, but the correlation decayed rapidly and even became negative in scenarios with stronger selection. Unlike in the BDMI scenario, where the correlation between π and *d_xy_* decayed due to higher gene flow in the chromosome center, it broke down in these simulations because positive selection caused *d_xy_* to increase more rapidly in the chromosome ends (Fig. S13). Highly heterogeneous differentiation landscapes, characterized by lower *F_ST_*in the chromosome center, emerged rapidly in all three scenarios. This pattern appeared irrespective of the parameter values examined and was not heavily influenced by other factors, like gene flow, the inclusion of BGS, or whether adaptation occurred from standing variation. However, some of these factors may have a larger influence in simulations with different non-neutral mutation rates and selection strengths.

Overall, the results of these simulations suggest that background selection (BGS) is not primarily responsible for heterogeneous genomic landscapes. Although our simulations should be interpreted cautiously, because we have not thoroughly explored parameter space associated with each model, the results of other recent simulation studies also support this conclusion. For example, Matthey-Doret & Whitlock (Matthey-Doret and Whitlock 2018) simulated population divergence with BGS under different scenarios with simulations using parameters estimated from humans and stickleback. They found that BGS was unable to generate heterogeneous differentiation landscapes over short timescales, with or without gene flow. Similarly, Rettelbach *et al*. (Rettelbach et al. 2019) simulated the evolution of diversity landscapes using empirical estimates of recombination rate and functional densities from the collared flycatcher genome. They found that BGS was able to generate modest variation in levels of diversity across chromosomes, but they concluded that it was not sufficient to explain the pronounced dips observed in empirical studies of flycatcher genomes (e.g., (Burri et al. 2015)). Both of these studies suggest that other processes (probably positive selection) are responsible for generating the observed patterns (Matthey-Doret and Whitlock 2018, Rettelbach et al. 2019).

Our simulations suggest that divergence histories involving positive selection and/or incompatibilities are plausible explanations for heterogeneous genomic landscapes that emerge rapidly after a population split. However, in order to account for the presence of strong correlations between measures of variation and genomic features across multiple comparisons, these scenarios assume that the genomic basis of adaptation and/or reproductive isolation is diffuse, shared across multiple taxa, and evolves rapidly. Although this was once considered unlikely, recent studies suggest that these conditions may be common. For example, adaptation and speciation are now thought to be highly polygenic (Rockman 2012, Jiggins and Martin 2017, Martin et al. 2019), and recent evidence suggests that adaptation draws heavily on ancestral mutations, rather than those arising independently in multiple, related descendent lineages (Barrett et al. 2008, Nelson and Cresko 2018, Marques et al. 2019). Similarly, the interaction between widespread hybrid incompatibilities (intrinsic or extrinsic) and intrinsic genomic features is thought to have caused similar patterns of genome-wide variation to evolve between different pairs of hybridizing taxa (Martin et al. 2019).

Given the rapid and extensive trait diversification that has occurred during the bush monkeyflower radiation (Stankowski et al. 2015, Chase et al. 2017, Sobel et al. 2019), positive selection has probably played an important role in shaping patterns of genome-wide variation across the group. This includes the striking divergence of floral traits (Fig. 1A), which is thought to be due to divergent pollinator-mediated selection across southern California (Grant 1981, 1993b, a). In the taxa that have been studied in detail, crossing experiments have shown that most of these traits are polygenic, and there is evidence that they are targets of selection (Sobel and Streisfeld 2015, Stankowski et al. 2015). Intrinsic genetic incompatibilities may also influence patterns of variation between some of the taxa, but they are unlikely to explain the correlated landscapes that have evolved across the whole group, as most of these taxa show little or no evidence for intrinsic incompatibilities between them (McMinn 1951, Sobel and Streisfeld 2015). However, local adaptation also can generate a porous isolating barrier (Wu 2001), so it is likely that gene flow has contributed to genomic landscape evolution among some taxa, particularly those that are geographically proximate and/or hybridize in areas where their distributions overlap.

### Radiation-wide evidence for selection against gene flow

To determine if gene flow has contributed to the evolution of these correlated genomic landscapes, we first tested for evidence of introgression among these taxa using *D*-statistics (Green et al. 2010). Patterson’s *D* measures asymmetry between the numbers of sites with ABBA and BABA patterns (where A and B are ancestral and derived alleles, respectively) across three in-group taxa and an out-group (in our case, *M. clevelandii*) that have the relationship (((P1, P2), P3) O). A significant excess of either pattern gives a non-zero value of *D*, which is taken as evidence that gene flow has occurred between P3 and one of the in-group taxa (Green et al. 2010). Overall, these tests provide evidence that gene flow has occurred during this radiation. All 48 of the appropriate four-taxon combinations yielded significant non-zero values of Patterson’s *D* (*p* < 0.0001; Table S7; Fig. S14). The admixture proportions (*f*) for each test indicate that an average of 2.5% of the genome has been transferred between pairs of in-group taxa, but the proportion varies among the tests, ranging from less than 1% to more than 10% (Fig. S4.

Although gene flow initially involves the exchange of whole genomes between populations, selection against gene flow reduces effective migration (*m*_e_) in regions of the genome that are associated with barrier loci (Jiggins and Martin 2017, Ravinet et al. 2017). Therefore, porous isolating barriers should result in a heterogeneous pattern of introgression across the genome (Barton and Hewitt 1985, Wu 2001). To test this hypothesis, we calculated the *f_d_* statistic, a version of the admixture proportion modified for application to genomic windows (Martin et al. 2015). As predicted, estimates of *f_d_*, varied markedly among genomic regions (Fig. 2; Fig. S14). Most windows showed levels of *f_d_*at or near zero, indicating that foreign alleles have been purged from these regions by selection. However, a proportion of windows showed substantial admixture proportions. In some tests, values of *f_d_* exceeded 0.5, indicating that more than half of the sites in some windows have been shared between taxa.

Because gene flow opposes differentiation, we would expect to see lower *F_ST_* in regions with a higher proportion of introgressed variants. Consistent with this prediction, we observed a substantial negative correlation (*r* = −0.57; *p* < 0.0001) between mean *f_d_* (for each window, averaged over the 48 tests) and PC1 *F_ST_.* This result is not driven by a limited number of the four-taxon tests, as 44 of the 48 comparisons showed the same significant negative relationship (Table S15). Also consistent with models of widespread selection against gene flow, regions of the genome with higher *f_d_*scores tended to have higher diversity (*r* = 0.45; *p* < 0.0001), lower tree concordance scores (*r* = −0.59; *p* < 0.0001), fewer functional genes (*r* = −0.36; *p* < 0.0001), and a higher recombination rate (*r* = −0.18; *p* < 0.0001).

Overall, these results indicate that widespread gene flow has played a key role in the formation of the genomic landscape in this system. In addition to reductions in diversity and increased differentiation owing to selection against gene flow, the persistence of introgressed variants has probably resulted in higher levels of diversity in regions with fewer genes and higher recombination rate. The four-taxon tests show that the impact of gene flow is widespread across the radiation, though some caution should be exercised when interpreting the specific pattern of individual introgression events.

Given that there is the potential for gene flow between so many taxa and ancestral lineages, it is difficult to infer the source and timing of individual admixture events. For example, rather than reflecting recent gene flow between all pairs of taxa, some introgression events may have occurred deeper in history, and their consequences inherited by multiple taxa. Although more sophisticated methods will be needed to model gene flow across this group, these results clearly show that it has contributed to the rapid evolution of correlated genomic landscapes during this radiation.

### Conclusions and implications for understanding genomic landscape evolution and the basis of adaptation and speciation

Facilitated by a new chromosome-level genome assembly, we have shed light on the causes of correlated genomic landscapes across a radiation of monkeyflowers. Adaptive divergence and gene flow are hallmarks of rapid radiations (Schluter 2000), and our data suggest that the indirect effects of selection resulting from these processes have contributed to a common pattern of differentiation among these taxa. Our ‘time-course’ analysis shows that the common landscape emerged rapidly after populations split and has become more correlated with the distribution of genomic features as divergence time has increased. Although background selection may play some role, its effects are probably too subtle to have made a strong contribution to the correlations during over the timeframe of this radiation.

In addition to enhancing our understanding of the processes that have shaped the genomic landscape during this radiation, our study contributes toward a more general understanding of the role that natural selection plays in shaping genome-wide variation. In line with the findings of other recent studies, our results indicate that little, if any, of the genome evolves free of the effects of natural selection (Hahn 2008, Corbett-Detig et al. 2015). Moreover, our ‘time-course’ analysis shows that between-taxon signatures of selection can emerge very rapidly after a population split and can be substantial, even between populations at the early stages of divergence. Overall, these results suggest that patterns of between-population variation cannot be understood without taking the effects of natural selection into account.

An important point arising from this work is that multiple divergence histories can generate heterogeneous differentiation landscapes that are correlated both among taxa and with the distribution of intrinsic genomic properties. When divergence is recent, possible explanations include polygenic adaptation across multiple populations and porous barriers to gene flow arising from divergent ecological selection and/or intrinsic incompatibilities. Although it has often been assumed that recurrent background selection is the primary cause of correlated landscapes, it is important to remember that all forms of selection can indirectly impact levels of variation at associated sites. In fact, our simulations show that alternative explanations may be more likely when divergence times are short and there is opportunity for gene flow among taxa. Thus, we advocate for a more nuanced approach when interpreting correlated differentiation landscapes, rather than assuming that they are caused by a single evolutionary process.

Finally, our results have important practical implications for studies attempting to identify the genomic basis of adaptation and speciation from patterns of genome-wide variation. Indeed, detecting these loci is a major goal of adaptation and speciation studies, and genome scans are now commonly used to identify promising candidate regions (Ravinet et al. 2017). In an effort to correct for the potentially confounding effects of background selection (BGS), it has been proposed that the correlation in *F_ST_*among multiple population pairs can be used as a baseline for outlier detection (Berner and Salzburger 2015, Burri 2017b). A core premise of this method is that correlated differentiation landscapes are caused primarily by BGS, so removing this signature should expose the loci relevant to adaptation and/or speciation. Although this approach may be successful in identifying large-effect loci targeted by lineage-specific positive selection, as we illustrated for the *MaMyb2* locus, we caution against treating the common pattern of differentiation simply as background noise. Rather than parsing out the effects of BGS, studies that use this approach may actually be discarding the genomic signature of polygenic adaptation and speciation.

## Data Accessibility

Raw sequencing reads used for the genome assembly, linkage map construction, and genome resequencing and are available on the Short-Read Archive (SRA) under the bioproject ID xxx. The genetic map, annotation, VCF file, and analysis scripts have been deposited on DRYAD. The reference assembly is available at mimubase.org. Code used to run the simulations is available on Github at: https://github.com/mufernando/mimulus_sims.

## Acknowledgments

We thank William A. Cresko, Thomas Nelson, Roger K. Butlin, Anja M. Westram, Mark Ravinet, Sarah. P. Otto, and Michael C. Whitlock for stimulating discussion. John Willis performed the Illumina mate-pair library preps used for the genome assembly. Julian Catchen, Clay Small, Susan Bassham, Mark Rausher and Janna Fierst provided technical or analytical support or advice. Doug Turnbull and Maggie Weitzman conducted the Illumina sequencing at the University of Oregon Genomics Core facility. Thomas Nelson, Martin Garlovsky, Roger K. Butlin, Anja M. Westram and Jeff Ross-Ibarra provided comments on an earlier version of this manuscript. We thank Reto Burri, Stuart J.E. Baird, and two anonymous reviewers for providing insightful comments that greatly improved the paper. We also thank people who contributed to insightful discussions at the 2017 Speciation Genomics Meeting, Cambridge UK, the 2019 Gordon Speciation Conference, Ventura CA, USA, and on Twitter. Funding was provided by the National Science Foundation grant DEB-1258199 to MS and the Sloan Foundation to PR.

## Materials and Methods

### Genome Assembly

We used a combination of short-read Illumina and long-read Single Molecule, Real Time (SMRT) sequencing to assemble the genome of a single individual from the red ecotype of *M. aurantiacus* subspecies *puniceus* (population UCSD; Table S1). Genomic DNA was isolated from leaf tissue using either ZR plant/seed DNA miniprep kits (Zymo Research) or GeneJet Plant Genomic DNA purification kits (Thermo Fisher). Illumina libraries were generated following the *Allpaths-LG* assembly pipeline (Gnerre et al. 2011), which included a single fragment library with average 180 bp insert size and three mate pair libraries (average insert sizes: 3.5-5 kb, 5-7 kb, and 7-13 kb). Libraries were sequenced on the Illumina HiSeq 2500 using paired-end 100 bp reads. An initial scaffold-level assembly was performed with *Allpaths-LG* using default parameters and the *haploidify* function enabled. This assembly yielded 11,123 contigs (N50 = 40.5 kb) and 2,299 scaffolds (N50 = 1,310 kb), for a total assembly size of 193.3 Mbp. Long-read sequencing was performed from the same individual using 12 SMRT cells sequenced on the Pacific Biosystems RS II machine at Duke University. We obtained a total of 6.4 Gbp of sequence, which corresponds to ∼21× coverage of the genome. The PacBio reads were used to re-scaffold the *Allpaths-LG* scaffolds using *Opera-LG* (Gao et al. 2016). This reduced the number of scaffolds to 1,547 (N50 = 1,578 kb).

We then manually improved the scaffold containing the flower color gene *MaMyb2* (Streisfeld et al. 2013). We first aligned this scaffold to a previously published draft sequence assembly from this same individual (Stankowski et al. 2017), which was generated using Illumina short-reads and the *Velvet* assembler (Zerbino and Birney 2008). We used long range PCR and cloning to generate Sanger sequences across three regions within 20 kb of *MaMyb2* that did not assemble well. Genomic DNA was amplified using Phusion high fidelity polymerase (NEB). PCR products were cloned into the pCR2.1 TOPO-TA vector (Life Technologies), and purified plasmids were sequenced with Sanger technology. Resulting sequences were aligned to the scaffold containing *MaMyb2*, and new PCR primers were designed to sequence internal fragments until the entire insert was sequenced. Using this approach, we sequenced a total of 9,824 bp across the three regions. The reference sequence in the assembly was corrected manually to match the Sanger data.

Finally, we gap filled the assembly using the PacBio data and the program *PBJelly* (English et al. 2012). Resulting scaffolds were assembled into pseudomolecules using *Chromonomer* (http://catchenlab.life.illinois.edu/chromonomer/), according to the online manual. This software anchored and oriented scaffolds based on the order of markers in a high-density linkage map (see below) and made corrections to scaffolds when differences occurred between the genetic and physical positions of markers in the map. A final round of gap filling with *PBJelly* was performed to fill any gaps that were created by broken scaffolds in *Chromonomer*. To assess the completeness of the gene space in the assembly, we used both the BUSCO and CEGMA pipelines to estimate the proportion of 956 single copy plant genes (BUSCO) or 248 core eukaryotic genes (CEGMA) that were completely or partially assembled (Parra et al. 2007, Simao et al. 2015). The proportion of these genes present in an assembly has been shown to be correlated with the total proportion of assembled gene space, and thus serves as a good predictor of assembly completeness.

### Construction of high-density linkage map

We generated an outbred F_2_ mapping population by crossing two F_1_ individuals, each the product of crosses between different greenhouse-raised red and yellow ecotype plants collected from one red ecotype and one yellow ecotype population (populations UCSD and LO, respectively; Table S1). We then used restriction-site associated DNA sequencing (RADseq) to genotype F_1_ and F_2_ individuals. DNA was extracted from leaf material using Zymo ZR plant/seed DNA miniprep kits, and RAD library preparation followed the protocol outlined in Sobel and Streisfeld (Sobel and Streisfeld 2015). Libraries were sequenced on the Illumina HiSeq 2000 platform using single-end 100 bp reads at the Genomics Core Facility, University of Oregon.

Reads were filtered based on quality, and errors in the barcode sequence or RAD site were corrected using the *process_radtags* script in *Stacks v. 1.35* (Catchen et al. 2011, Catchen et al. 2013). Loci were created using the *denovo_map.pl* function of *Stacks*, with three identical raw reads required to create a stack, two mismatches allowed between loci for an individual, and two mismatches allowed when processing the catalog. Single nucleotide polymorphisms (SNPs) were determined and genotypes called using a maximum-likelihood (ML) statistical model implemented in *Stacks* and a stringent χ^2^ significance level of 0.01 to distinguish between heterozygotes and homozygotes. We then used the *genotypes* program implemented in *Stacks* to identify a set of 9,029 mappable markers. We specified a ‘CP’ cross design (F_1_ individuals coded as the parents), requiring that a marker was present in at least 85% of progeny at a minimum depth of 12 reads per individual, and we allowed automated corrections to be made to the data.

Linkage map construction was performed using *Lep-MAP2* (Rastas et al. 2016). The data were filtered using the *Filtering* module to include only individuals with less than 15% missing data and excluded markers that showed evidence for extreme segregation distortion (χ^2^ test, *P* < 0.01). To assign markers to linkage groups, we used the *SeparateChromosomes* module with a logarithm of odds (LOD) score limit of 20 and no minimum size for linkage groups (LG). This assigned 7,217 markers to 10 linkage groups, which matches the number of chromosomes in *M. aurantiacus*. The *JoinSingles* module was executed again with a LOD limit of 10 to join an additional 877 ungrouped markers to the 10 previously formed LGs. Fifty-seven singles that were not joined at this stage were discarded from the dataset. Initial marker orders were determined using sex-averaged and sex-specific recombination rates using the *OrderMarkers* module. For each LG, we conducted 10 independent runs using the Kosambi mapping function (*useKosambi=1*), with the dataset split into seven pseudo-families to take advantage of parallel processing. When multiple markers had identical genotypes, only the duplicate marker with the least missing data was used in marker ordering. We retained the marker order from the run with the best likelihood. After removing markers with an error rate > 0.05, the ML order was re-evaluated using the *evaluateOrder* flag. The map contained 8,094 informative loci from 269 F2 individuals, with an average of 3.5% ± SD 3.86 missing data per individual.

After the integration of our assembly and genetic map using the *Chromonomer* software (Amores et al. 2014), we made corrections to the map order based on the physical position of markers within assembled scaffolds. Using the output of *Chromonomer*, we identified markers that were out of order in the map compared to their local assembly order and aligned these markers to the assembly from *Chromonomer* using *Bowtie2* v. 2.2.5 (Langmead and Salzberg 2012) with the *very_sensitive* settings to obtain their physical order. We then re-estimated the map using the *evaluateOrder* flag in *Lep-MAP2* as described above, but with the marker order constrained to the physical order (*improveOrder=0*) and with all duplicate markers included in the analysis (*removeDuplicates=0*). After initial map construction, we removed 17 markers with an estimated error rate greater than 5% and estimated the map one last time using the same settings. The final map contained 7,589 markers across the 10 linkage groups.

### Genome annotation

Prior to genome annotation, the assembly was soft-masked for repetitive elements and areas of low complexity with *RepeatMasker* (RepeatMasker Open-4.0) using a custom *Mimulus aurantiacus* library created by *RepeatModeler* (RepeatModeler Open-1.0), Repbase repeat libraries (Jurka et al. 2005), and a list of known transposable elements provided by *MAKER* (Holt and Yandell 2011). In total, 30.99% of the genome assembly was masked by *RepeatMasker*. Repetitive elements were annotated with *RepeatModeler*. Hidden Markov Models for gene prediction were generated by *SNAP* (Korf 2004) and *Augustus* (Stanke and Waack 2003) and were trained iteratively to the assembly using *MAKER*, as described by Cantarel et al. (Cantarel et al. 2008). Training was performed on the 14.5 Mbp sequence from LG9. Evidence used by *MAKER* for annotation included protein sequences from *Arabidopsis thaliana*, *Oryza sativa*, *Solanum lycopersicum*, *Solanum tuberosum*, *Daucus carota*, *Vitis vinifera* (all downloaded from EnsemblPlants on 9 August 2016), *Salvia miltiorrhiza* (downloaded from Herbal Medicine Omics Database on 9 August 2016), *Mimulus guttatus v. 2* (downloaded from JGI Genome Portal on 9 August 2016), and all Uniprot/swissprot proteins (downloaded on 18 August 2016) (Goodstein et al. 2012, Nordberg et al. 2013, Kersey et al. 2016) (Herbal Medicine Omics Database; Uniprot). We filtered the annotations with *MAKER* to include: 1) only evidence-based information that also contained assembled protein support, and 2) those *ab initio* gene predictions that did not overlap with the evidence-based annotations and that contained protein family domains, as detected with InterProScan (Quevillon et al. 2005).

### Genome re-sequencing and variant calling

We collected leaf tissue from each taxon (Collection locations available in Table S3 and Fig. S2) and extracted DNA from dried tissue using the Zymo Plant/Seed MiniPrep DNA kit following the manufacturer’s instructions. We prepared sequencing libraries using the Kapa Biosystems HyperPrep kit, and libraries with an insert size between 400-600 bp were sequenced on the Illumina HiSeq 4000 using paired-end 150 bp reads at the Genomics Core Facility, University of Oregon.

We filtered raw reads using the *process_shortreads* script in *Stacks* v1.46 to remove reads with uncalled bases or poor quality scores. We then aligned the retained reads to the reference assembly using the BWA-MEM algorithm in *BWA* v0.7.15 (Li 2013). An average of 91.7% of reads aligned (range: 82.6-96.0%), and the average sequencing depth was 21x (range: 15.16 – 30.86x). We then marked PCR duplicates using *Picard* (http://broadinstitute.github.io/picard). We performed an initial run of variant calling using the UnifiedGenotyper tool in *GATK* v3.8 (McKenna et al. 2010) and filtered the data to remove variants with a mapping quality < 50, a quality depth < 4, and a Fisher Strand score > 50. We then used these variants to perform base quality score recalibration for each individual, before performing another run of the UnifiedGenotyper to call final variants. After the second run of variant calling, we removed variants with a mapping quality < 40, a quality depth < 2, and a Fisher Strand score > 60. The final dataset contained 13,233,829 SNPs across the nine taxa. Finally, we ran UnifiedGenotyper with the EMIT_ALL_SITES option to output all variant and invariant genotyped sites.

### Phylogenetic and principal component analyses

We used *RAxML* v8 to reconstruct the evolutionary relationships among the nine taxa by concatenating variant sites from across the genome. To investigate patterns of phylogenetic discordance across the genome, we also built trees from windows across the genome. We phased SNPs using *BEAGLE* v4.1 (Browning and Browning 2007), using a window size of 100 kb and an overlap of 10 kb. We incorporated information on recombination rate from the genetic map and did not impute missing genotypes. After phasing, we used *MVFtools* (https://www.github.com/jbpease/mvftools) to run *RAxML* from 100 kb and 500 kb nonoverlapping windows, with the *M. clevelandii* samples set as the outgroups. We then visualized the window trees in *DensiTree* v2.01 (Bouckaert 2010).

To assess concordance between the window-based trees and the whole-genome tree, we converted trees to distance matrices using the *Ape* package in R (Paradis et al. 2004). We then calculated the Pearson’s correlation coefficient between the distance matrix from each window and the whole-genome tree, with a stronger correlation indicating higher concordance with the whole-genome tree. We used one-dimensional autocorrelation analysis to determine if variation in tree concordance was randomly distributed across the genome. This involved estimating the autocorrelation between genomic position and tree concordance for each LG with a lag size of 2 Mbp. The significance of the observed value for each LG was determined from a null distribution of autocorrelation coefficients estimated from 1000 random permutations of the genome-wide data.

We also conducted a Principal Components Analysis (PCA) based on all variant sites from across the entire genome using *Plink* v. 1.90 (Chang et al. 2015). Initially, we ran the PCA with all 37 samples, but we consecutively re-ran the analysis by removing different taxa in order to assess clustering patterns among more closely related samples.

### Population genomic analyses

To examine how genome-wide patterns of diversity, differentiation, and divergence varied among taxa, we calculated nucleotide diversity (π), between-taxon differentiation (*F_ST_*), and between-taxon divergence (*d_xy_*), respectively, in non-overlapping and overlapping 100 kb (step size = 10kb) and 500 kb (step size = 50 kb) windows using custom Python scripts downloaded from https://github.com/simonhmartin/genomics_general. We calculated measures of differentiation and divergence across all 36 pairwise comparisons among the nine taxa, and diversity was estimated separately for each taxon. These scripts estimated π and *d_xy_* by dividing the number of sequence differences within a window (either within or between taxa) by the total number of sites in that window. To account for missing data, the script counted the number of differences between each sample, divided by the total number of variant sites that were genotyped within those samples, and then averaged across all pairs of samples. To provide an unbiased estimate of diversity and divergence, we incorporated invariant sites into the calculation by dividing the number of pairwise differences (within and between taxa, respectively) by the total number of genotyped sites (variant and invariant) within a window. *F_ST_* was calculated following the measure of *K_ST_* (Hudson et al. 1992), equation 9), but was modified to incorporate missing data using the same approach as π and *d_xy_*. We filtered the data separately for each taxonomic comparison, so that each site was required to be genotyped in at least three individuals for comparisons within the *M. aurantiacus* complex or at least two individuals for each comparison involving *M. clevelandii*.

We summarized the variation in each statistic across comparisons using a Principal Components Analysis (PCA), with taxon or taxon pair as the variables. Thus, across each window, the first principal component of π, *F_ST_*, and *d_xy_* provided multivariate measures that explained the greatest covariance in the data. We used a one-dimensional autocorrelation analysis and permutation test to determine if the genome-wide patterns of PC1 π, *F_ST_*, and *d_xy_*departed from a random expectation, as described above for tree concordance (see section ‘phylogenetic analyses’).

To examine the relationships among PC1 diversity, differentiation, and divergence, we estimated Pearson’s correlation coefficient among all three statistics across genomic windows. Further, we estimated correlations among these three statistics and tree concordance, gene density, and recombination rate. Recombination rate was estimated by comparing the genetic and physical distance (in cM/Mbp) between all pairs of adjacent markers on each LG from the genetic linkage map described above. We removed the top 5% of recombination rates, as these represented unrealistically high values of recombination. A minimum of three estimates was required to obtain an average recombination rate estimate within each window. Gene density was calculated from the number of predicted genes in each window, as determined from the annotation described above. Genes spanning two windows were counted in both.

To determine how the correlations among the statistics (diversity, differentiation, divergence, recombination rate, gene count) changed with increasing divergence time, we examined the correlation coefficient among all pairs of statistics individually for each of the 36 pairwise comparisons. Because diversity was measured within taxa, we calculated the mean value of π between each pair of taxa. Also, because many of the pairwise comparisons are non-independent, we applied the phylogenetic correction outlined by (Felsenstein 1985, Coyne and Orr 1989) to produce a statistically independent set of data points for this analysis.

We calculated the divergence time between *M. clevelandii* and *M. aurantiacus*, based on a corrected estimate of sequence divergence (*d*_a_) between individuals of *M. clevelandii* and all subspecies of *M. aurantiacus* combined. We then converted this value into a divergence time *T* (in generations) using the equation: *T* = *d*_a_/(2µ), where µ is the mutation rate, 1.5 x 10^-8^ (Koch et al. 2001, Brandvain et al. 2014). This value was then converted into years by multiplying by a generation time of two years.

### Simulations

To assess the plausibility of different scenarios in producing heterogeneous genomic landscapes, we implemented forward-time Wright-Fisher simulations using SLiM (Haller et al. 2019, Haller and Messer 2019). The basic model consisted of an ancestral population with a fixed population size of N=10,000 that split after 10N non-overlapping generations into two daughter populations, each with a fixed size of N. These populations were then allowed to evolve for a further 10N non-overlapping generations. We simulated a 21 Mbp chromosome with a recombination rate of 2×10^-8^ and a mutation rate of 10^-8^, both per base pair and per generation.

We explored the following six modifications of this basic model. (*i*) *Neutral evolution*: mutations did not impact fitness; (*ii*) *Background selection* (BGS): mutations in the ancestor and daughter populations were neutral in the middle third of the chromosome but can be deleterious in the chromosome ends; (*iii*) *Bateson-Dobzhansky-Muller incompatibility* (BDMI): Neutral and deleterious mutations occurred in the ancestor. After the split, we allowed migration between the two daughter populations. To simulate BDM incompatibilities, a fraction of the selected mutations was deleterious in each of the populations and neutral in the other. (*iv*) *Positive selection*: mutations in the ancestor and daughter population were neutral in the middle third of the chromosome but could be beneficial in the remaining regions. (*v*) *Local adaptation:* mutations in the ancestor and daughter population were neutral in the middle third of the chromosome but could be beneficial in the remainder. After the split, we allowed migration between the daughter populations. To simulate local adaptation, a fraction of the selected mutations was beneficial in each population and neutral in the other. Unlike the BDMI simulation, we allowed local adaptation from standing variation, so some of the mutations that were neutral in the ancestral population became beneficial after the split. *BGS and positive selection*: mutations in the ancestor and daughter population were neutral in the middle third of the chromosome but could be either deleterious or beneficial in the tails.

Each scenario was simulated with varying proportions of beneficial/deleterious to neutral mutations, and varying mean selective coefficients and migration rates, where applicable (Table S7). For each combination of parameters, we ran five replicate simulations. As described in Kelleher et al. (Kelleher et al. 2018), to improve simulation speed we recorded genealogies and non-neutral mutations in a tree sequence and added neutral mutations afterwards with *msprime* (Kelleher et al. 2016). We used *scikit-allel* (v1.1.8) (Miles and Harding 2017) to calculate π, *F_ST_,* and *d_xy_* in 500 kb regions. All of the SLiM code used is available on GitHub (https://github.com/mufernando/mimulus_sims).

### Tests for genome-wide admixture

We tested for introgression and quantified levels of admixture by calculating Patterson’s *D* and the admixture proportion (Green et al. 2010, Durand et al. 2011). Patterson’s *D* takes four taxa with the relationship (((P1, P2), P3) O) and looks for an excess of either ABBA or BABA sites, where A and B represent the ancestral and derived alleles respectively. We calculated Patterson’s *D* for all possible groups of three ingroup taxa based on the set of relationships inferred from the genome-wide (concatenated) dataset. *M. clevelandii* was used as the outgroup for all tests. The *D* statistic was calculated from allele frequency data for biallelic sites, with the ancestral allele called as the allele fixed within *M. clevelandii.* Therefore, only sites fixed within. *M. clevelandii* were included in the analysis. We converted genotype data into allele frequency data using a Python script download from https://github.com/simonhmartin/genomics. Patterson’s *D* was calculated from the proportion of ABBA and BABA sites in R using a custom script. To evaluate the significance of each test, we performed block jack-knifing with a block size of 500 kb. We then calculated the genome-wide admixture proportion, *f*, for all tests with a significant value of Patterson’s *D*. This test compares the excess of ABBA sites shared between P2 and P3 to the expected excess of ABBA sites between two completely admixed populations. The expected excess is calculated by splitting the samples from P3 into two different groups (P3a and P3b) and calculating the excess of ABBA to BABA sites with one group taking the place of P2 and the other the place of P3.

For tests with significant values of Patterson’s *D*, we then estimated levels of admixture in each 500 kb genomic window. Although Patterson’s *D* is a powerful test for detecting introgression genome-wide, it is not suited for estimating admixture in defined genomic regions (Martin et al. 2015). Thus, we calculated the four-taxon statistic *f_d_*, which is a modified version of the admixture proportion developed for estimating local admixture (Martin et al. 2015). We calculated *f_d_*in 500 kb non-overlapping windows, using Python scripts (https://github.com/simonhmartin/genomics_general). Because *f_d_* is designed to detect an excess of ABBA sites (*i.e.,* gene flow from P3 to P2), we switched the order of P1 and P2 in tests where *D* was negative (excess of BABA) in the genome-wide tests for introgression. Note that this does not affect the set of relationships in these comparisons. We summarized genome-wide variation in *f_d_* across the different tests by taking the average value of *f_d_* within each window and the maximum value of *f_d_* from the 48 different tests in each window, and estimated the Pearson’s correlation coefficient between these values other statistics.

## Supplementary tables

**Table S1.**
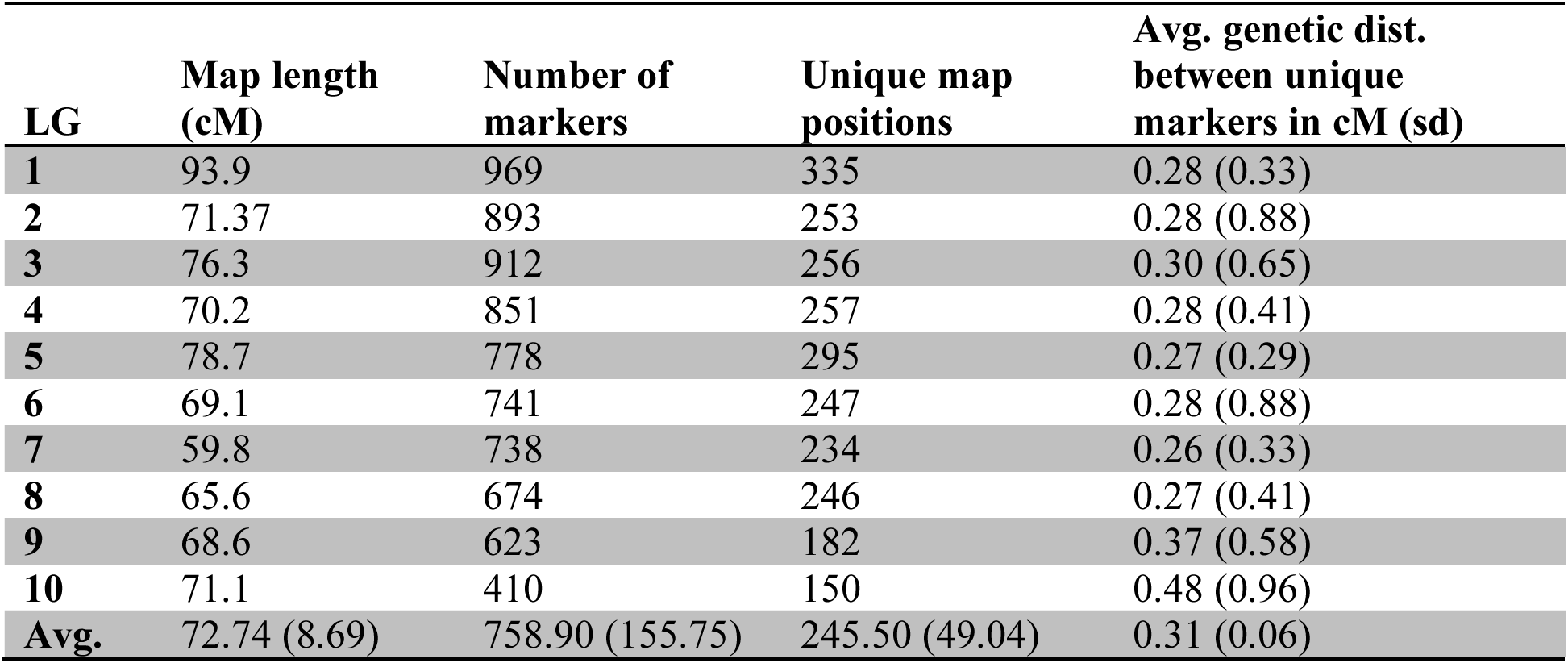
Summary of the genetic linkage map constructed using an F2 intercross between the red and yellow ecotypes of subspecies *puniceus*. The table includes map length in cM for each linkage group (LG), the number of markers associated with each LG, the number of unique map positions, and the average genetic distance in cM between each unique map position. Standard deviations are given in parentheses.

**Table S2.** Analysis of gene space completeness in the M. aurantiacus genome using CEGMA and BUSCO. The number and percent of core genes found in the final assembly are shown for each analysis (CEGMA, *n* = 248; BUSCO, *n* = 1440).

| Analysis | # Genes | % Found in Assembly |
| --- | --- | --- |
| **CEGMA Complete** | 233 | 93.95 |
| **CEGMA Partial** | 244 | 98.39 |
| **BUSCO total complete (duplicated)** | 1340 (61) | 93 (4.2) |
| **BUSCO Fragmented** | 29 | 2.0 |
| **BUSCO Missing** | 71 | 5.0 |

**Table S3.**
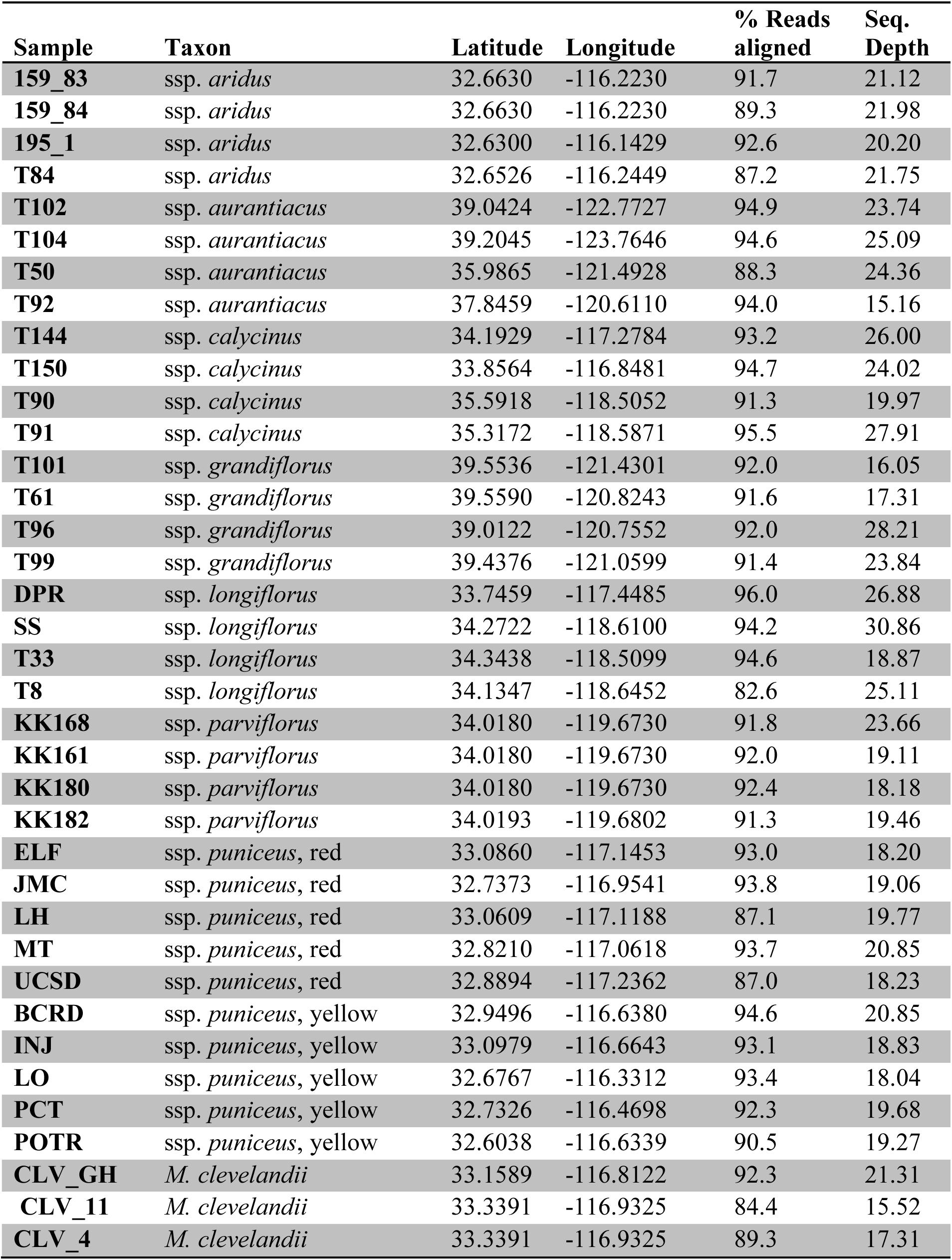
Sample information for the 37 sequenced individuals. Includes their taxon identity, sampling location, percent read alignment, and average sequencing depth.

**Table S4.** Loadings for principal component 1 calculated across all 36 pairwise comparisons (for *F_ST_* and *d_xy_*) or all nine taxa (for π).

| **Comparison** | **$F_{ST}$** | **$d_{xy}$ PC1** |
| --- | --- | --- |
| **Aur_Ari** | 0.85707 | 0.95575 |
| **Gra_Ari** | 0.89606 | 0.86182 |
| **Cal_Ari** | 0.88738 | 0.95423 |
| **CLV_Ari** | 0.8969 | 0.39278 |
| **Par_Ari** | 0.85342 | 0.94563 |
| **Lon_Ari** | 0.9046 | 0.96057 |
| **Aur_Gra** | 0.81191 | 0.85563 |
| **Aur_Cal** | 0.32407 | 0.90376 |
| **Aur_CLV** | 0.90725 | 0.46698 |
| **Aur_Par** | 0.79434 | 0.94778 |
| **Lon_Aur** | 0.40962 | 0.90381 |
| **Ari_Y** | 0.90097 | 0.94262 |
| **Aur_Y** | 0.451 | 0.90512 |
| **Gra_Y** | 0.91203 | 0.90305 |
| **Cal_Y** | 0.49255 | 0.88634 |
| **CLV_Y** | 0.90576 | 0.54264 |
| **Par_Y** | 0.86511 | 0.94318 |
| **Lon_Y** | 0.59046 | 0.88521 |
| **Cal_Gra** | 0.89043 | 0.88319 |
| **CLV_Gra** | 0.91181 | 0.32045 |
| **Par_Gra** | 0.89579 | 0.86769 |
| **Lon_Gra** | 0.88549 | 0.89344 |
| **Cal_Par** | 0.83543 | 0.94124 |
| **Lon_Cal** | 0.3786 | 0.87337 |
| **Cal_CLV** | 0.90548 | 0.52607 |
| **Par_CLV** | 0.91991 | 0.45233 |
| **Lon_CLV** | 0.90738 | 0.51482 |
| **Lon_Par** | 0.84197 | 0.94507 |
| **R_Ari** | 0.90783 | 0.94574 |
| **R_Aur** | 0.51788 | 0.89861 |
| **R_Y** | 0.51423 | 0.84798 |
| **R_Gra** | 0.91936 | 0.90695 |
| **R_Cal** | 0.62051 | 0.88316 |
| **R_CLV** | 0.90758 | 0.55149 |
| **R_Par** | 0.87615 | 0.93825 |
| **R_Lon** | 0.66912 | 0.88264 |

**Table S4 continued.**
| **Taxon** | **$\pi$ PC1** |
| --- | --- |
| **Ari** | 0.9251 |
| **Aur** | 0.86508 |
| **Cal** | 0.96734 |
| **CLV** | 0.92017 |
| **Gra** | 0.77181 |
| **Lon** | 0.95166 |
| **Par** | 0.91044 |
| **R** | 0.97282 |
| **Y** | 0.97255 |

**Table S5.** Details for the linear regressions presented in **Figure 5** of the main text.

| **Variables** | **Pearson's r** | **Regression equation** | ***p*** |
| --- | --- | --- | --- |
| $d_{xy}$ & $\pi$ | -0.93 | $y = -60.5x + 1.33$ | $< 0.001$ |
| $\pi$ & gene count | 0.59 | $y = 3.9x - 0.85$ | 0.130 |
| $\pi$ & cM/mbp | -0.79 | $y = -5.4x + 0.46$ | 0.020 |
| $F_{ST}$ & $\pi$ | -0.89 | $y = -53.3x - 0.13$ | 0.003 |
| $F_{ST}$ & gene count | 0.81 | $y = 46.5x + 0.04$ | 0.016 |
| $F_{ST}$ & cM/mbp | -0.73 | $y = -25.5x + 0.01$ | 0.041 |

**Table S6.** Parameters used in SLiM to simulate genomic landscape evolution. (*N*=10,000).

| **Class** | **Mean Ns of deleterious mutations** | **Percent of deleterious mutations** | **Mean Ns of beneficial mutations** | **Percent of beneficial mutations** | **Migration rate (Nm)** |
| --- | --- | --- | --- | --- | --- |
| Neutral | - | - | - | - | - |
| BGS | -10 | 5% | - | - | - |
|  | -10 | 10% | - | - | - |
|  | -100 | 5% | - | - | - |
|  | -100 | 10% | - | - | - |
| BDMI | -10 | 5% | - | - | 0.1 |
|  | -10 | 10% | - | - | 1 |
|  | -10 | 5% | - | - | 0.1 |
|  | -10 | 10% | - | - | 1 |
|  | -100 | 5% | - | - | 0.1 |
|  | -100 | 10% | - | - | 1 |
|  | -100 | 5% | - | - | 0.1 |
|  | -100 | 10% | - | - | 1 |
| Positive | - | - | 100 | 0.1% | - |
|  | - | - | 100 | 0.5% | - |
| Local adaptation | - | - | 100 | 0.1% | 0.1 |
|  | - | - | 100 | 0.1% | 1 |
|  | - | - | 100 | 0.5% | 0.1 |
|  | - | - | 100 | 0.5% | 1 |
| BGS & positive | -10 | 5% | 100 | 0.1% | - |
|  | -10 | 5% | 100 | 0.5% | - |
|  | -10 | 10% | 100 | 0.1% | - |
|  | -10 | 10% | 100 | 0.5% | - |
|  | -100 | 5% | 100 | 0.1% | - |
|  | -100 | 5% | 100 | 0.5% | - |
|  | -100 | 10% | 100 | 0.1% | - |
|  | -100 | 10% | 100 | 0.5% | - |

**Table S7.** Genome-wide estimates of admixture across the bush monkeyflower radiation. Measures of Patterson’s *D* and the admixture proportion, *f*, are given for all 48 possible pairs of four-taxon tests, with *M. clevelandii* as the outgroup for each test. All values of Patterson’s *D* are statistically significant (P < 0.0001) based on a block jackknife approach.

| **P1** | **P2** | **P3** | **Patterson's D** | **Admixture proportion** |
| --- | --- | --- | --- | --- |
| Aur | Cal | Ari | 0.0808 | 0.0209 |
| Lon | Cal | Ari | 0.0233 | 0.0055 |
| Aur | Lon | Ari | 0.0633 | 0.0155 |
| Aur | R | Ari | 0.1360 | 0.0389 |
| Cal | R | Ari | 0.0651 | 0.0184 |
| Lon | R | Ari | 0.0883 | 0.0238 |
| Aur | Y | Ari | 0.1595 | 0.0474 |
| Cal | Y | Ari | 0.0949 | 0.0270 |
| Lon | Y | Ari | 0.1177 | 0.0324 |
| R | Y | Ari | 0.0371 | 0.0088 |
| R | Cal | Aur | 0.0831 | 0.0903 |
| Y | Cal | Aur | 0.0976 | 0.1034 |
| R | Lon | Aur | 0.0866 | 0.0909 |
| Y | Lon | Aur | 0.1009 | 0.1041 |
| Y | R | Aur | 0.0188 | 0.0145 |
| Ari | Par | Gra | 0.1053 | 0.0243 |
| Ari | Aur | Gra | 0.1821 | 0.0477 |
| Cal | Aur | Gra | 0.0648 | 0.0096 |
| Lon | Aur | Gra | 0.0936 | 0.0141 |
| Y | Aur | Gra | 0.1392 | 0.0218 |
| R | Aur | Gra | 0.1471 | 0.0222 |
| Par | Aur | Gra | 0.1072 | 0.0240 |
| Ari | Cal | Gra | 0.1523 | 0.0384 |
| Lon | Cal | Gra | 0.0344 | 0.0045 |
| Y | Cal | Gra | 0.0886 | 0.0123 |
| R | Cal | Gra | 0.0943 | 0.0126 |
| Par | Cal | Gra | 0.0665 | 0.0145 |
| Ari | Lon | Gra | 0.1371 | 0.0341 |
| Par | Lon | Gra | 0.0478 | 0.0101 |
| Y | Lon | Gra | 0.0589 | 0.0078 |
| R | Lon | Gra | 0.0650 | 0.0082 |
| Ari | R | Gra | 0.1072 | 0.0261 |
| Par | R | Gra | 0.0099 | 0.0019 |
| Ari | Y | Gra | 0.1103 | 0.0265 |
| Par | Y | Gra | 0.0127 | 0.0023 |
| R | Y | Gra | 0.0059 | 0.0004 |
| R | Y | Cal | 0.0149 | 0.0438 |
| Aur | Cal | Par | 0.0415 | 0.0182 |
| R | Cal | Par | 0.0104 | 0.0055 |
| Y | Cal | Par | 0.0176 | 0.0077 |
| Aur | Lon | Par | 0.0456 | 0.0207 |
| Cal | Lon | Par | 0.0049 | 0.0025 |
| R | Lon | Par | 0.0151 | 0.0080 |
| Y | Lon | Par | 0.0222 | 0.0102 |
| Aur | R | Par | 0.0311 | 0.0128 |
| Y | R | Par | 0.0090 | 0.0022 |
| Aur | Y | Par | 0.0240 | 0.0106 |
| R | Y | Lon | 0.0001 | -0.0018 |

**Table S8.**
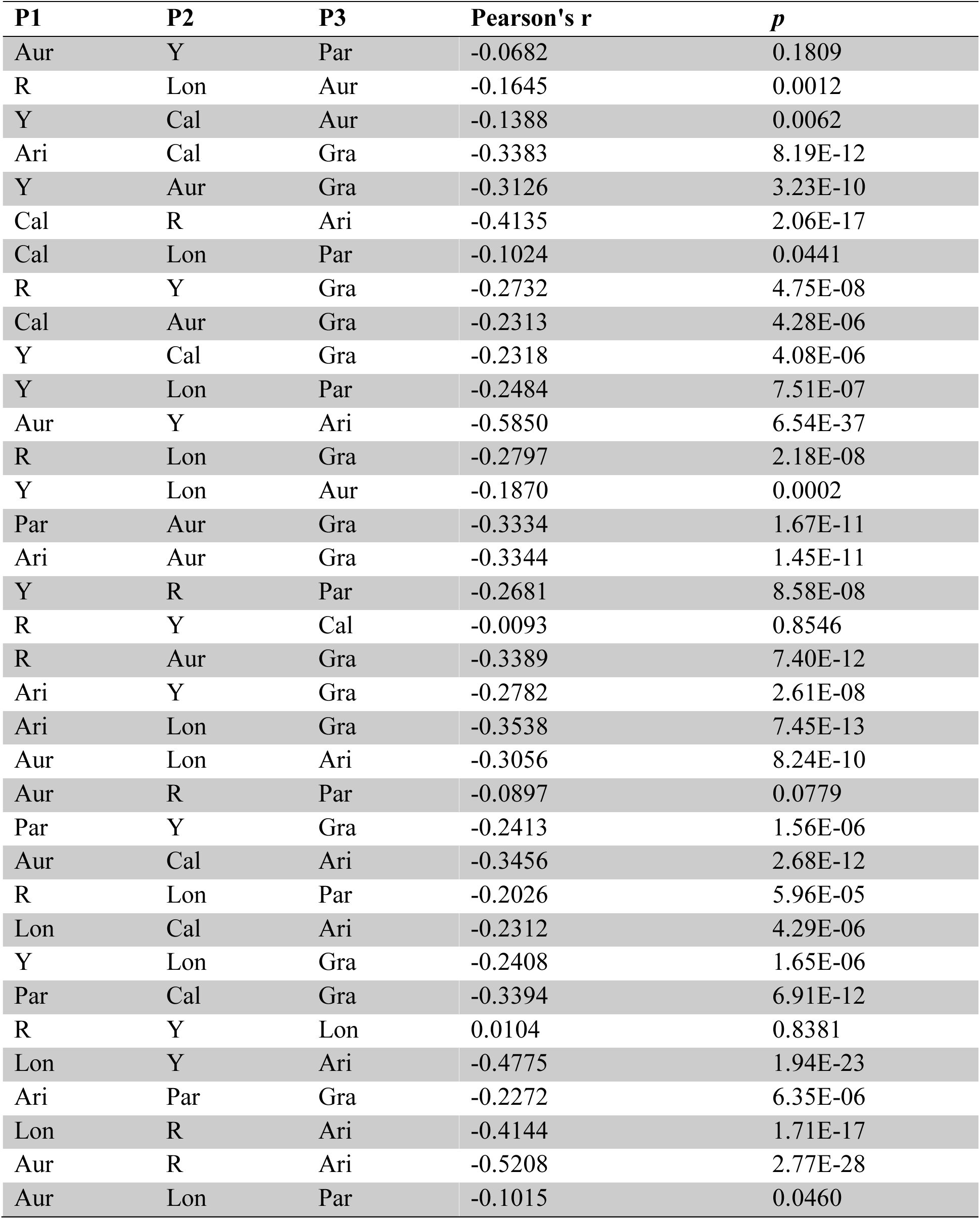
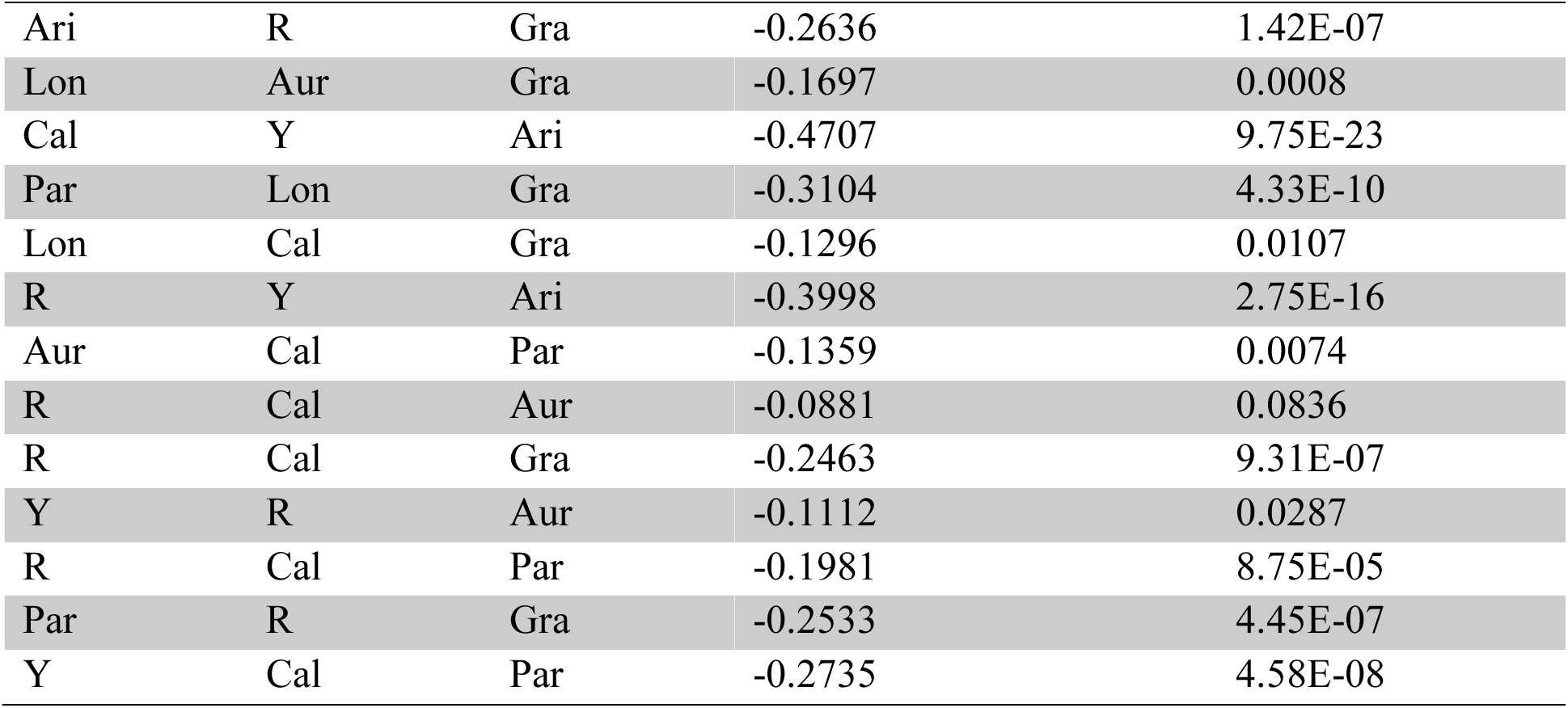
Correlation coefficients between the window-based admixture proportion (f_d_) and PC1 F_ST_. Pearson’s r was calculated between PC1 *F_ST_* and *f_d_* for each of the 48 four-taxon tests, measured in 500kb non-overlapping windows. *M. clevelandii* is the outgroup for each test.

| P1 | P2 | P3 | Pearson's $r$ | $p$ |
| --- | --- | --- | --- | --- |
| Aur | Y | Par | -0.0682 | 0.1809 |
| R | Lon | Aur | -0.1645 | 0.0012 |
| Y | Cal | Aur | -0.1388 | 0.0062 |
| Ari | Cal | Gra | -0.3383 | 8.19E-12 |
| Y | Aur | Gra | -0.3126 | 3.23E-10 |
| Cal | R | Ari | -0.4135 | 2.06E-17 |
| Cal | Lon | Par | -0.1024 | 0.0441 |
| R | Y | Gra | -0.2732 | 4.75E-08 |
| Cal | Aur | Gra | -0.2313 | 4.28E-06 |
| Y | Cal | Gra | -0.2318 | 4.08E-06 |
| Y | Lon | Par | -0.2484 | 7.51E-07 |
| Aur | Y | Ari | -0.5850 | 6.54E-37 |
| R | Lon | Gra | -0.2797 | 2.18E-08 |
| Y | Lon | Aur | -0.1870 | 0.0002 |
| Par | Aur | Gra | -0.3334 | 1.67E-11 |
| Ari | Aur | Gra | -0.3344 | 1.45E-11 |
| Y | R | Par | -0.2681 | 8.58E-08 |
| R | Y | Cal | -0.0093 | 0.8546 |
| R | Aur | Gra | -0.3389 | 7.40E-12 |
| Ari | Y | Gra | -0.2782 | 2.61E-08 |
| Ari | Lon | Gra | -0.3538 | 7.45E-13 |
| Aur | Lon | Ari | -0.3056 | 8.24E-10 |
| Aur | R | Par | -0.0897 | 0.0779 |
| Par | Y | Gra | -0.2413 | 1.56E-06 |
| Aur | Cal | Ari | -0.3456 | 2.68E-12 |
| R | Lon | Par | -0.2026 | 5.96E-05 |
| Lon | Cal | Ari | -0.2312 | 4.29E-06 |
| Y | Lon | Gra | -0.2408 | 1.65E-06 |
| Par | Cal | Gra | -0.3394 | 6.91E-12 |
| R | Y | Lon | 0.0104 | 0.8381 |
| Lon | Y | Ari | -0.4775 | 1.94E-23 |
| Ari | Par | Gra | -0.2272 | 6.35E-06 |
| Lon | R | Ari | -0.4144 | 1.71E-17 |
| Aur | R | Ari | -0.5208 | 2.77E-28 |
| Aur | Lon | Par | -0.1015 | 0.0460 |
| Ari | R | Gra | -0.2636 | 1.42E-07 |
| Lon | Aur | Gra | -0.1697 | 0.0008 |
| Cal | Y | Ari | -0.4707 | 9.75E-23 |
| Par | Lon | Gra | -0.3104 | 4.33E-10 |
| Lon | Cal | Gra | -0.1296 | 0.0107 |
| R | Y | Ari | -0.3998 | 2.75E-16 |
| Aur | Cal | Par | -0.1359 | 0.0074 |
| R | Cal | Aur | -0.0881 | 0.0836 |
| R | Cal | Gra | -0.2463 | 9.31E-07 |
| Y | R | Aur | -0.1112 | 0.0287 |
| R | Cal | Par | -0.1981 | 8.75E-05 |
| Par | R | Gra | -0.2533 | 4.45E-07 |
| Y | Cal | Par | -0.2735 | 4.58E-08 |

## Supplementary figures

**Figure. S1.**
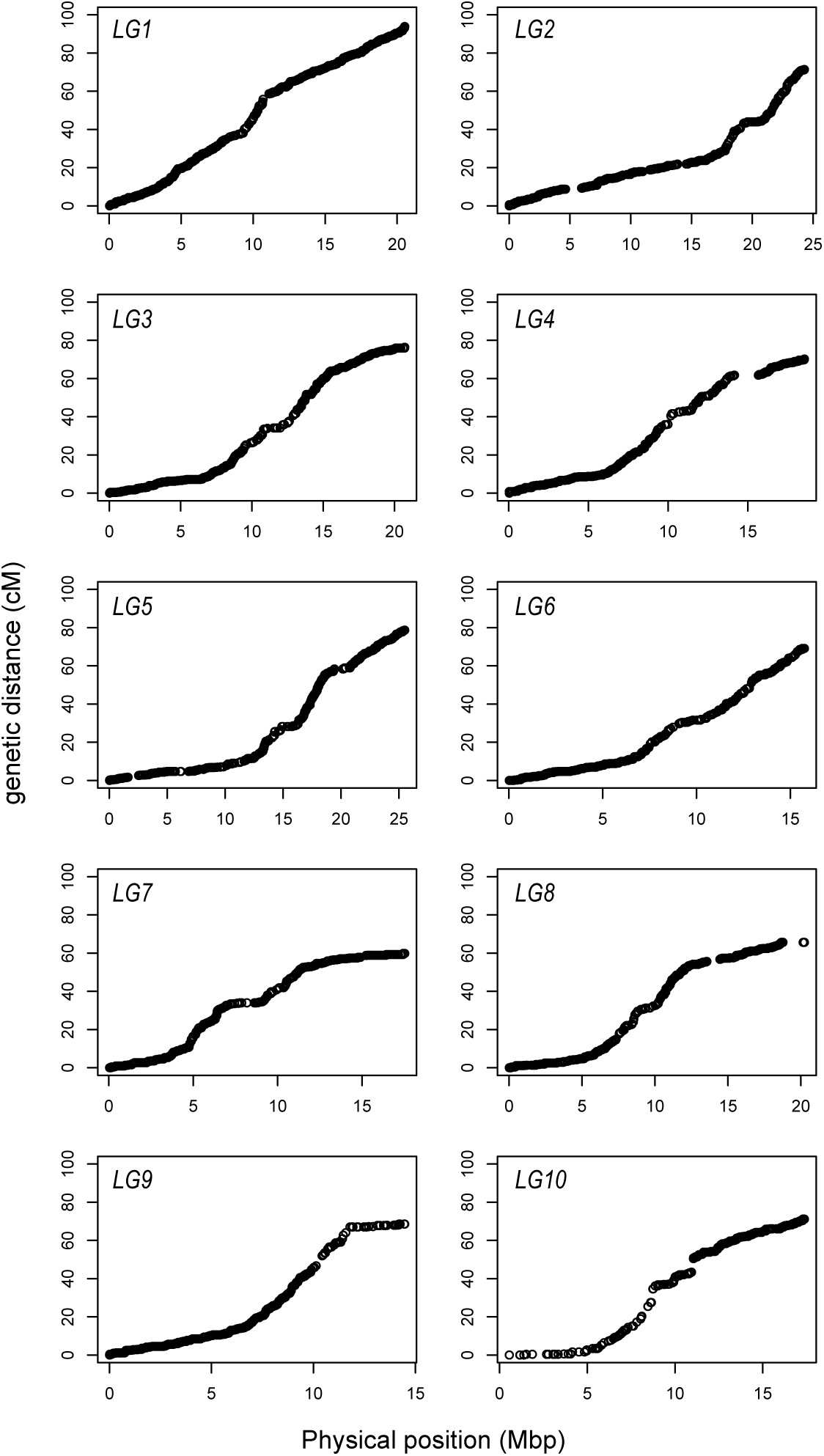
Map distance (cM/Mbp) vs. physical distance across the 10 linkage groups. Recombination for each marker is estimated relative to the start of the linkage group and plotted at its physical location on each chromosome in the reference assembly.

**Figure S2.**
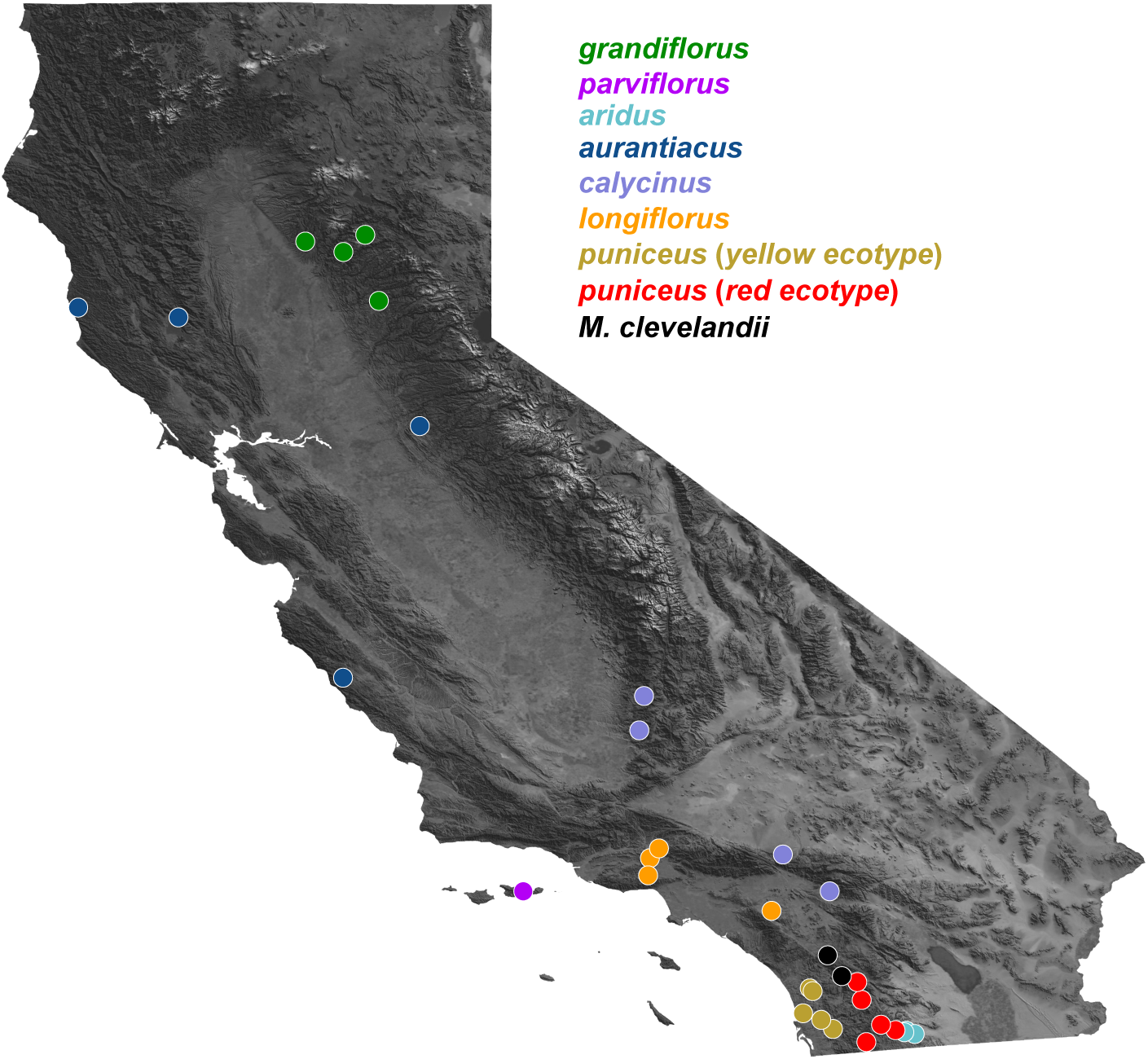
Geographic distribution of sampling locations for each sample sequenced in this study. Detailed position information for each population can be found in Table S3.

**Figure S3.**
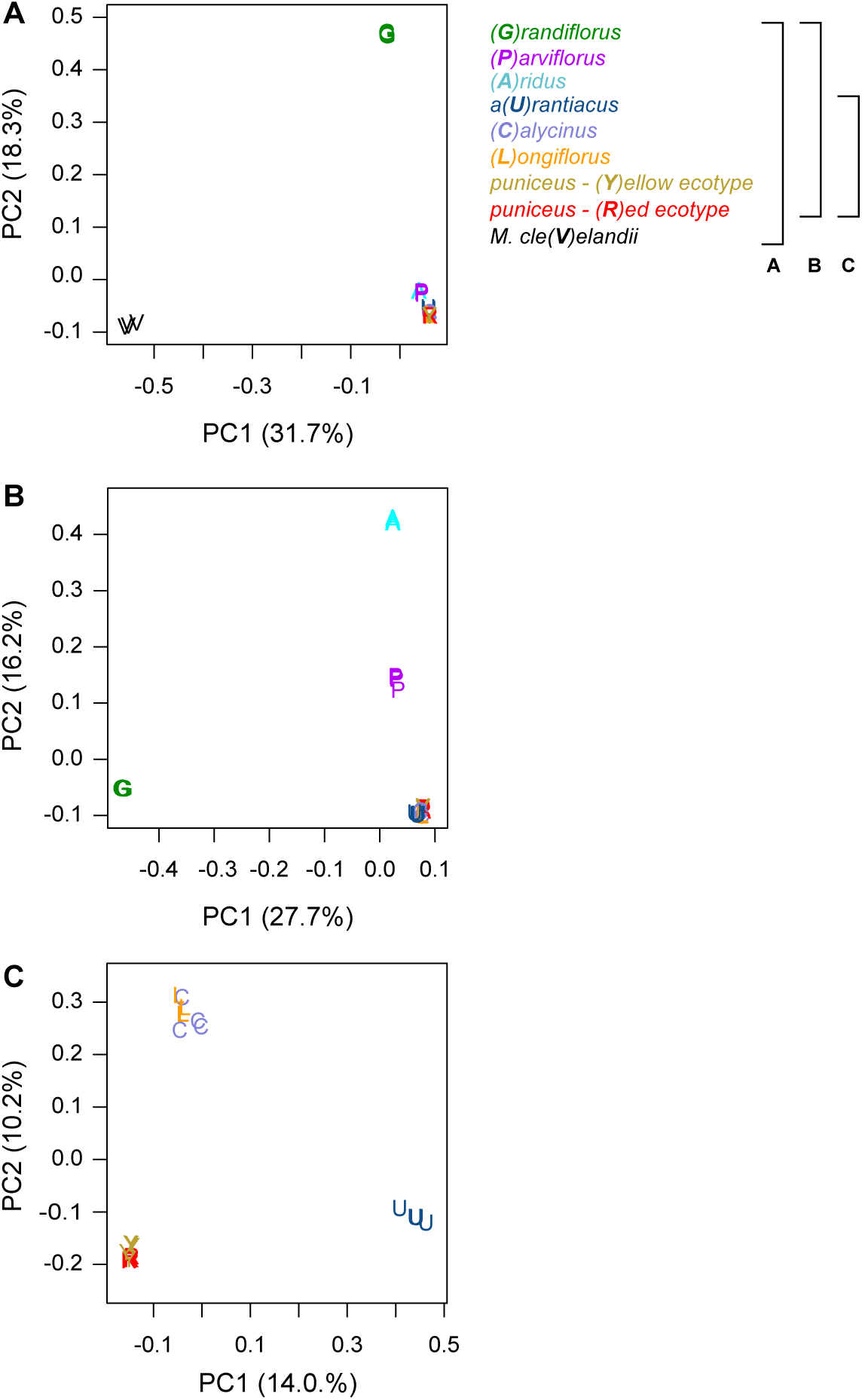
Genome-wide Principal Components Analysis (PCA). Each plot is a separate PCA performed using different sets of taxa. The legend to the right describes the set of taxa included in each analysis, with the capital letter in parentheses and the color representing the specific taxon. A) All taxa; B) all subspecies of *M. aurantiacus*, but excluding *M. clevelandii*; C) only subspecies *aurantiacus*, *longiflorus*, *calycinus*, and the red and yellow ecotypes of subspecies *puniceus*. The percent variation explained by each principal component is given in parentheses.

**Figure S4.**
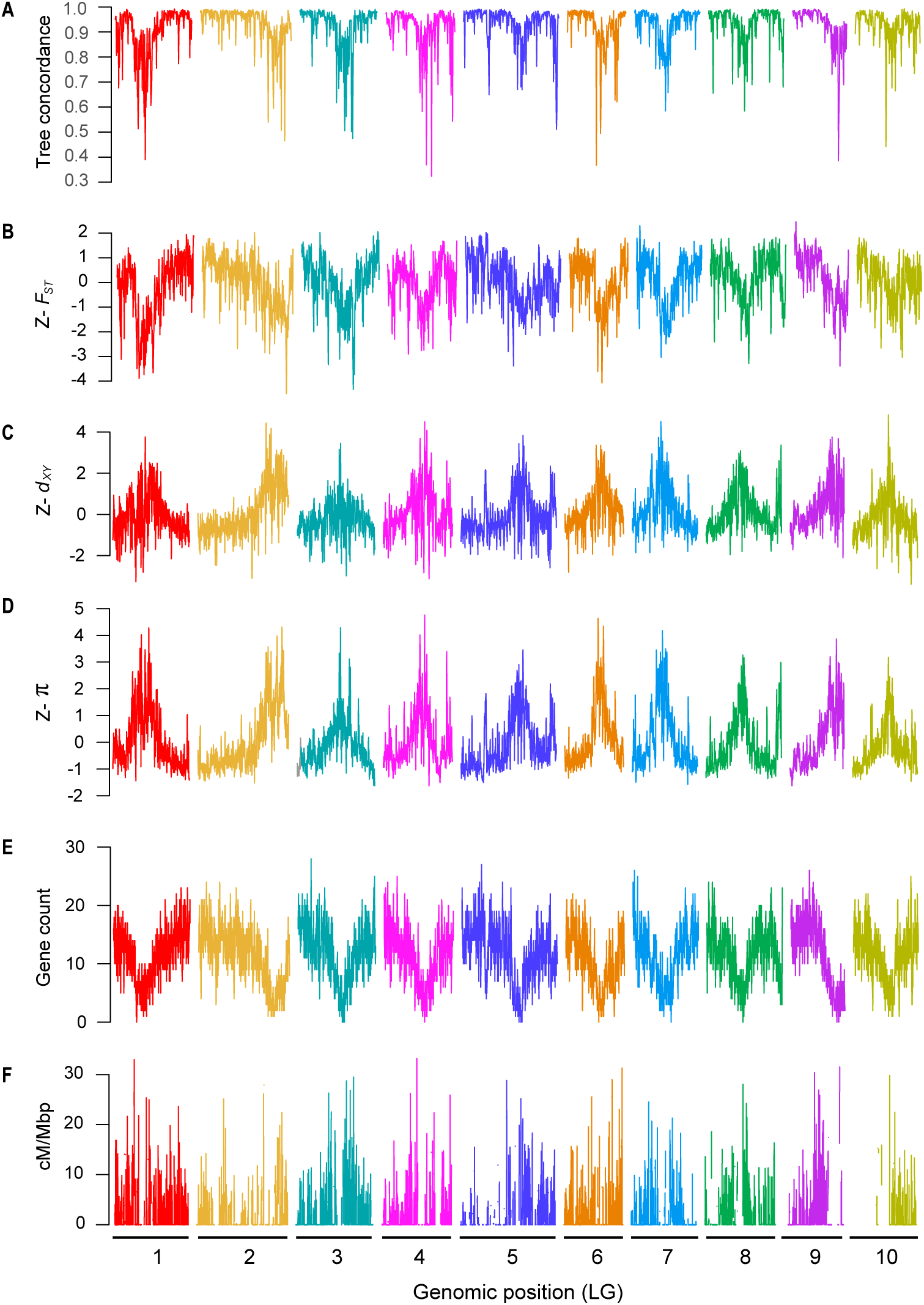
Common patterns of genome-wide variation mirror variation in the local properties of the genome. Plots are the same as in Fig. 2 of the main text, but for 100 kb windows (step size 10 kbp). A) Tree concordance; B-D) Z-transformed PC1 for *F*_ST_, *d*_xy_ and π, respectively; E) gene count; and F) recombination rate (cM/Mbp) are plotted across the 10 monkeyflower LGs.

**Figure S5.**
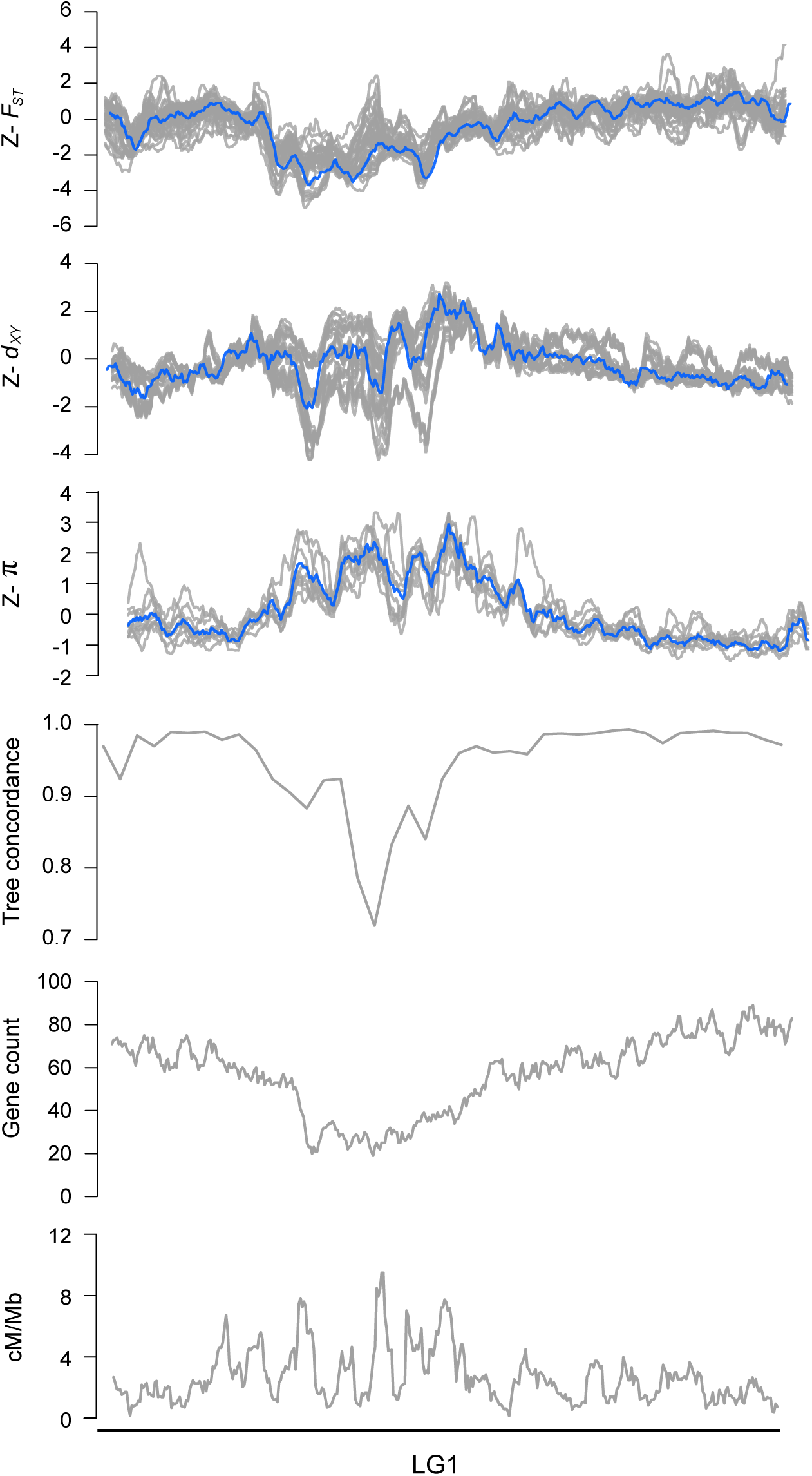
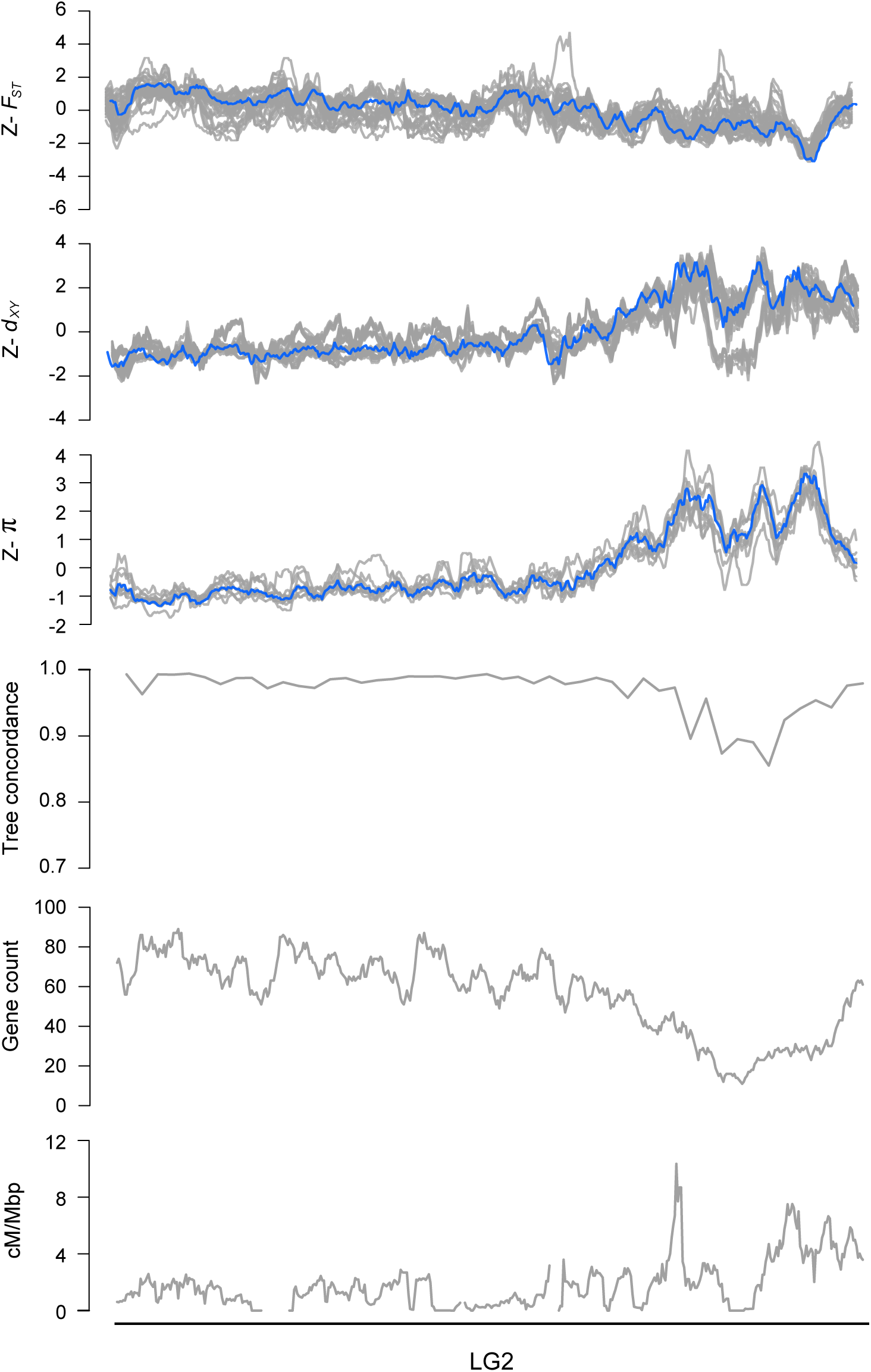
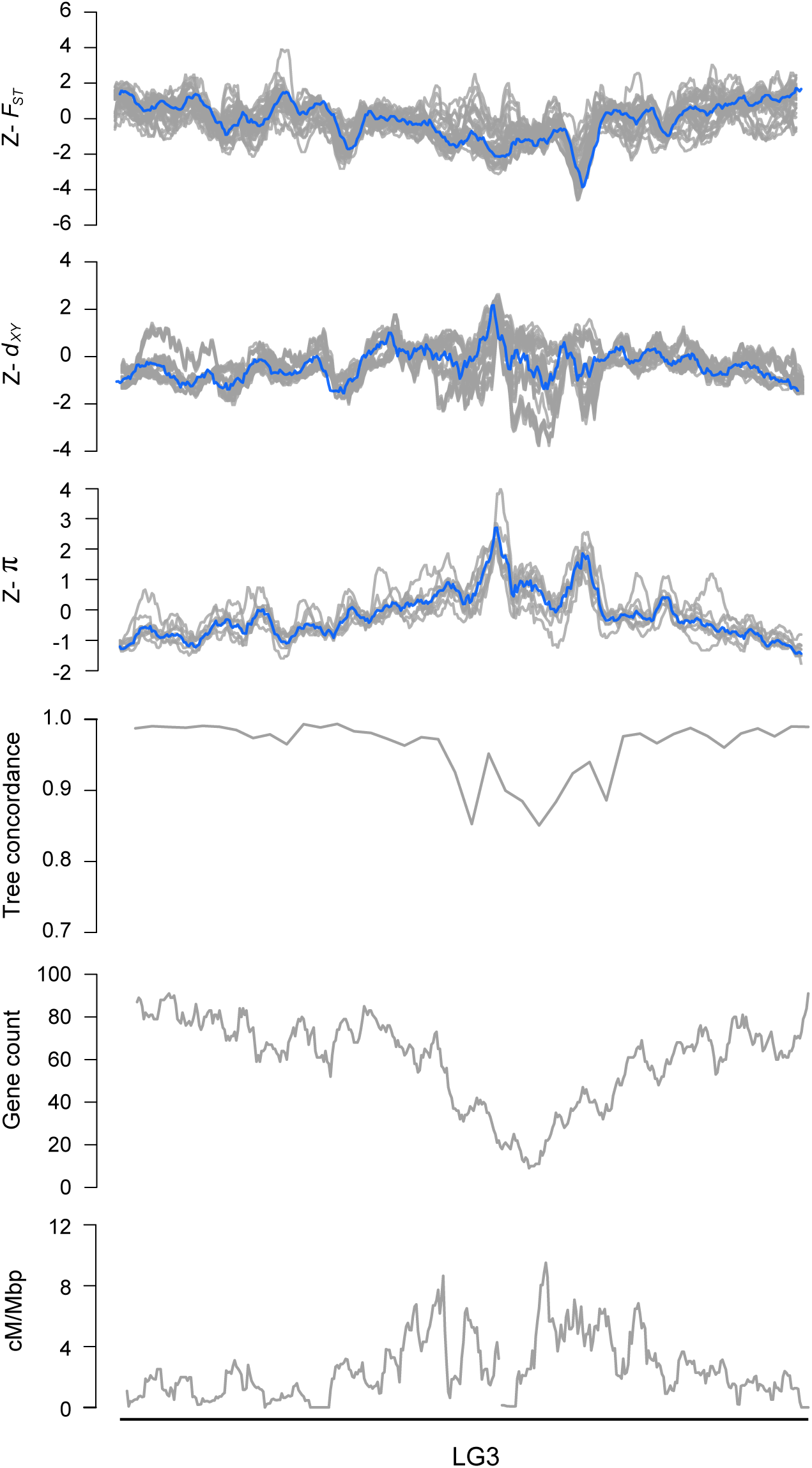
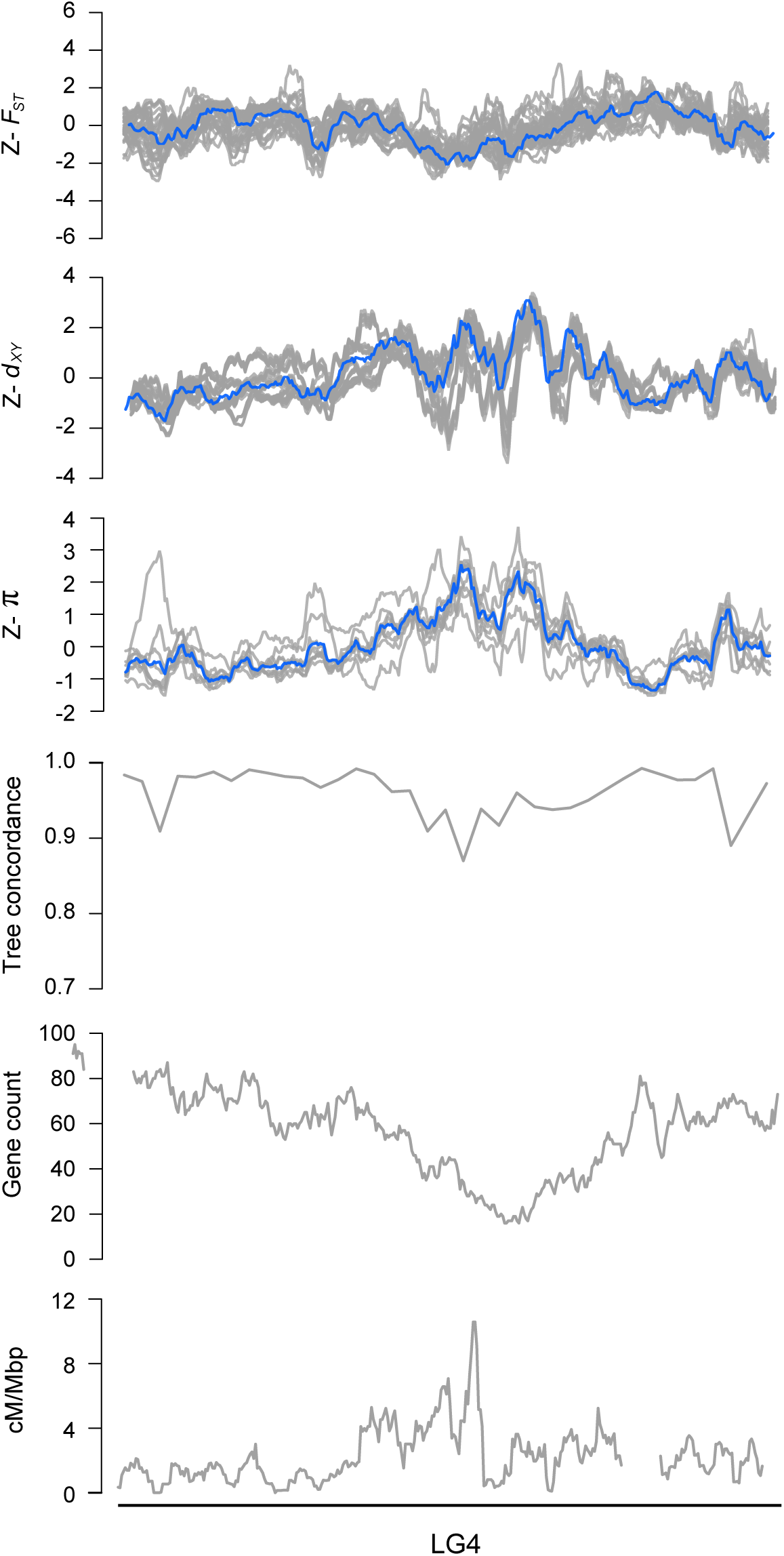
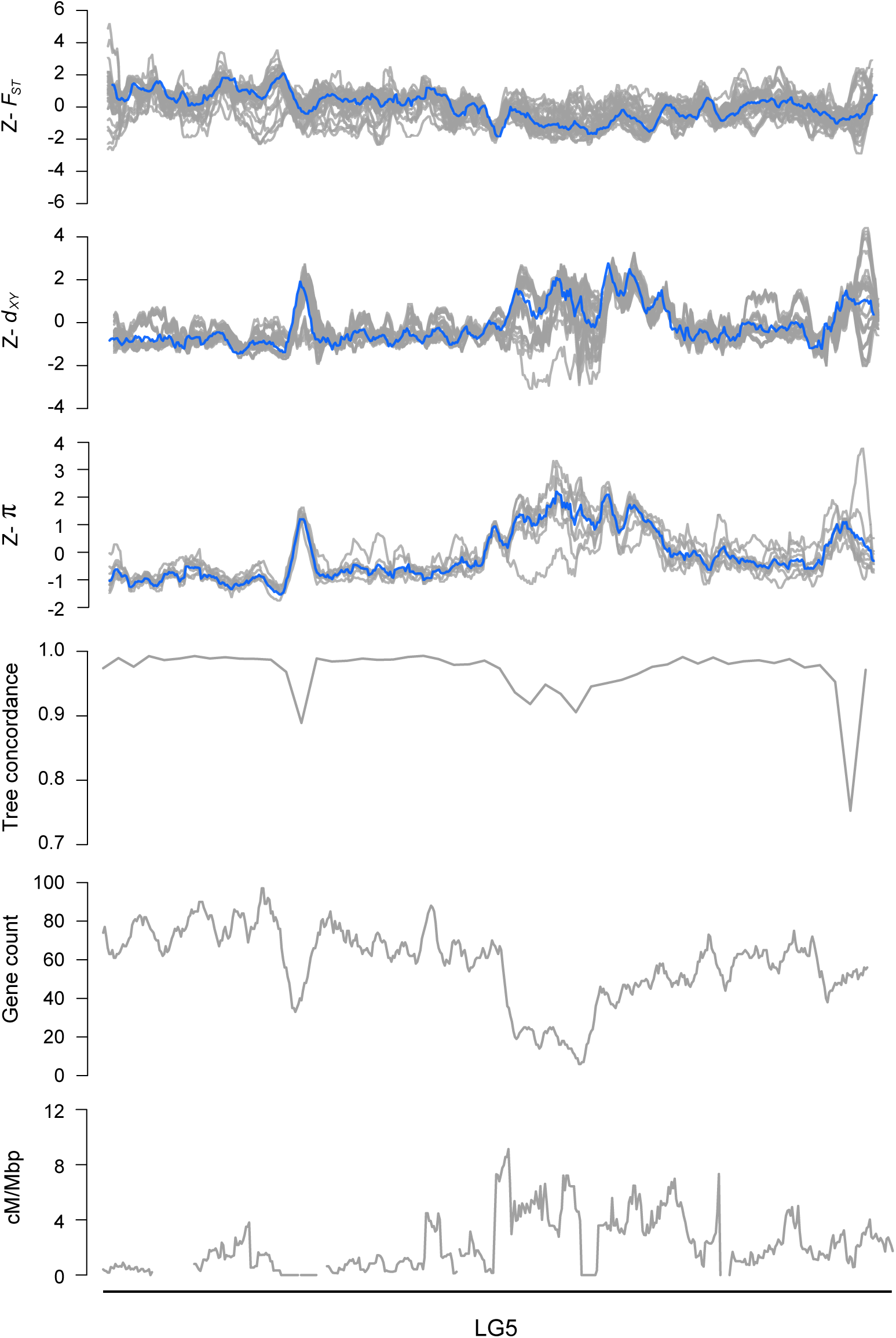
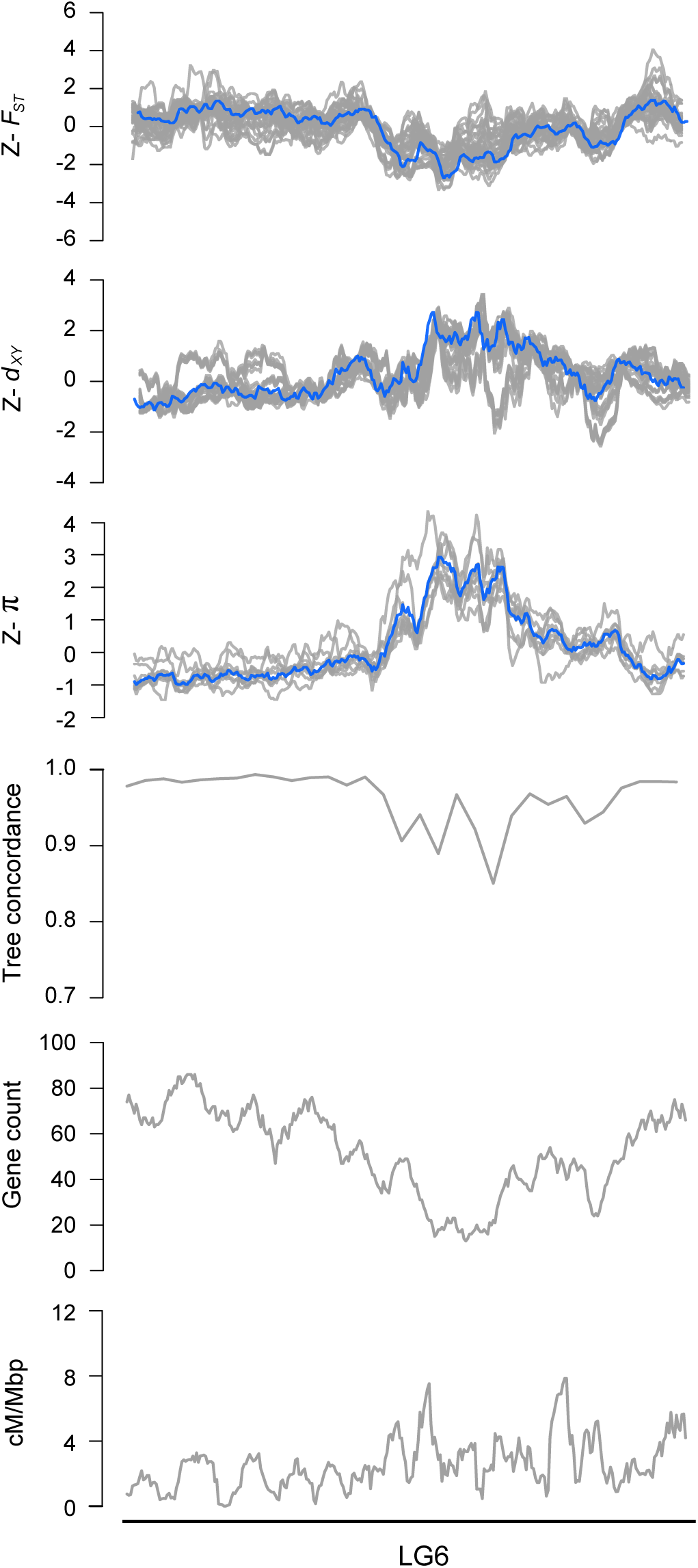
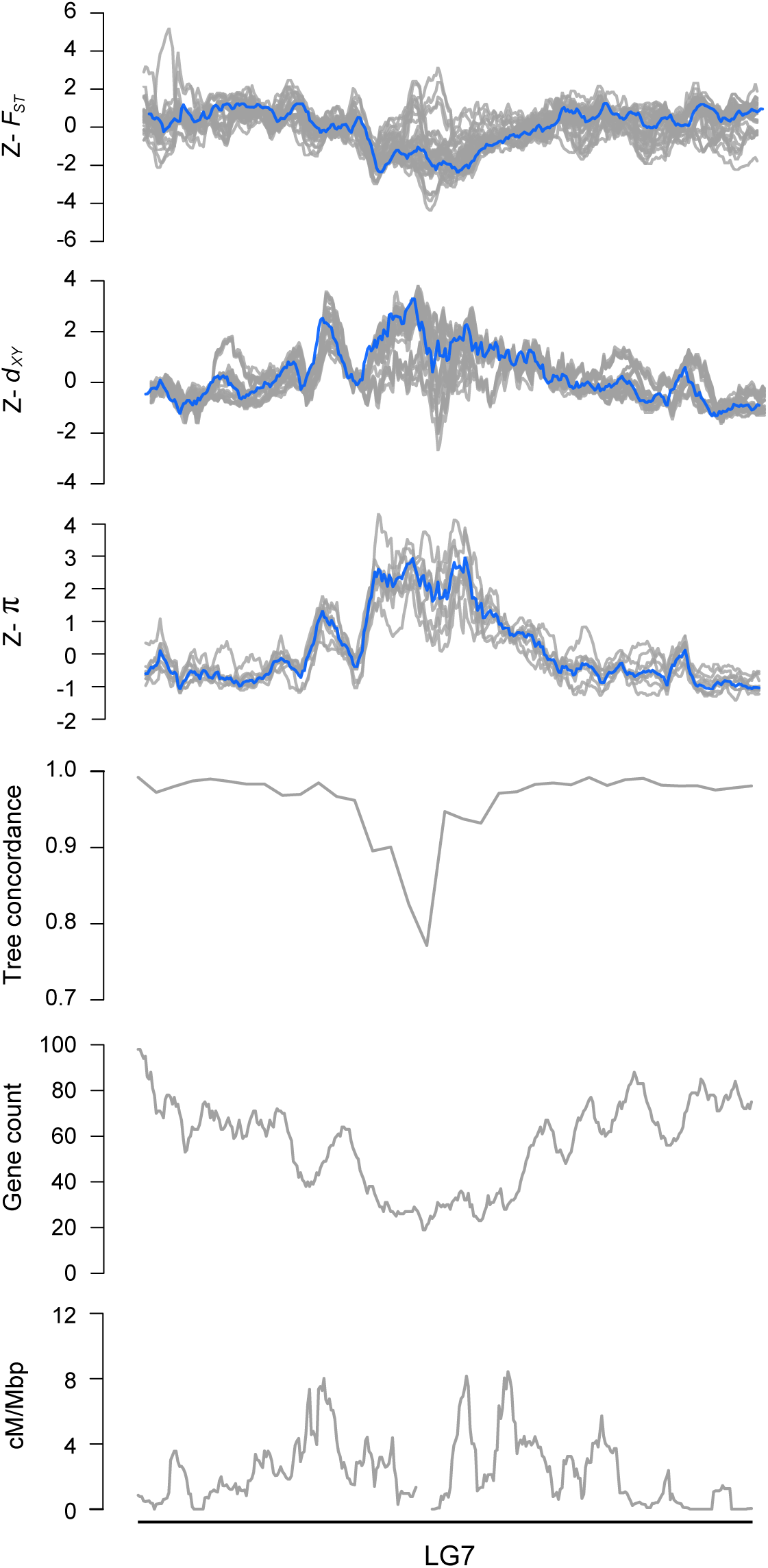
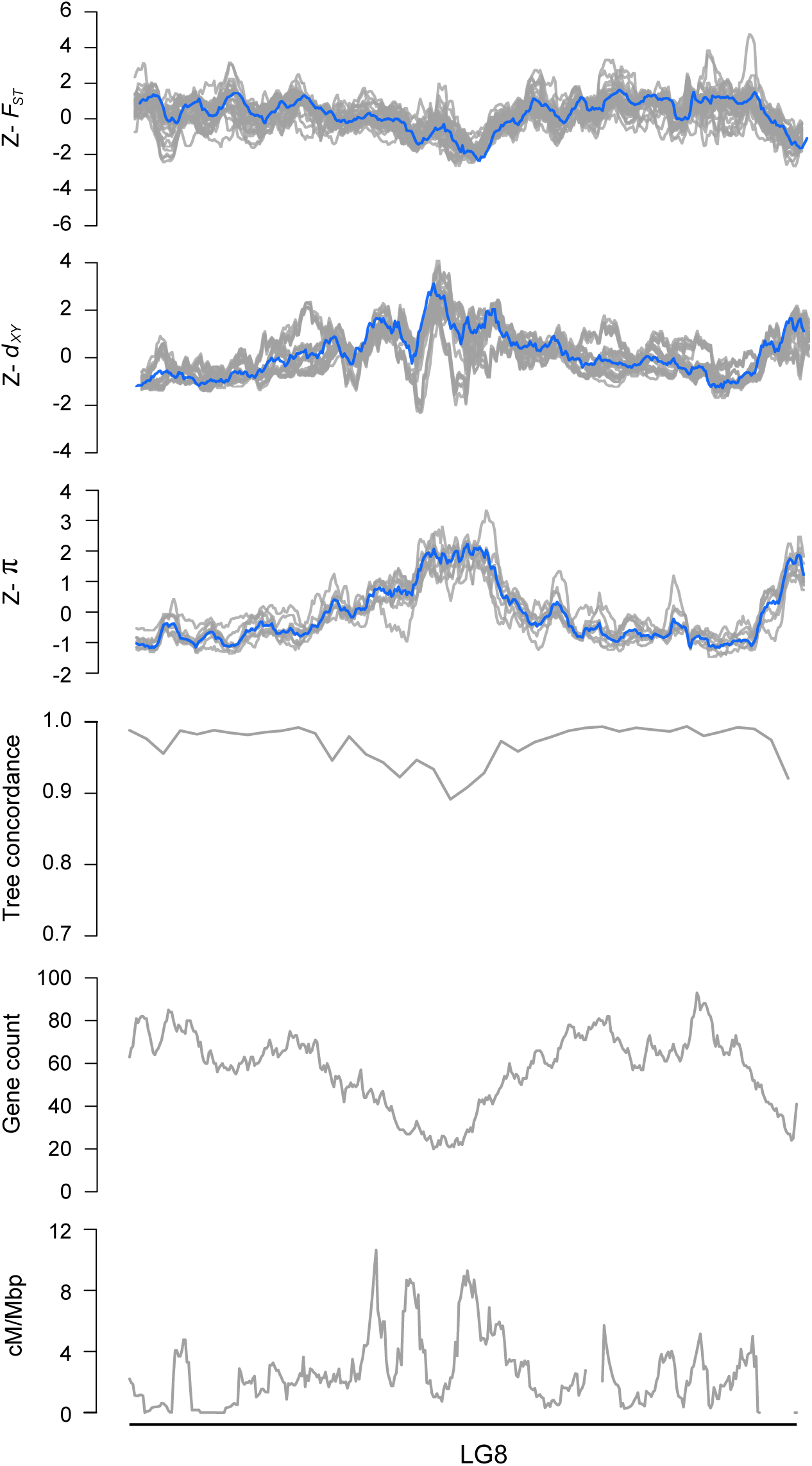
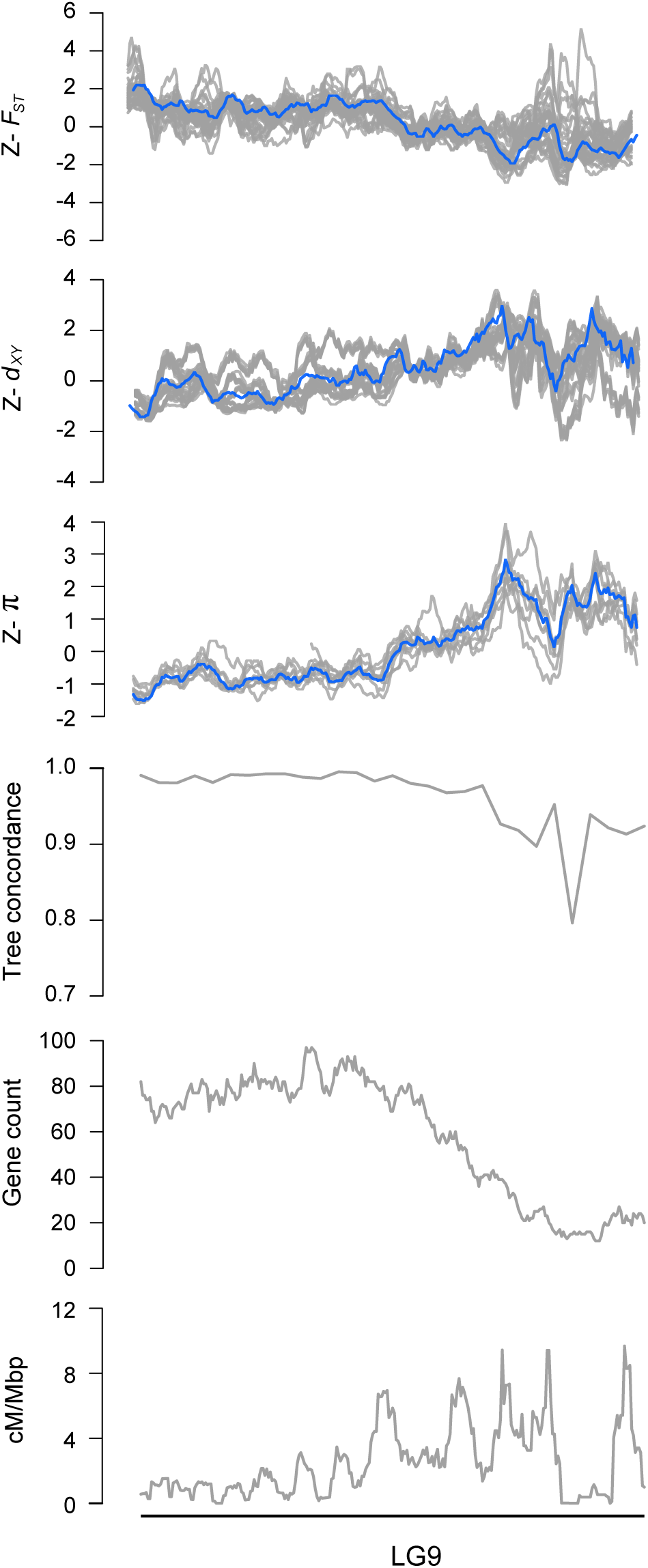
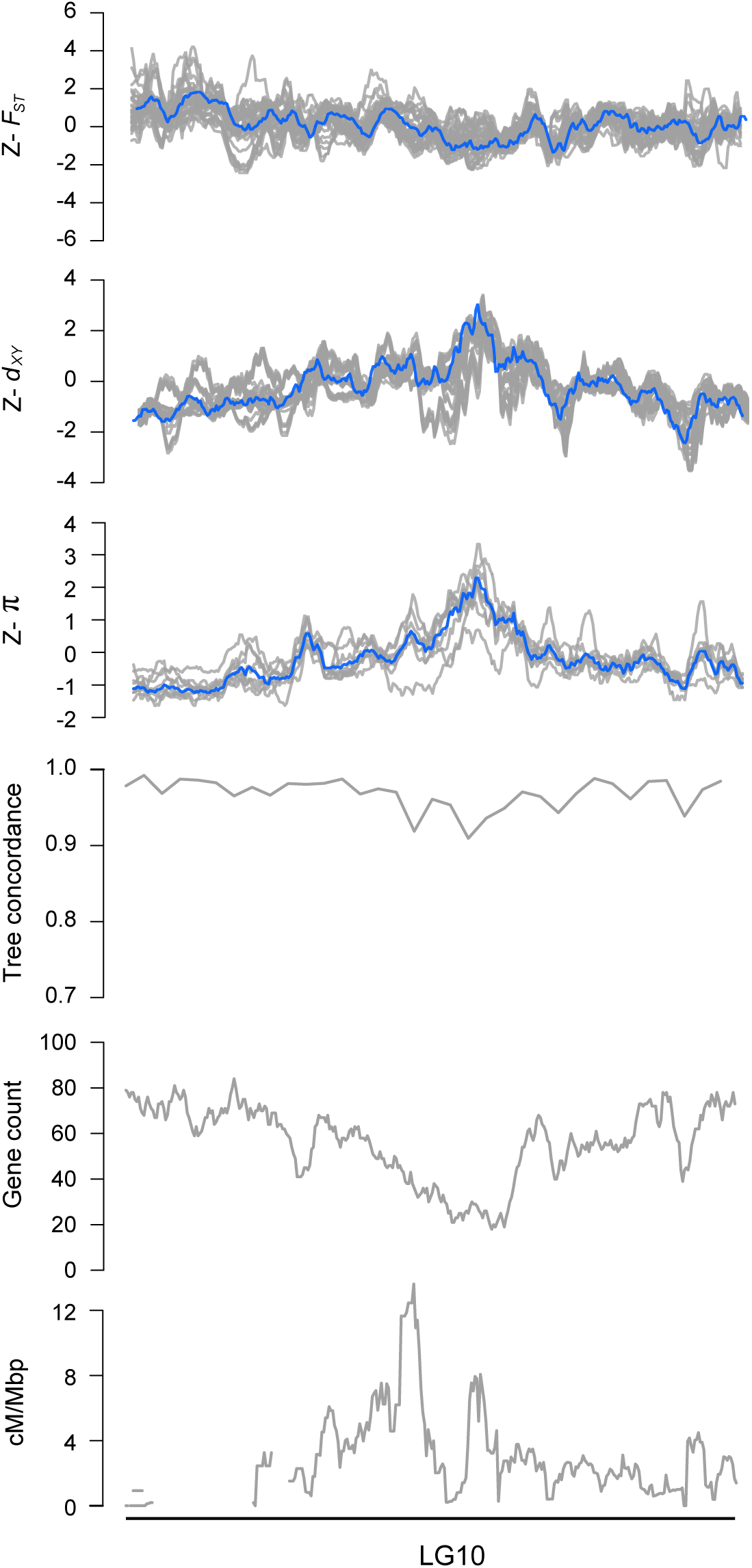
Patterns of variation plotted across each bush monkeyflower linkage group. Z-transformed *F_ST_*, *d_xy_*, and π in overlapping 500 kb windows (step size = 50 kbp). The gray lines are z-transformed scores for each of the 36 pairwise comparisons (*F_ST_*and *d_xy_*) or nine taxa (π), and the blue line is the z-transformed score for the first principal component (PC1). Estimates of tree concordance, gene count and recombination rate (cM/Mbp) are also shown.

**Figure S6.**
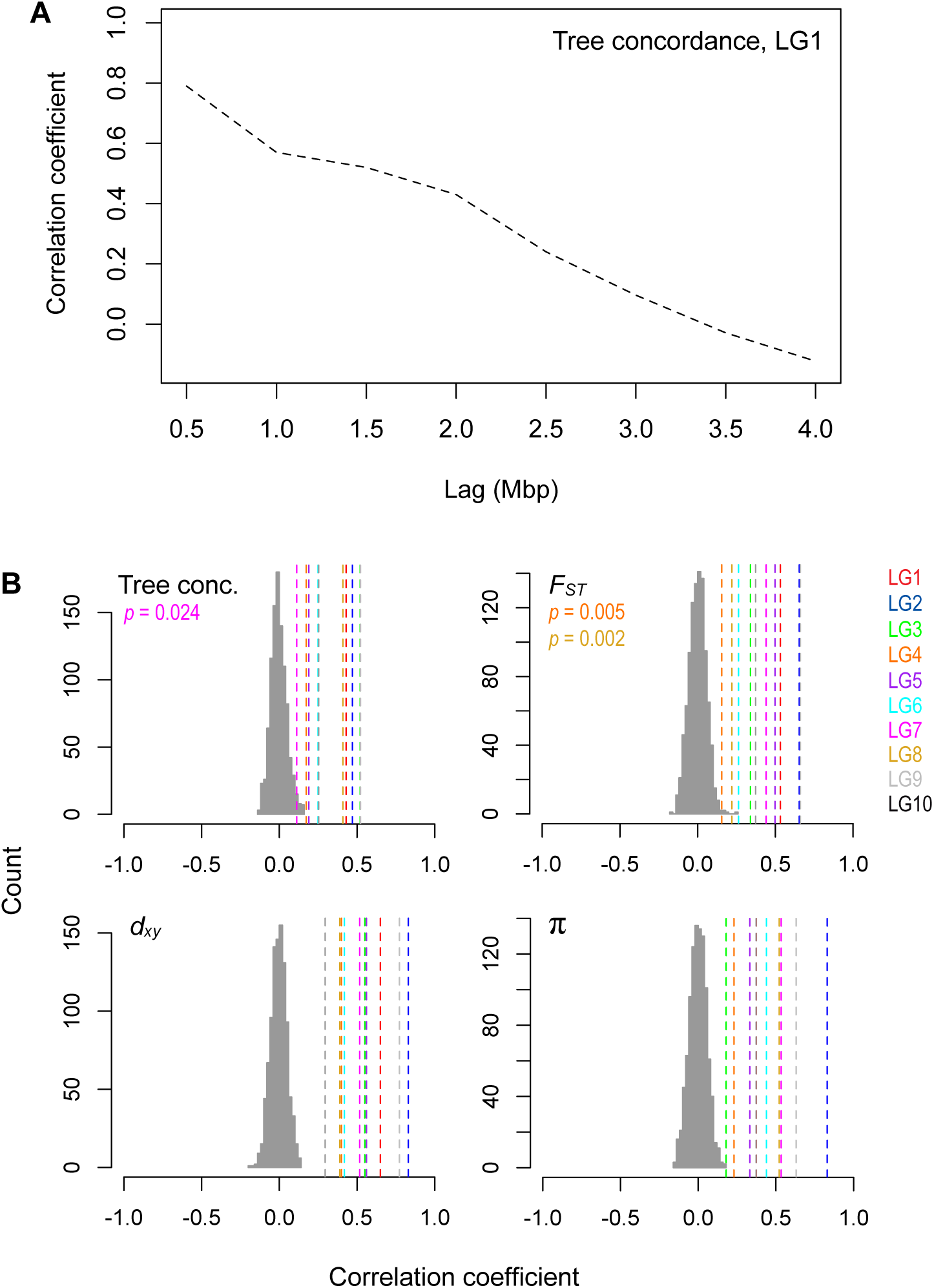
Patterns of variation are non-randomly distributed across the genome. A) Strong autocorrelation of tree concordance scores on LG1 over a Mbp scale. B) Levels of tree concordance, *F_ST_*, *d*_xy_, and π all show significant autocorrelation at the 2 Mbp scale. The dashed vertical lines show the observed autocorrelation coefficients for each LG with a 2 Mbp lag. The histogram shows the null distribution of autocorrelation coefficients (same lag) generated from 1000 random permutations of the genome-wide values. The observed data are significant at *p* = 0.001 unless stated otherwise.

**Figure S7.**
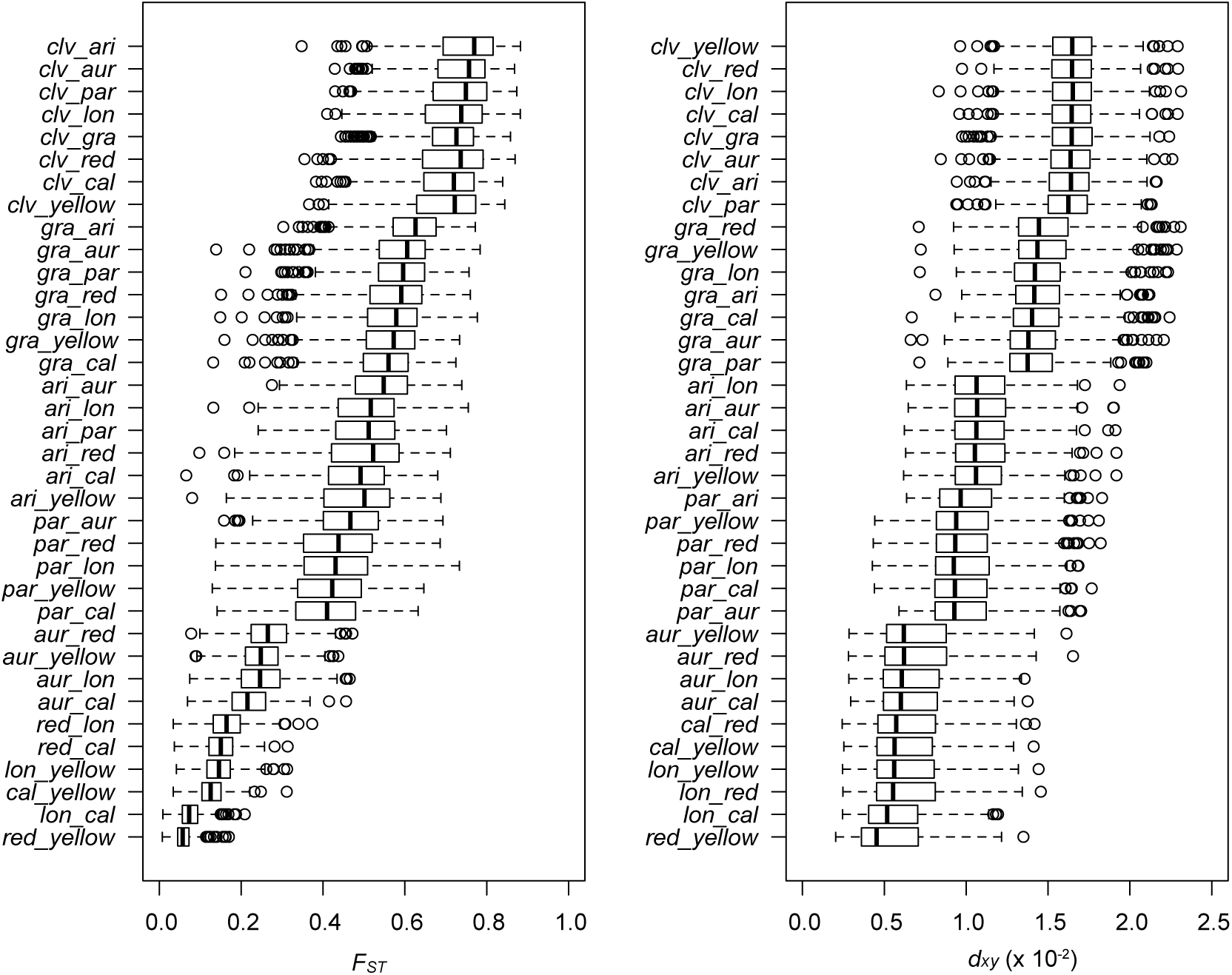
Patterns of differentiation and divergence for all 36 pairs of taxa. Box plots for each of the 36 pairwise taxonomic comparisons reveal the range of variation in *F_ST_* and *d*_xy_ across the radiation. Moreover, the data show extensive variance among genomic windows within each comparison. Vertical black lines indicate the median, boxes represent the lower and upper quartiles, and whiskers extend to 1.5 times the interquartile range. Taxon abbreviations: *cal, calycinus; lon, longiflorus; aur, aurantiacus; par, parviflorus; ari, aridus; gra, grandiflorus; clv, M. clevelandii*.

**Figure S8.**
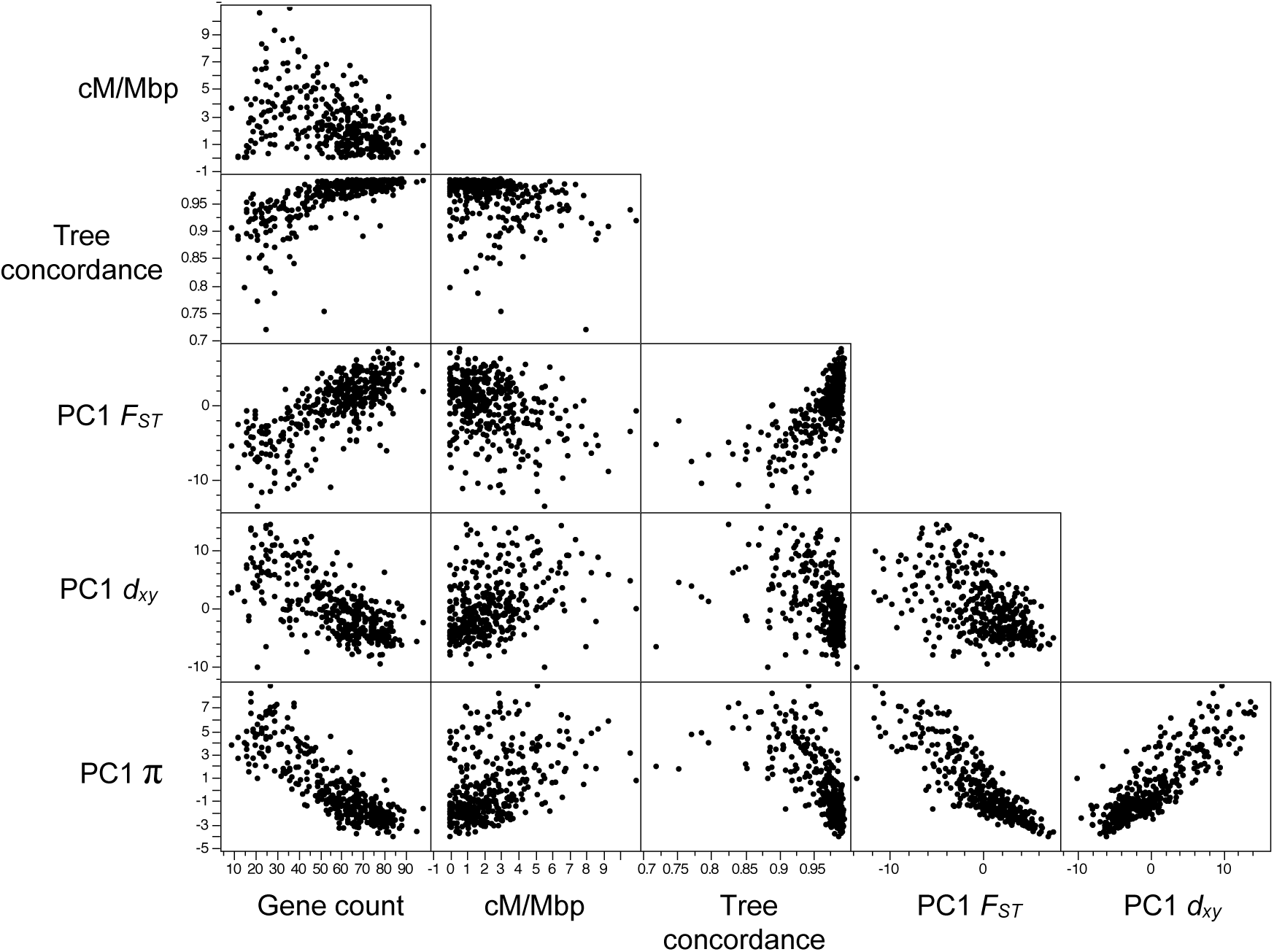
Bivariate plots among measures of variation and genomic features across 500 kb genomic windows. Note that this is the same as Figure 3 but with axes units. Also note that the axes are different across rows and columns of the matrix.

**Figure S9.**
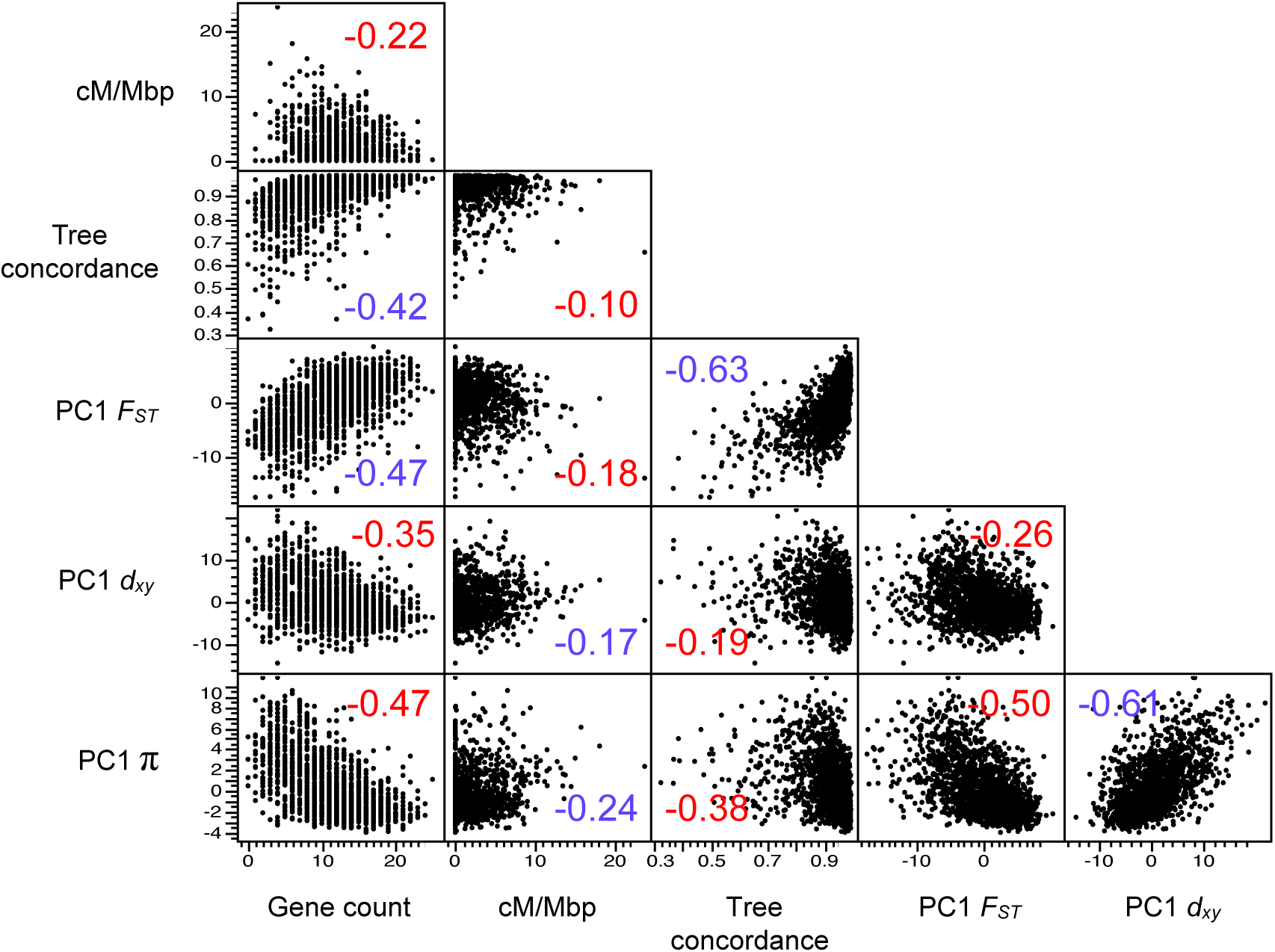
Bivariate plots among measures of variation and genomic features across 100 kb genomic windows. The number is the correlation coefficient. Positive correlation coefficients are colored blue and negative coefficients are colored red.

**Figure S10.**
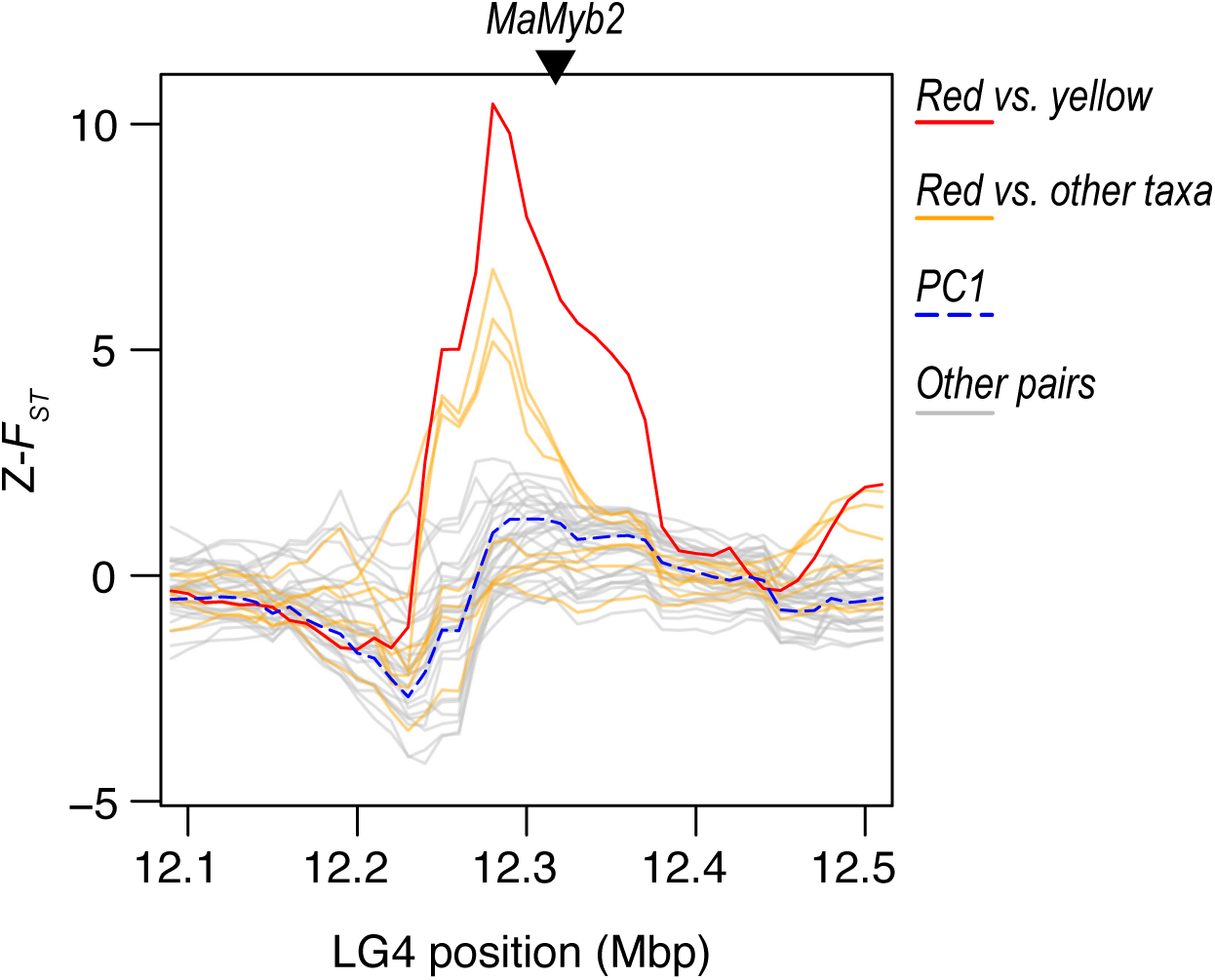
A large-effect adaptive locus shows a lineage-specific signature of positive selection. Plots of Z-transformed *F_ST_* across the genome, estimated in 100 kb sliding windows (step size 10 kb). The red line shows values between the red and yellow ecotypes of subspecies *puniceus*. The orange lines show other comparisons with the red ecotype of *punicues*, and the gray lines show the values of all other comparisons. The dashed blue line shows the first PC calculated across all of the comparisons. The triangle marks the position of the gene *MaMyb2*. A *cis-*regulatory mutation that is tightly linked to this gene is responsible for the shift from yellow to red flowers (Corresponds to Figure 4 in the main text).

**Figure S11.**
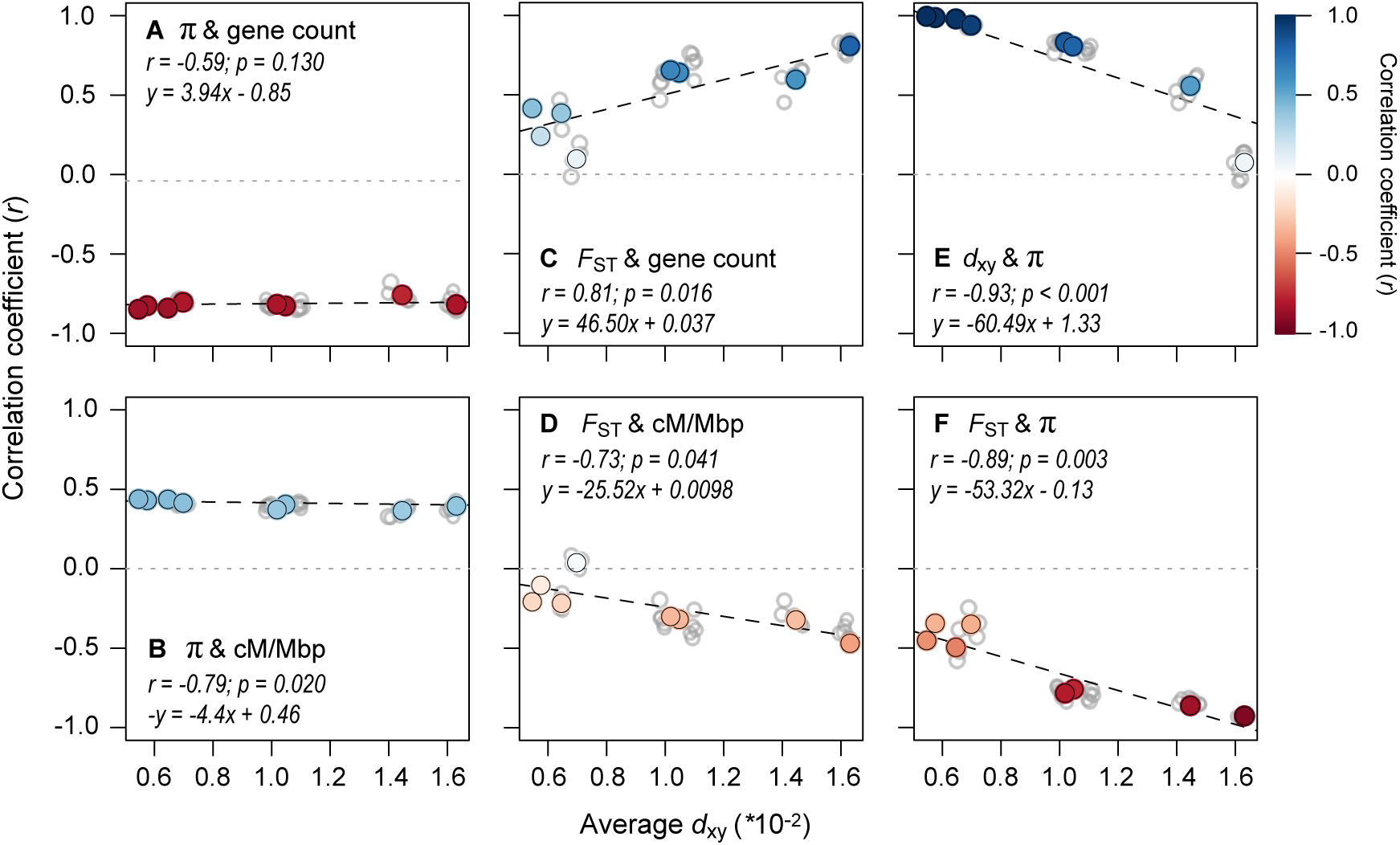
The range of divergence times reveals static and dynamic signatures of recurrent indirect selection. Correlations between variables (500 kb windows) for all 36 taxonomic comparisons (gray dots) plotted against the average *d*_xy_ as a measure of divergence time. The left panels show how the relationships between π (each window averaged across a pair of taxa) and (A) gene count and (B) recombination rate vary with increasing divergence time. The middle panels (C & D) show the same relationships, but with *F_ST_*. The right panels show the relationships between (E) *d_xy_* and π and (F) *F_ST_* and π. The regressions (dashed lines) in each plot are fitted to the eight independent contrasts (colored points) obtained using a phylogenetic correction, with the regression equation and strength of the correlation given in each panel. The color gradient shows the strength of the correlation.

**Figure S12.**
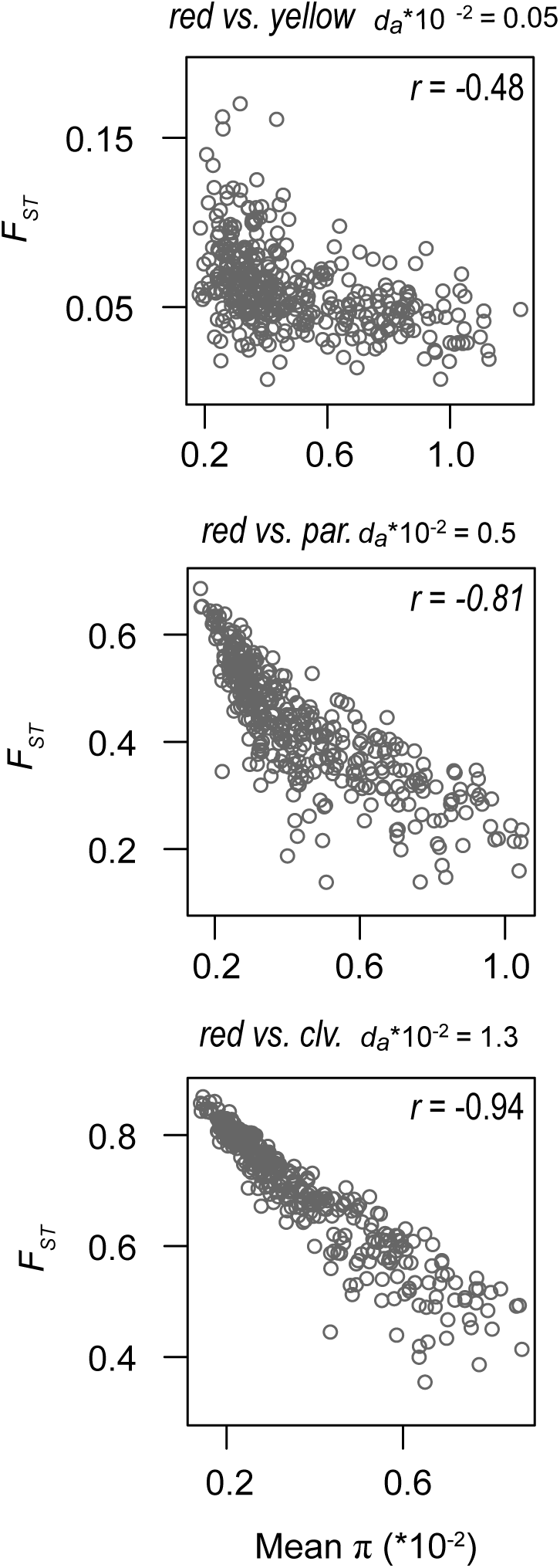
Negative correlation between nucleotide diversity and differentiation becomes stronger with increasing divergence time. Bivariate plots of the correlation between *F_ST_* and π at varying levels of sequence divergence (*d*_xy_).

**Figure S13.**
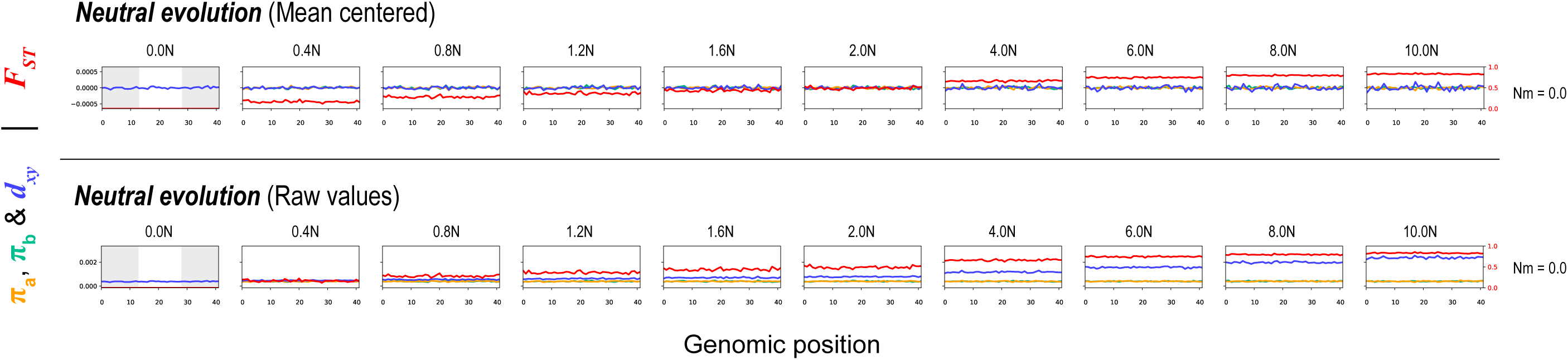
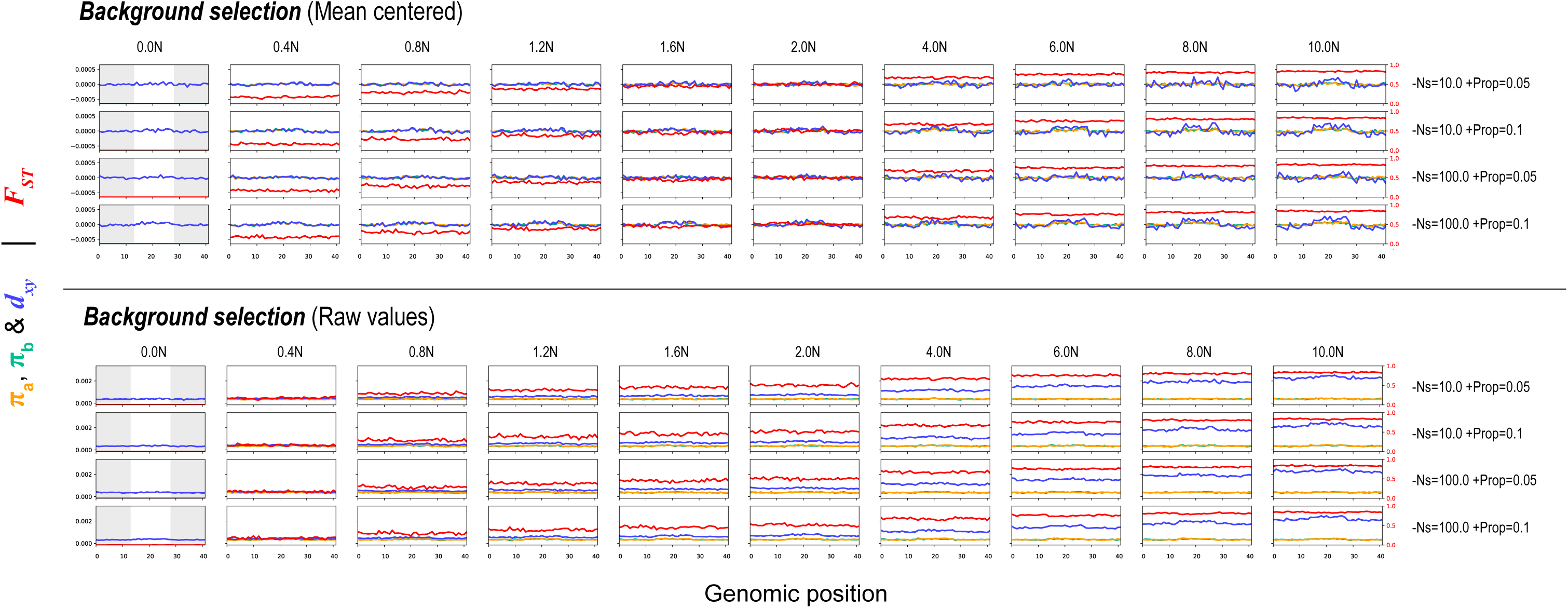
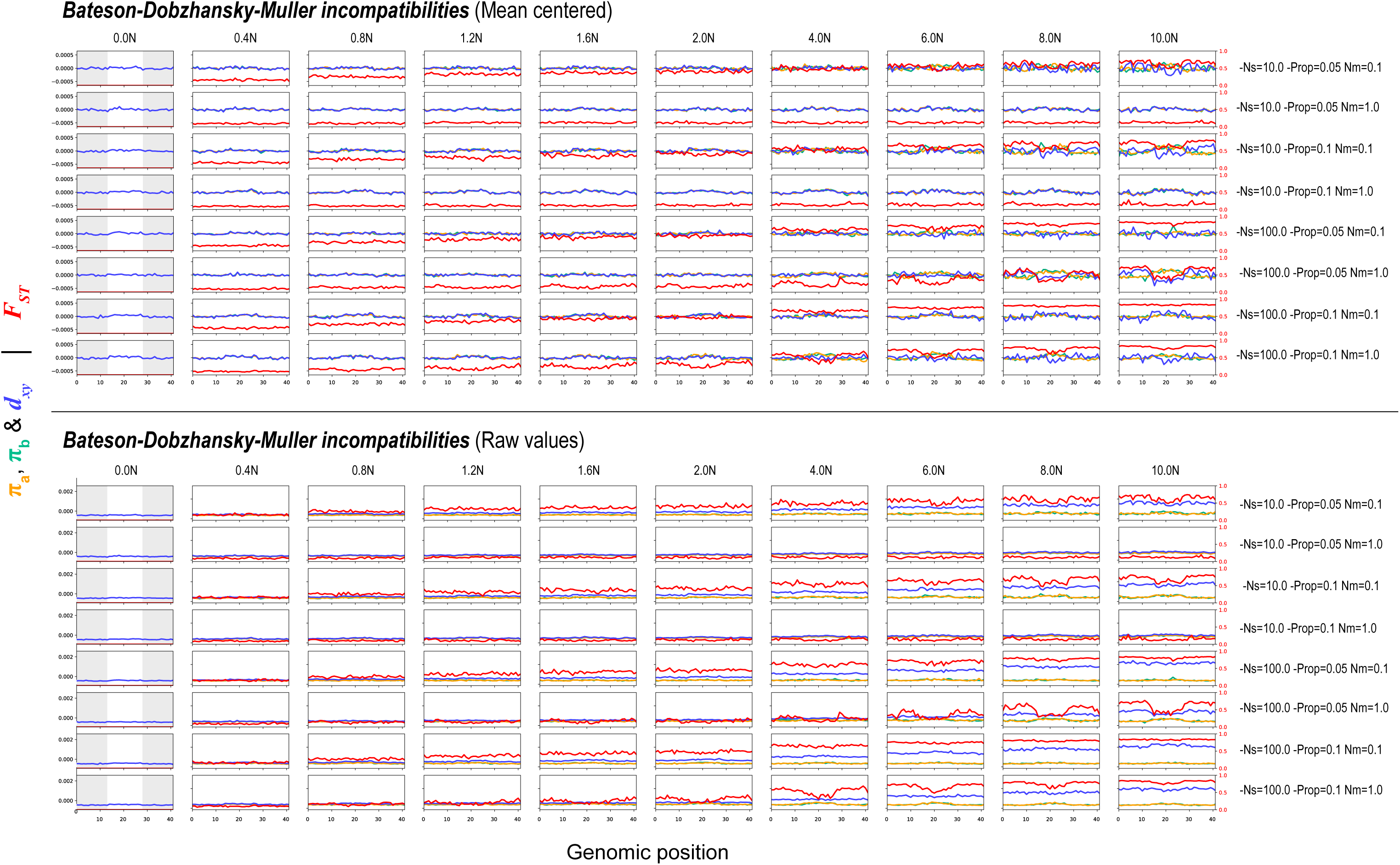
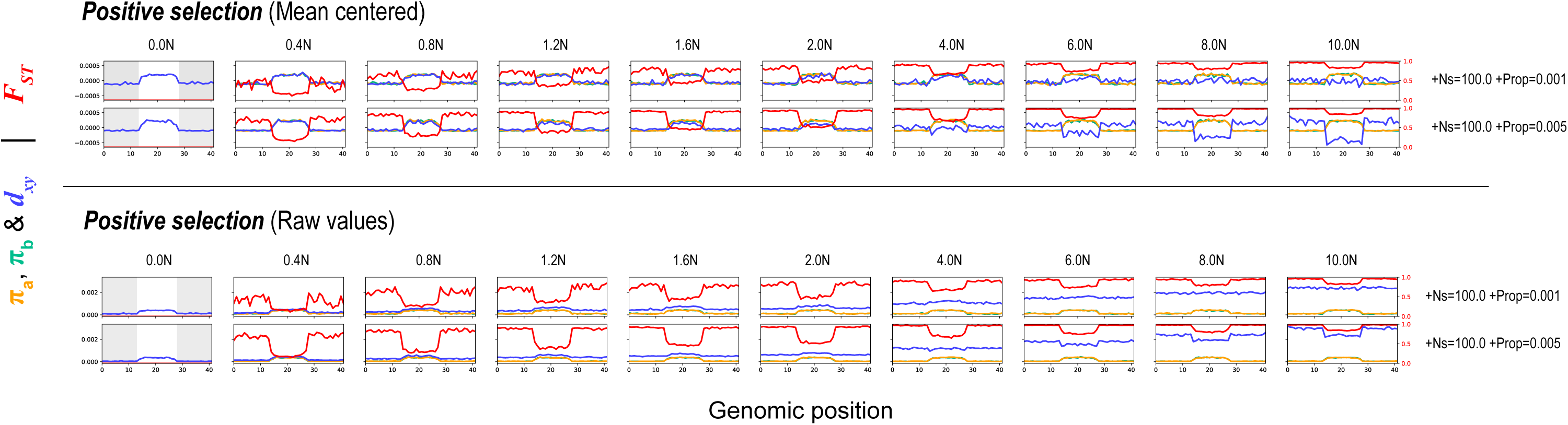
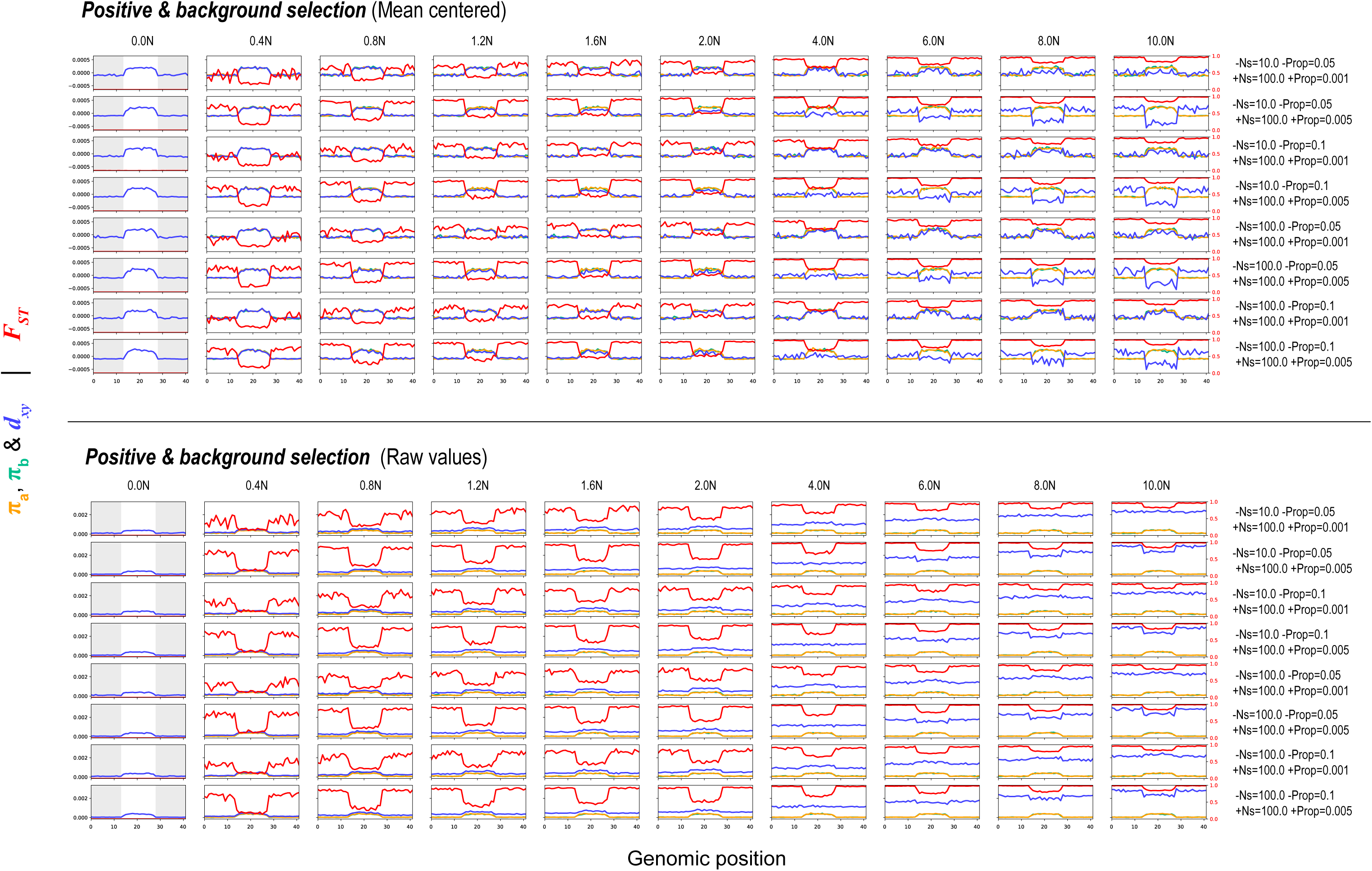
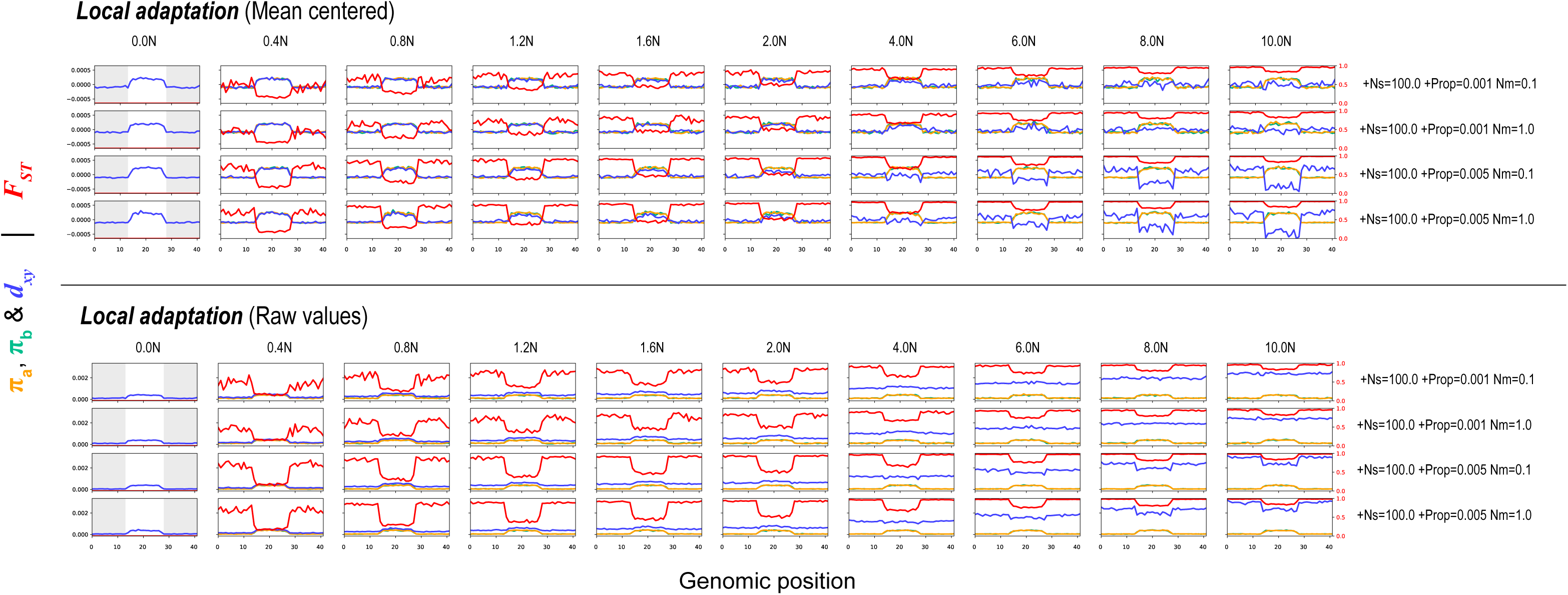
Genomic landscapes simulated under different different divergence histories. Each row of plots shows patterns of within- and between-population variation (π, *d_xy_*, and *F_ST_*) across a 21Mb chromosome (500 kb windows) at ten timepoints (in N generations, where N = 10,000) for one parameter combination of six scenarios: neutral divergence, background selection, Bateson-Dobzhansky-Muller incompatibilities, positive selection, background selection and positive selection, and local adaptation. The grey boxes in the first column show the areas of the chromosome that are constrained by selection. Mean centered (above line) and raw values (below line) of π, *d_xy_*. The parameter Ns modulates the average selective coefficient (where s = Ns/N) while Prop is the proportion of new mutations that are not neutral, Nm is the average number of migrants per generation.

**Figure S14.**
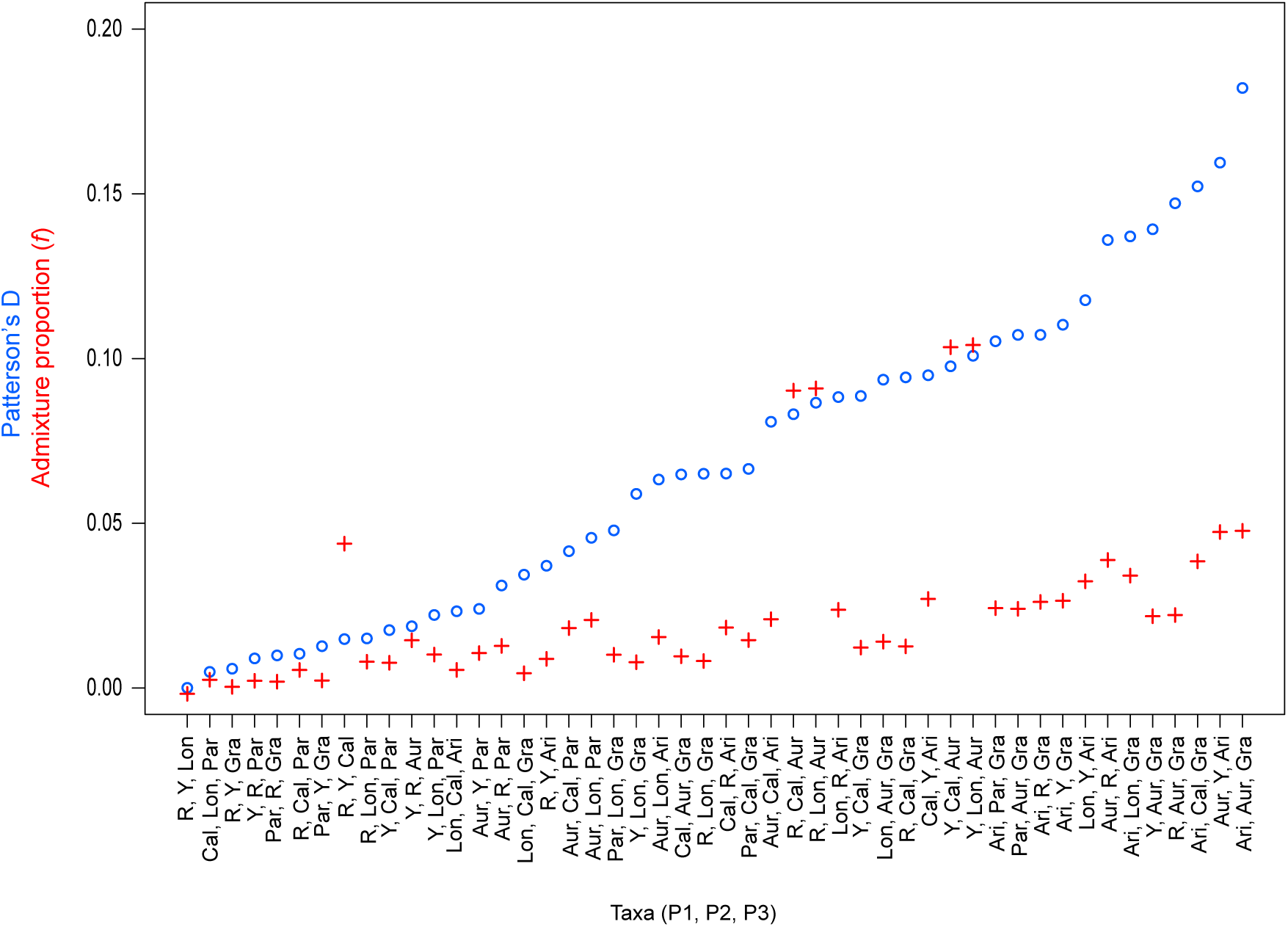
Evidence for widespread gene flow across the bush monkeyflower radiation. Genome-wide estimates of Patterson’s *D* (blue circles) and the admixture proportion (*f;* red crosses) are shown for all 48 possible four taxon comparisons, with *M. clevelandii* as the outgroup in each test. The tests are ordered by increasing values of *D*, and each value is significant based on a block jackknife approach. The taxa included in each test are shown in the order P1, P2, P3 and *D* is always positive, meaning gene flow occurs between P2 and P3.

**Figure S15.**
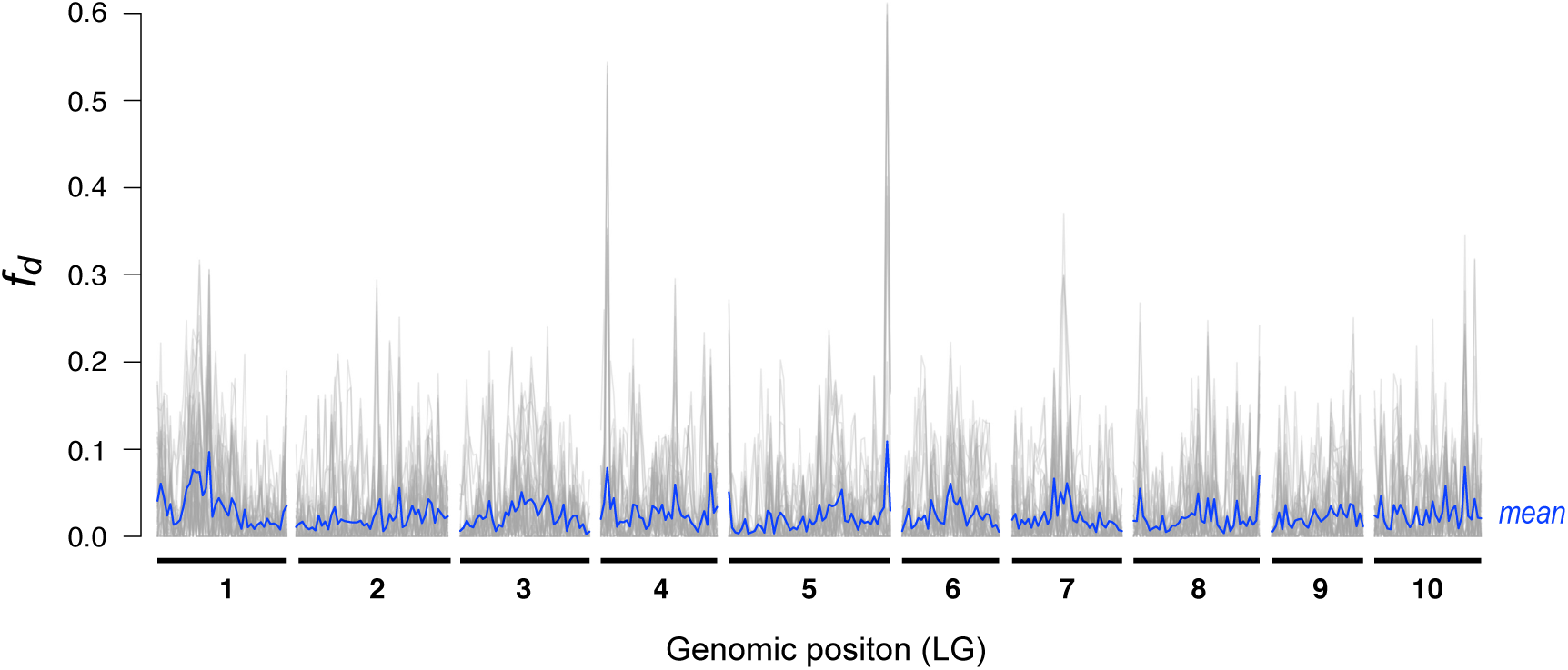
Variation in levels of admixture across the genome. The grey lines show measures of a modified test of the admixture proportion, *f_d_,* estimated in 500kb non-overlapping windows for 48 different four-taxon comparisons, plotted across the 10 linkage groups of the bush monkeyflower genome. The blue line gives the mean value of *f_d_*, calculated by taking the average value across all 48 tests in each genomic window.

## Notes

#### Summary of Updates

Taking into account reviewer comments and feedback from other people.

